# A platform for lab management, note-keeping and automation

**DOI:** 10.1101/2024.07.08.602487

**Authors:** Aubin Fleiss, Alexander S. Mishin, Karen S. Sarkisyan

## Abstract

We report a lab management concept and its no-code implementation based on general-purpose database services, such as Airtable. The solution we describe allows for integrated management of samples, lab procedures, experimental notes and data within a single browser-based application, and supports custom automations. We believe that this system can benefit a wide scientific audience by offering communication-less retrieval of information, collaborative editing, unified sample labelling and data keeping style. A template database is available at airtable.com/universe/expPcKlB7VCHE6wVK/lab-management.

## Main text

Research laboratories rely on dynamic registries of samples, experiments, and data. Keeping and updating them is essential for productive and coherent research. In addition to samples and data management, most researchers keep notes of their experimental work, typically in the form of physical lab notebooks or their electronic equivalents.

While life science research becomes increasingly industrialised (Chui et al. 2020), most laboratories and small companies do not have resources to adapt industrial-scale database solutions to their research pipelines, nor do they have the work throughput to justify their maintenance. Based on the survey of 233 life science labs we performed for this study, the most commonly used solution is a lab notebook, physical or electronic, combined with registries of samples as spreadsheets (**Supplementary Data 1**).

*Lab notebooks* are a free-form note-keeping solution. Users decide entirely on the structure of their records. While serving the purpose of documenting experiments, notebooks remain personal and disparate in format (**Figure 1A**). They are also text-based and linear, lacking the capability to explicitly store the relationships between objects in the research workflow. This hampers information retrieval and transmission, and may limit team productivity — an effect further worsened by the regular staff turnover in academic labs.

**Figure 1.**
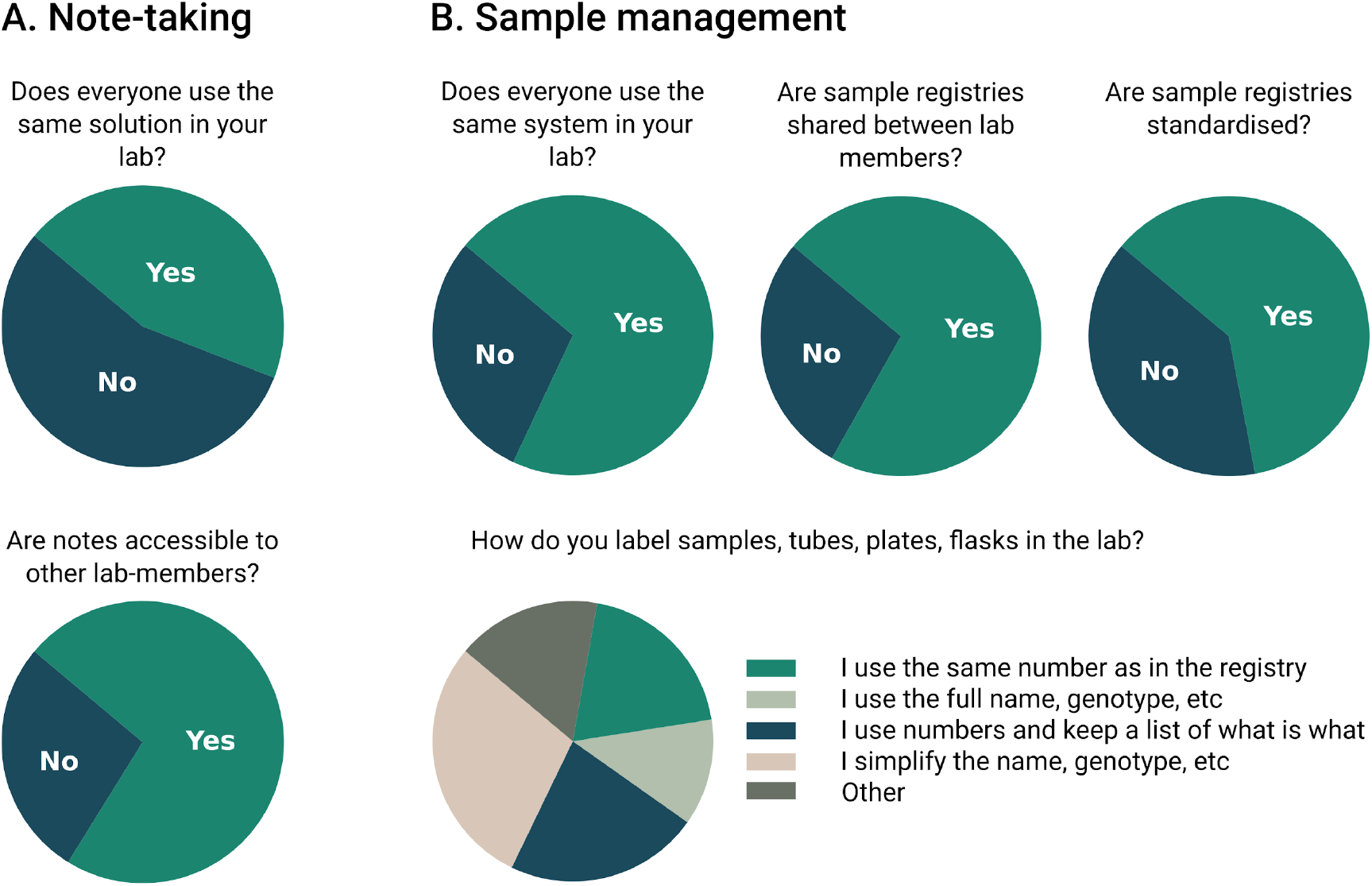
Selected survey results on sample management and note-taking practices. A. Lab-wide note-taking practices. B. Lab-wide sample-management practices.

*Spreadsheets* often complement notebooks as sample registries, providing a structured template to list research objects and their attributes. Spreadsheets do not store relationships between research objects, do not support different field types, such as file storage, and can accumulate hard-to-detect version conflicts if collaborative editing is not supported. They also do not provide means to engineer user behaviour through custom interfaces or enforceable formatting rules. In our experience, this results in “sporadic” data recording, with higher-complexity tasks often becoming under-documented due to oversights or users cutting corners on burdening bureaucracy.

Finally, *electronic lab notebooks* often integrate notekeeping with database-like registries. As databases allow storage of the relationship between objects of research workflows, such integrations simplify information retrieval, sharing, and referencing (Oleksik et al. 2014; Walsh and Cho 2013). These range from specialised Laboratory Information Management Systems (LIMS), such as Benchling Notebook, LabArchives, Labfolder, CERF Notebook or LabGuru, to general-purpose off-the-shelf software adapted for scientific use, such as Microsoft OneNote, Evernote, Notion (Wikipedia 2024; Eleven Therapeutics 2023). A number of open-source LIMS have also been developed, including OpenBIS (Barillari et al. 2016), LabInform ELN (Schröder and Biskup 2023), Leaf LIMS (Craig et al. 2017), Chemotion ELN (Tremouilhac et al. 2017) and eLabFTW (CARPi et al. 2017). However, we found that these solutions often had interfaces below modern industry standards, lacked essential features for life science laboratories (Myers et al. 2018) or required specialist support for deployment and maintenance. The apparent lack of an optimal solution rhymed with an observation from our survey that numerous labs employed note-taking or sample-management solutions that either did not support or only partially supported features they themselves have listed as essential (**Supplementary Data 1**).

While assessing existing approaches, we noticed that the *entire* research workflow of a lab is usually not reflected in the structure of solutions scientists use. We anticipated that integration of the entire research workflow within the lab management system could result in an “ecosystem” effect, creating a single source of truth (Wikipedia, 2024b) for all experiments performed in the lab, enabling *in-silico* automations, and improving productivity through reduction of human errors and communication burden.

To test this hypothesis, we conceptualised the workflow of a synthetic biology lab as a graph of interlinked objects of different types (**Supplementary Figure 1, Supplementary Video 1**). Some of the objects were *abstract*: for example, a Construct was an abstract object representing the expected DNA sequence of a plasmid. Others were *physical*, such as Molecules which are samples of purified DNA corresponding to that desired Construct.

Workflow objects often had Parent-Child relationship: several Molecules (Children) obtained from different bacterial clones were physical embodiments of a single expected Construct (Parent) (**Figure 2A**). Other examples of objects in the workflow included Glycerol stocks, PCR reactions, Golden Gate reactions, Transformations, Competent cells, People, Datasets, Protocols, etc (**Supplementary Figure 1**).

**Figure 2.**
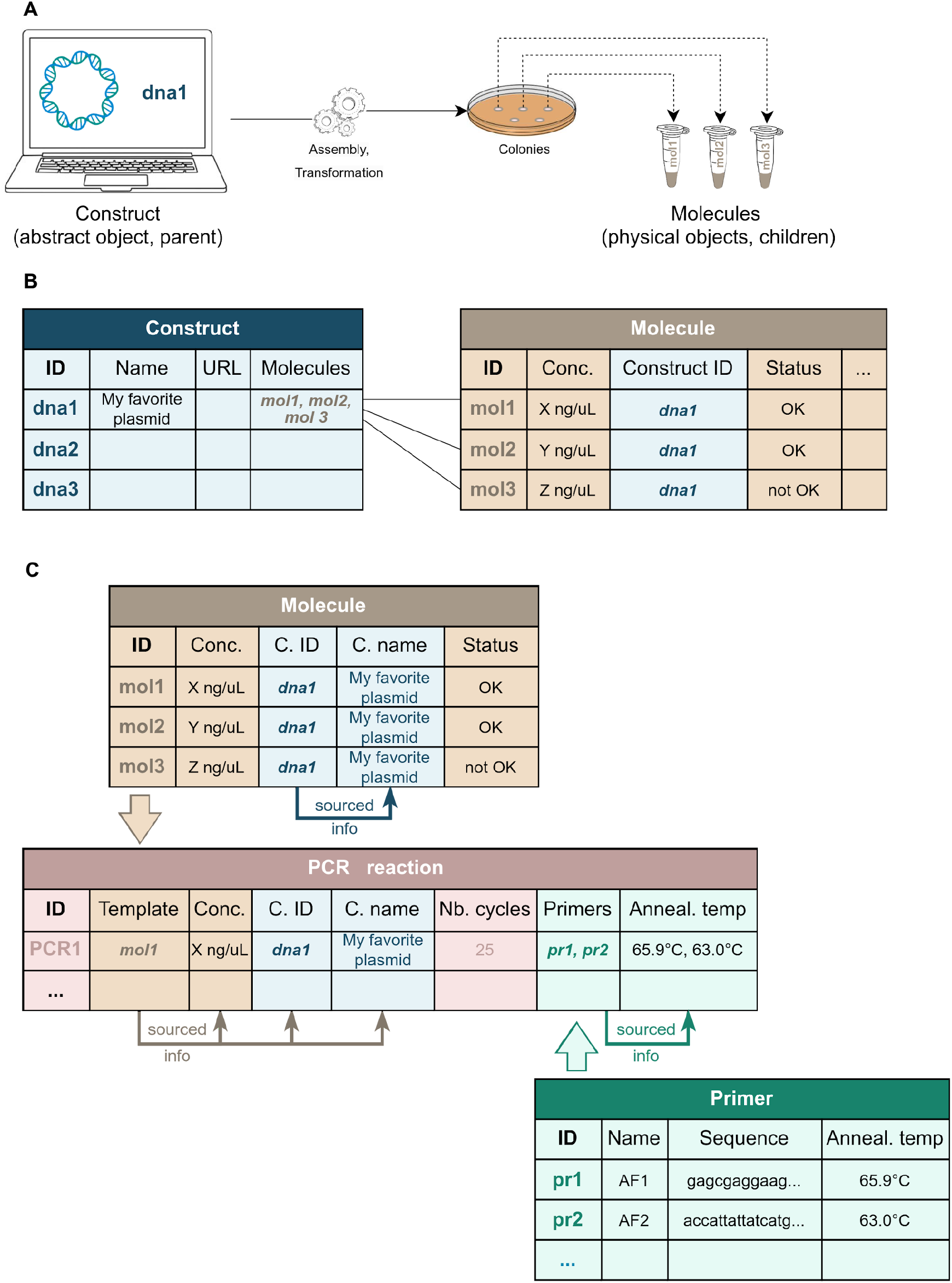
From workflow objects to table relationships and information propagation. A. Constructs are abstract parent objects with multiple physical children objects, which are the molecules. B. Transforming the concepts of “Construct” and “Molecule” into tables connected by a link field. The link field, containing IDs of records stored in a different table, is in italics. C. Building the PCR table with information propagation. Fields inherent to PCR are in pink. Other columns consist of links (shown in italics on the figure) and data sourced from other tables without duplication (shown with thin arrows). Column colour indicates the source table. In the table PCR, “C. name” comes from the table Molecule, which in turn sources this information from the table Construct.

We then structured the entire lab workflow as tables in a database, using the cloud-based platform Airtable (**Figure 2, Supplementary Figure 2, Supplementary Video 2**). Along with other similar platforms like APITable, Google Tables, Baserow, Seatable and others, Airtable provides a no-code interface to a spreadsheet-database hybrid. For each type of research objects, we created a table to store objects of that type, with links between tables reflecting object relationships in the workflow (**Figure 2B**). For example, a record in the “PCR reactions” table was linked to records in the “Primers” table and a record in the “Molecules” table that corresponded to the DNA template used in the reaction (**Figure 2C**).

In each table, columns provided pre-defined templates to describe objects. In contrast to spreadsheets, database columns could be configured to contain files, images, or longer formatted notes — or to lookup properties of linked objects. The latter is important, as it allows propagating information throughout tables of the database and displaying it in multiple places from a single source (**Figure 2C**). Thus, a change in a property of a parental object (for example, an update to the description of a Construct, **Figure 2B**) is immediately propagated to all linked objects and displayed in other tables, avoiding manual copying of the information and the accumulation of errors.

When implemented in the lab, this database-centric framework favoured atomisation of both research objects and information. Each workflow element was registered as a single record in its respective table, and assigned a unique tube-label-friendly ID. For instance, each time a plasmid was re-purified from a bacterial culture, it was registered under a new Molecule ID, while remaining linked to the same Construct. Similarly, we found it productive to apply atomisation to experimental data and other information (literature analyses, project designs, lab meeting presentations etc). Each new “atom of work” was registered in the Experiments table, along with links to the corresponding files stored in External Storage, metadata, protocols and other objects used. Atomising information allowed the team to refer to the past experiments with their Experiment IDs.

Finally, integration of the *entire* lab workflow within a single system allowed custom automations to source information from the database (**Supplementary Figure 3**). We created scripts to auto-check genetic designs, auto-generate plasmid or plate maps, or auto-process standard datasets as soon as they are registered. Any event in the database — a new entry, or update in a specific field — can be configured to trigger automation. For example, registering a new DNA assembly reaction may trigger a custom script verifying correctness of the assembly, using the virtual digestion and ligation of linked Construct objects, and creating the expected plasmid map. Automations reduce bureaucratic burden on users, spot human errors preventing costly reworks and delays, and improve communication within the team. Some of our custom automation scripts are available from our Github repository (https://github.com/sarkisyan-lab/lab-management).

We believe that the implementation described here allowed integrated management of samples, lab procedures, notekeeping and experimental data without undue bureaucratisation for the user, or support burden. Notably, the database allows the content to be accessed, sliced and visualised in different ways by different people, without mutual interference or modification of the underlying data. In the **Supplementary Note 1**, we provide several additional empirical rules that help promote unified lab practices and ensure that the database remains relevant over time.

Having tested the system for over five years in two academic labs and two startup companies, we believe that this solution does indeed boost team’s productivity, supports very detailed searchable records of lab work, and makes traditional lab notebooks largely unnecessary. The system can be adopted gradually, and evolves alongside the workflow to always provide the necessary support and flexibility. It scales well with the growth of the organisation, and allows for effective asynchronous communication. Quality information can be obtained even when collaborators are absent, and less technical interaction is needed between team members, lowering communication burden. In our experience, the benefits of implementation of this framework are bigger for larger teams.

## Acknowledgements

The Synthetic Biology Group is funded by the MRC Laboratory of Medical Sciences (UKRI MC-A658-5QEA0). Development of some of the automated scripts was funded by RSF, project number 22-14-00400 (https://rscf.ru/project/22-14-00400/).

## Code availability

A template database is available at airtable.com/universe/expPcKlB7VCHE6wVK. Automation scripts are available at github.com/sarkisyan-lab/lab-management.

## Supplementary Information

**Supplementary Figure 1.**
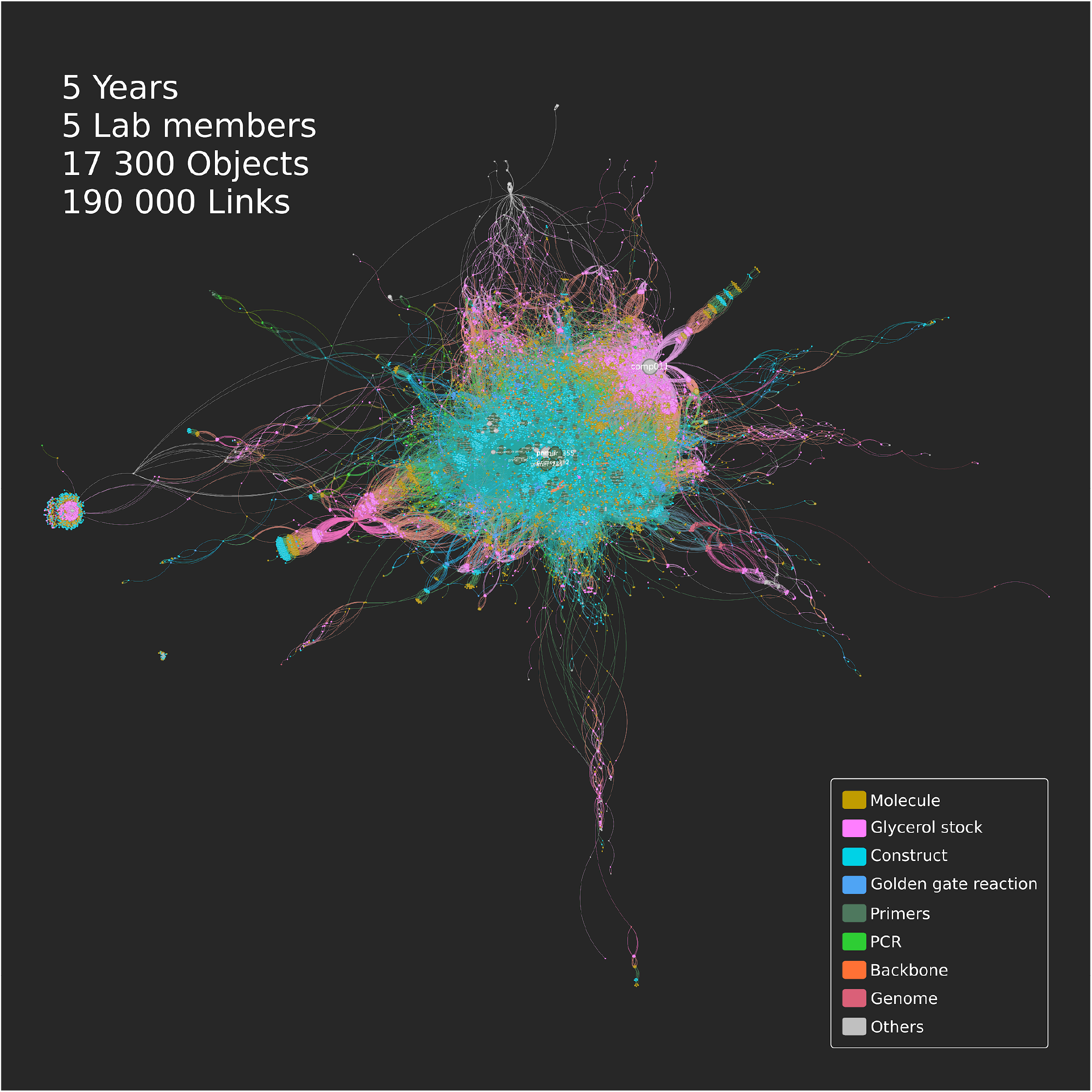
Graph showing all research objects registered in our laboratory by the five lab members over the past five years. Each object is represented as a node, with connections indicating their relationships. Nodes are coloured by object type, and their size reflects the number of connections. The ID of each object is displayed in white.

**Supplementary Figure 2.**
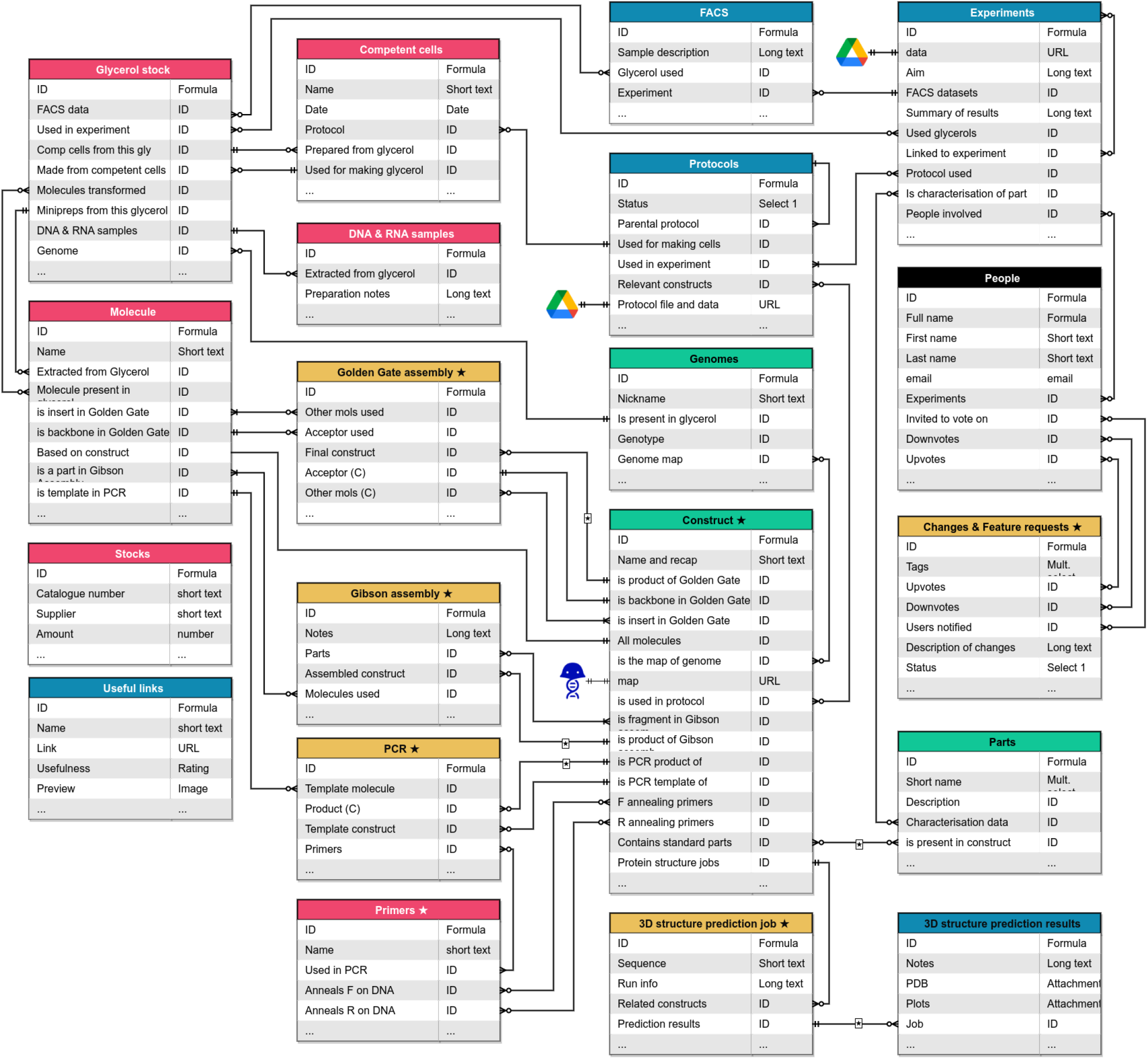
Database schema of Sarkisyan lab. Tables are shown as lists of column names and their respective data type. Colours indicate types of objects: red for physical assets, blue for datasets and documents, green for ideal objects, yellow for routine tasks, black for people. Stars ★ indicate where automation is involved in record validation, linking, or rendering visualisations. Cloud services are represented by their logo.

**Supplementary Figure 3.**
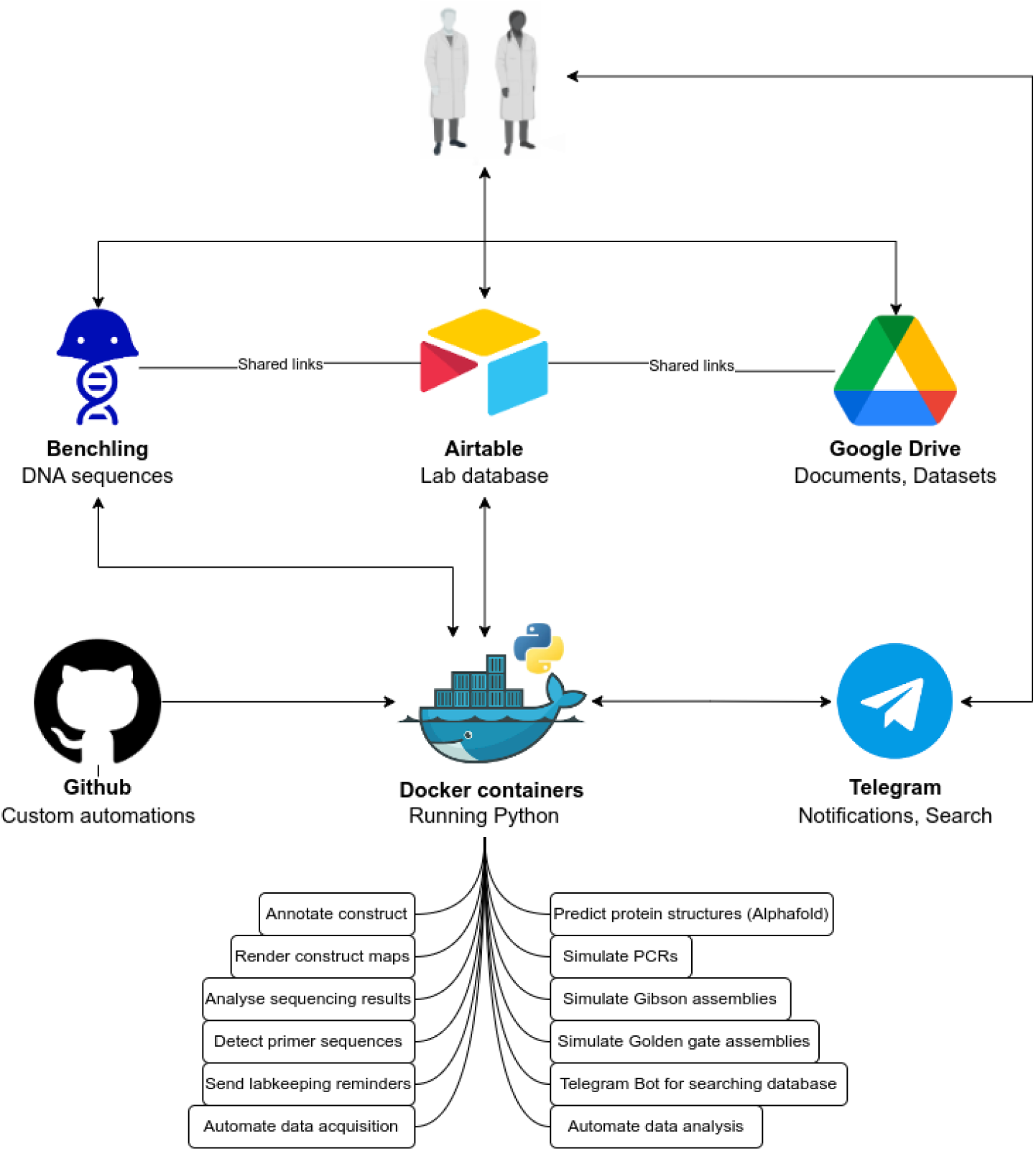
Cloud architecture of Sarkisyan lab management system. Users register experiments and biological samples in the database. Sequence maps stored on Benchling and free-form documents stored on Microsoft OneDrive or Google Drive are registered in the database via shared links. Some tables serve as an interface to submit computational jobs to Docker containers running custom Python scripts, deployed from a Github repository. A Telegram bot can be used to query the database interactively or relay notifications.

## Supplementary Note 1. Tips and guidelines

1. Single source of truth: every piece of information should only be stored in a single location in the database. If it needs to be shown in other places in the database, display it rather than copy it.
2. Set appropriate data types for all fields. This enforces unified format and helps make automations possible. For example, tags and categories should be picked from a list of predefined values to avoid variations in spelling.
3. Atomise object description and break information of different kinds into specialised fields.
4. Use concise names for tables and fields. Synchronise understanding of the database between the team members by adding descriptions to tables and fields.
5. Use concise tube-label-friendly names for records. Generate IDs automatically where possible. We recommend concatenating a human-readable prefix with a unique number (for example, mol713 or data2046).
6. Use record IDs to label tubes and other physical objects in the lab. Do not include other details. Handwritten details may get unreadable and become a source of error. Where automation is used, we recommend combining human-readable (text) and machine-readable information (barcodes) on the label. We recommend labelling tubes both on the lid and on the side.
7. By using consecutive numbering for objects and storing them in freezers, drawers and standardised boxes labelled with the first and last IDs they contain, the organisation of the lab becomes self-explanatory and it becomes unnecessary to register the location of an object in the database. We recommend sharing numbering between people working in the lab, and storing plasmids, glycerol stocks and other materials at a shared location.

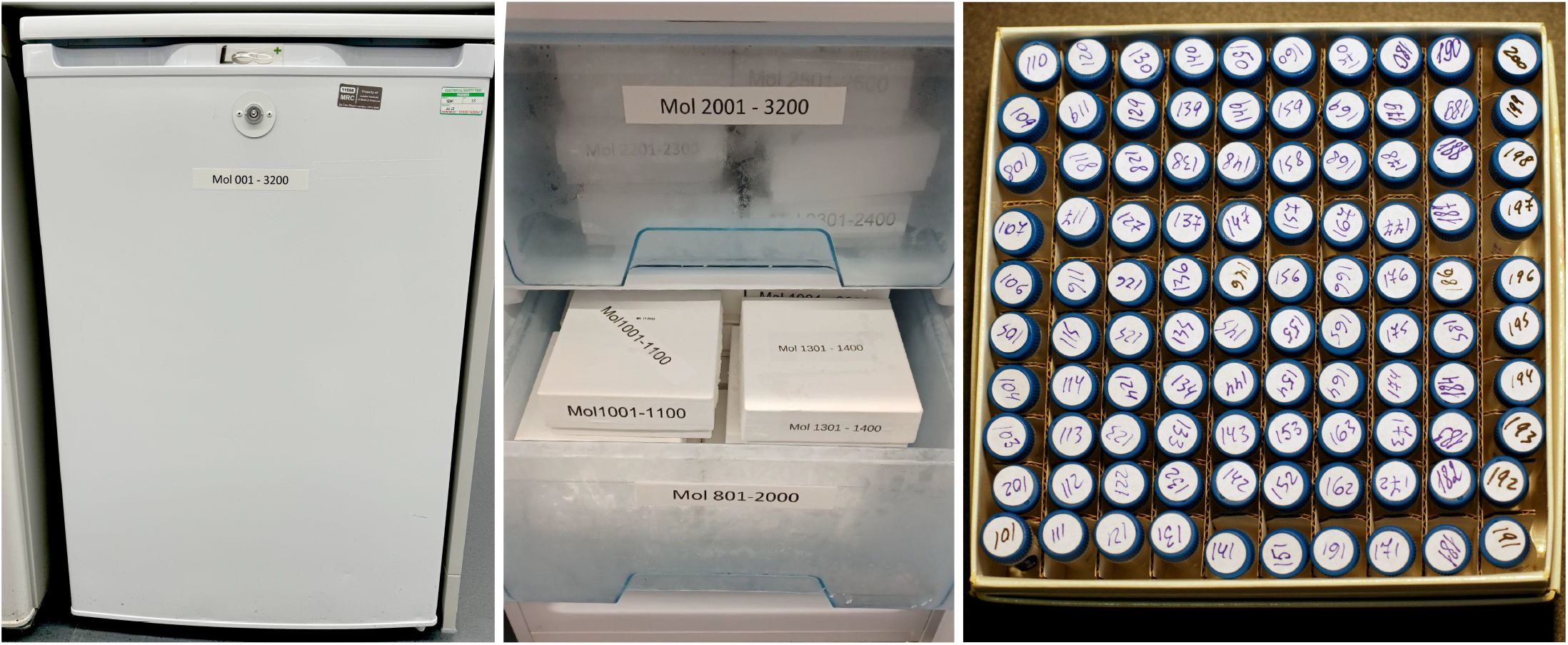
8. Use tags to label objects and facilitate database search. Tag search becomes more efficient than chronological search only a few weeks after an object is registered (Oleksik et al. 2014).
9. Keep the team synced using automated emails and notifications to lower communication efforts. For example, when a new experiment is registered in the database, people working on the project may be notified.
10. Some files may be attached and previewed in the database, but large file storage, file editing, access control and versioning should be managed on dedicated storage platforms. In our implementation, the database mostly stores metadata, while data and experimental notes are stored on Microsoft OneDrive and Google Drive, DNA sequences are stored on Benchling, code on Github, each linked to its corresponding entry in the database.

