## Supplementary Information 1 for "A platform for lab management, note-keeping and automation"

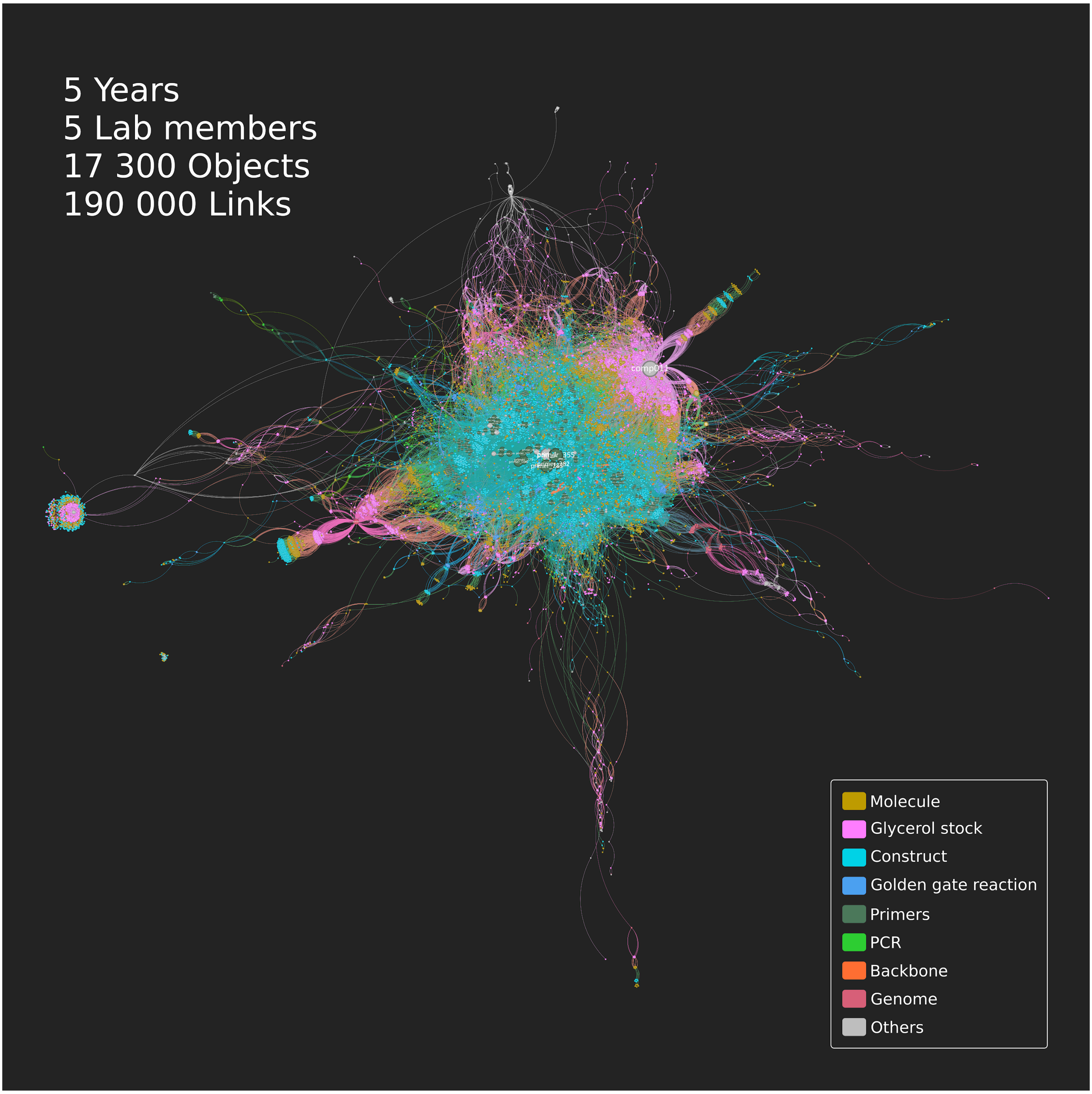


Supplementary Figure 1. Graph showing all research objects registered in our laboratory by the five lab members over the past five years. Each object is represented as a node, with connections indicating their relationships. Nodes are coloured by object type, and their size reflects the number of connections. The ID of each object is displayed in white.

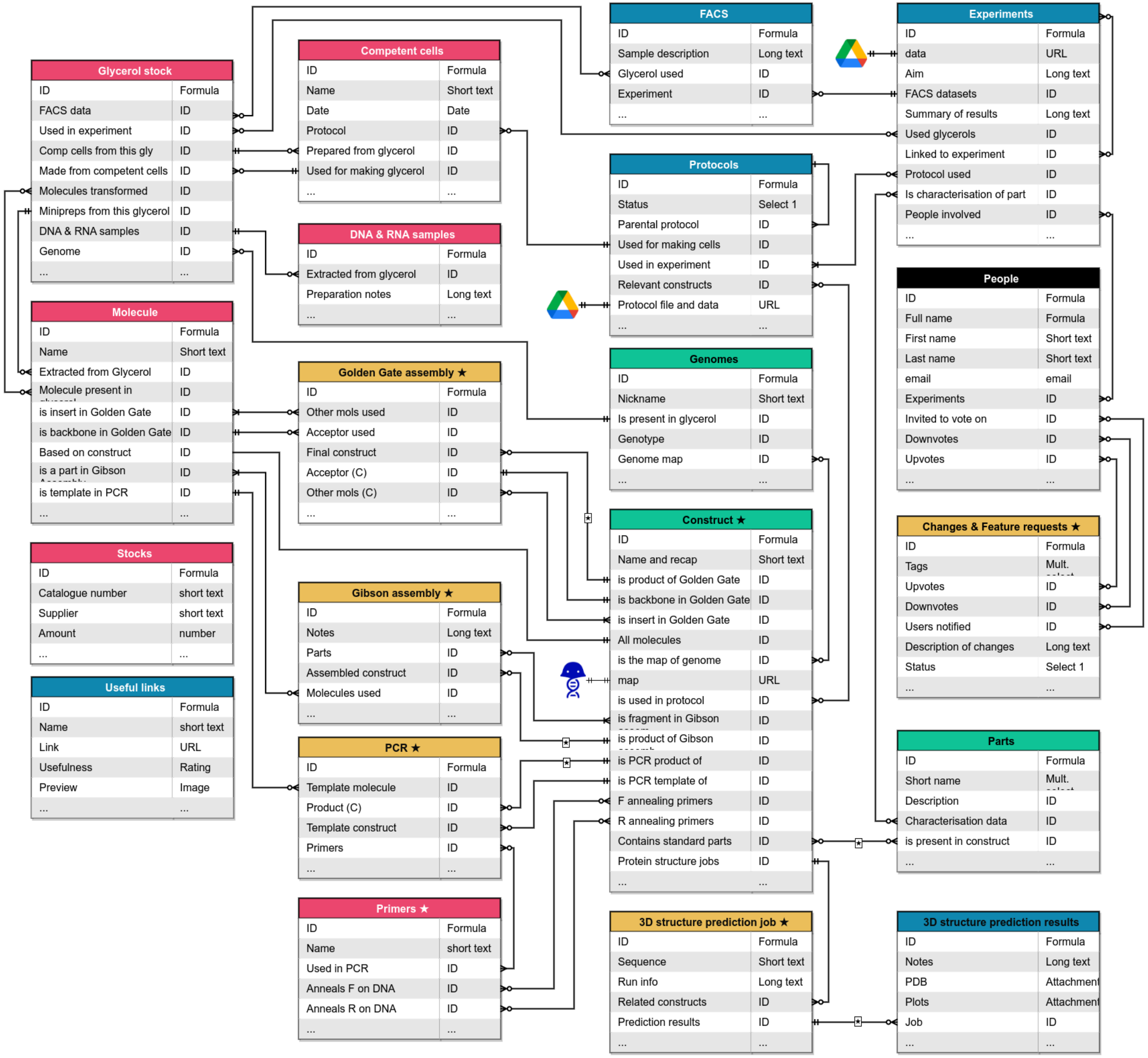


**Supplementary Figure 2.**
**Database schema of Sarkisyan lab.**
Tables are shown as lists of column names and their respective data type. Colours indicate types of objects: red for physical assets, blue for datasets and documents, green for ideal objects, yellow for routine tasks, black for people. Stars ★ indicate where automation is involved in record validation, linking, or rendering visualisations. Cloud services are represented by their logo.

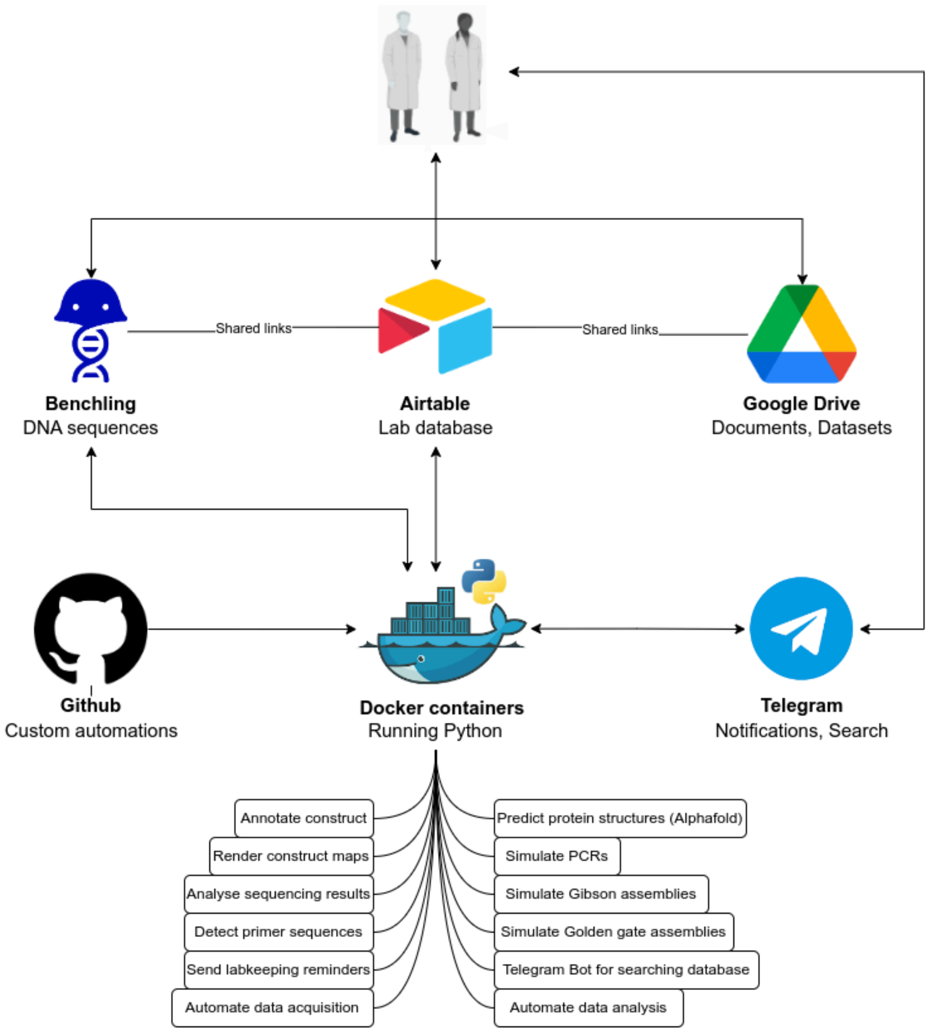


Supplementary Figure 3. Cloud architecture of Sarkisyan lab management system.
Users register experiments and biological samples in the database. Sequence maps stored on Benchling and free-form documents stored on Microsoft OneDrive or Google Drive are registered in the database via shared links. Some tables serve as an interface to submit computational jobs to Docker containers running custom Python scripts, deployed from a Github repository. A Telegram bot can be used to query the database interactively or relay notifications.

### Supplementary Note 1. Tips and guidelines

1. Single source of truth: every piece of information should only be stored in a single location in the database. If it needs to be shown in other places in the database, display it rather than copy it.

2. Set appropriate data types for all fields. This enforces unified format and helps make automations possible. For example, tags and categories should be picked from a list of predefined values to avoid variations in spelling.

3. Atomise object description and break information of different kinds into specialised fields.

4. Use concise names for tables and fields. Synchronise understanding of the database between the team members by adding descriptions to tables and fields.

5. Use concise tube-label-friendly names for records. Generate IDs automatically where possible. We recommend concatenating a human-readable prefix with a unique number (for example, mol713 or data2046).

6. Use record IDs to label tubes and other physical objects in the lab. Do not include other details. Handwritten details may get unreadable and become a source of error. Where automation is used, we recommend combining human-readable (text) and machine-readable information (barcodes) on the label. We recommend labelling tubes both on the lid and on the side.

7. By using consecutive numbering for objects and storing them in freezers, drawers and standardised boxes labelled with the first and last IDs they contain, the organisation of the lab becomes self-explanatory and it becomes unnecessary to register the location of an object in the database. We recommend sharing numbering between people working in the lab, and storing plasmids, glycerol stocks and other materials at a shared location.

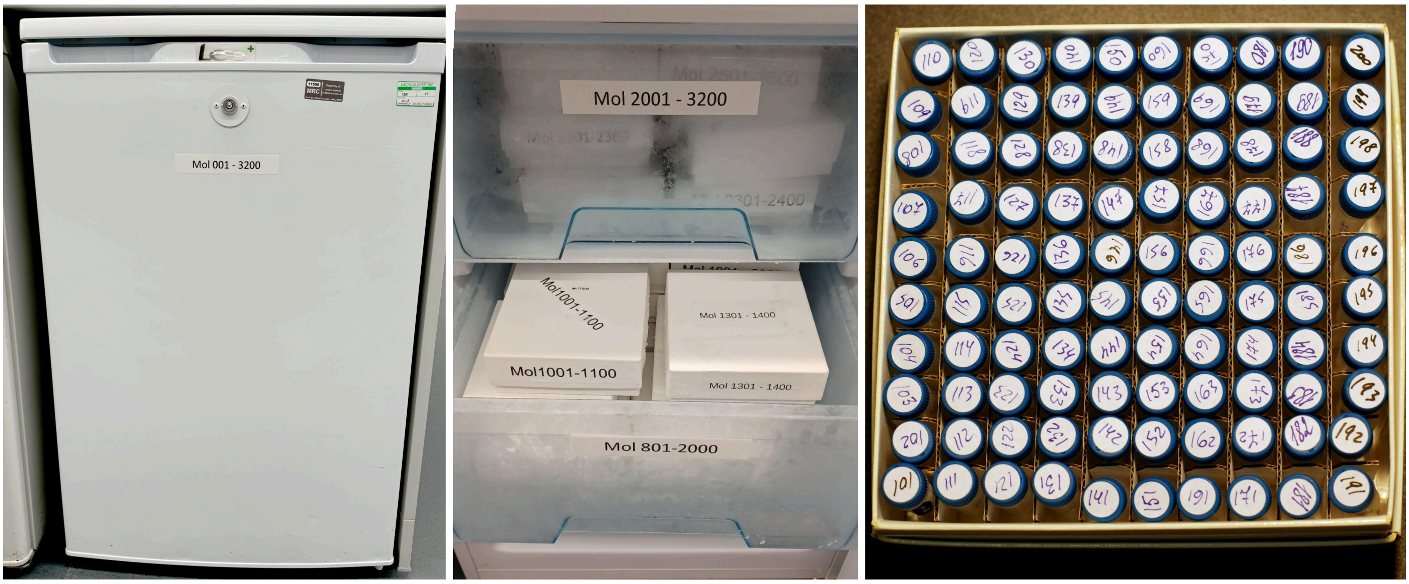


8. Use tags to label objects and facilitate database search. Tag search becomes more efficient than chronological search only a few weeks after an object is registered (Oleksik et al. 2014).

9. Keep the team synced using automated emails and notifications to lower communication efforts. For example, when a new experiment is registered in the database, people working on the project may be notified.

10. Some files may be attached and previewed in the database, but large file storage, file editing, access control and versioning should be managed on dedicated storage platforms. In our implementation, the database mostly stores metadata, while data and experimental notes are stored on Microsoft OneDrive and Google Drive, DNA sequences are stored on Benchling, code on Github, each linked to its corresponding entry in the database.
