## Supplementary Data 1 for "A platform for lab management, note-keeping and automation": Supplementary Data 1 - Note-taking.pdf

Benchling notebook

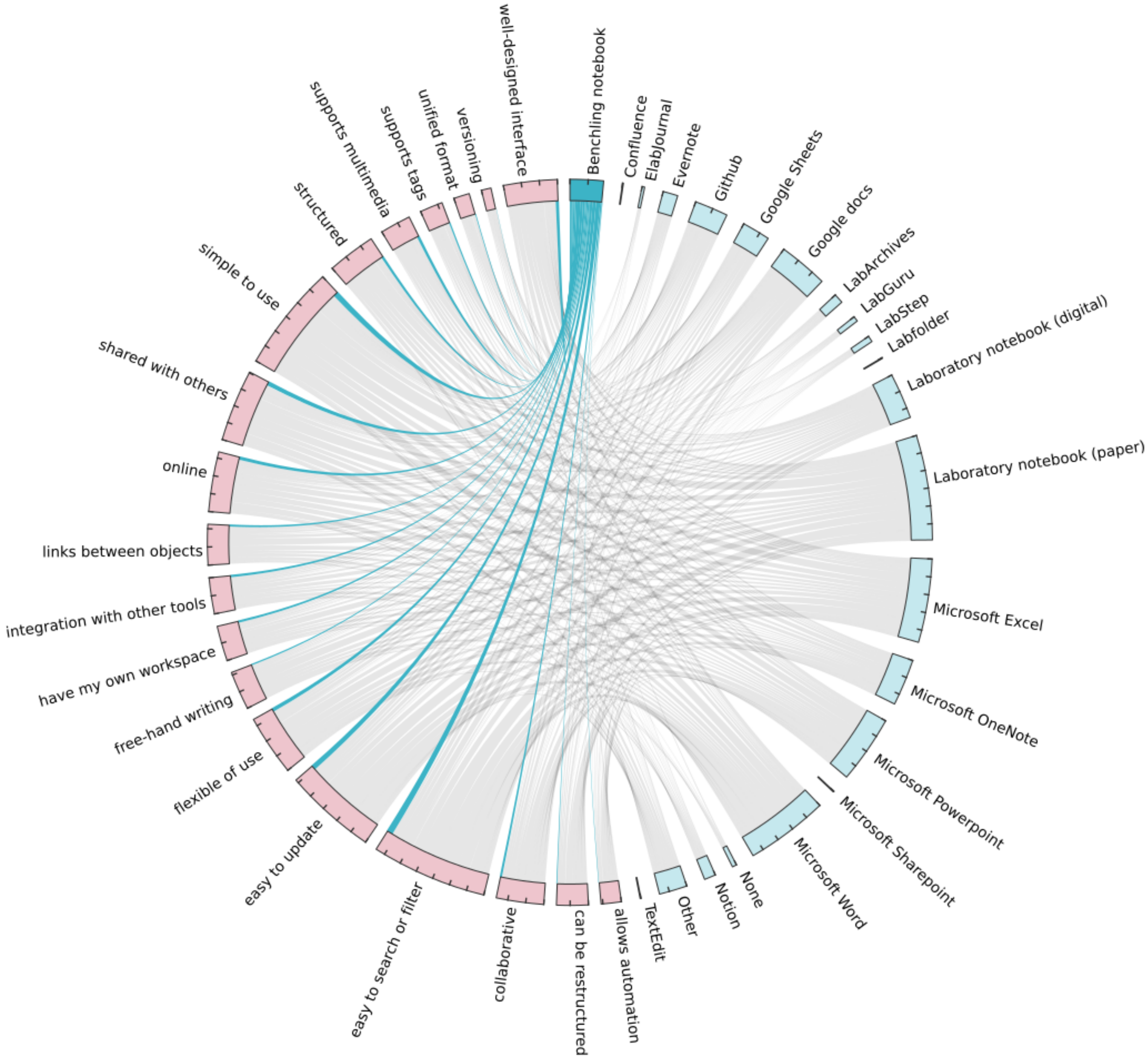


Note-taking solutions (blue arcs) vs desired features (red arcs). Arc length depicts the number of respondents, with ticks denoting 20 respondents. Link thickness shows the fraction of respondents using a solution and valuing a criterion. The chords connected to the arc 'Benchling notebook' are highlighted in color.

Confluence

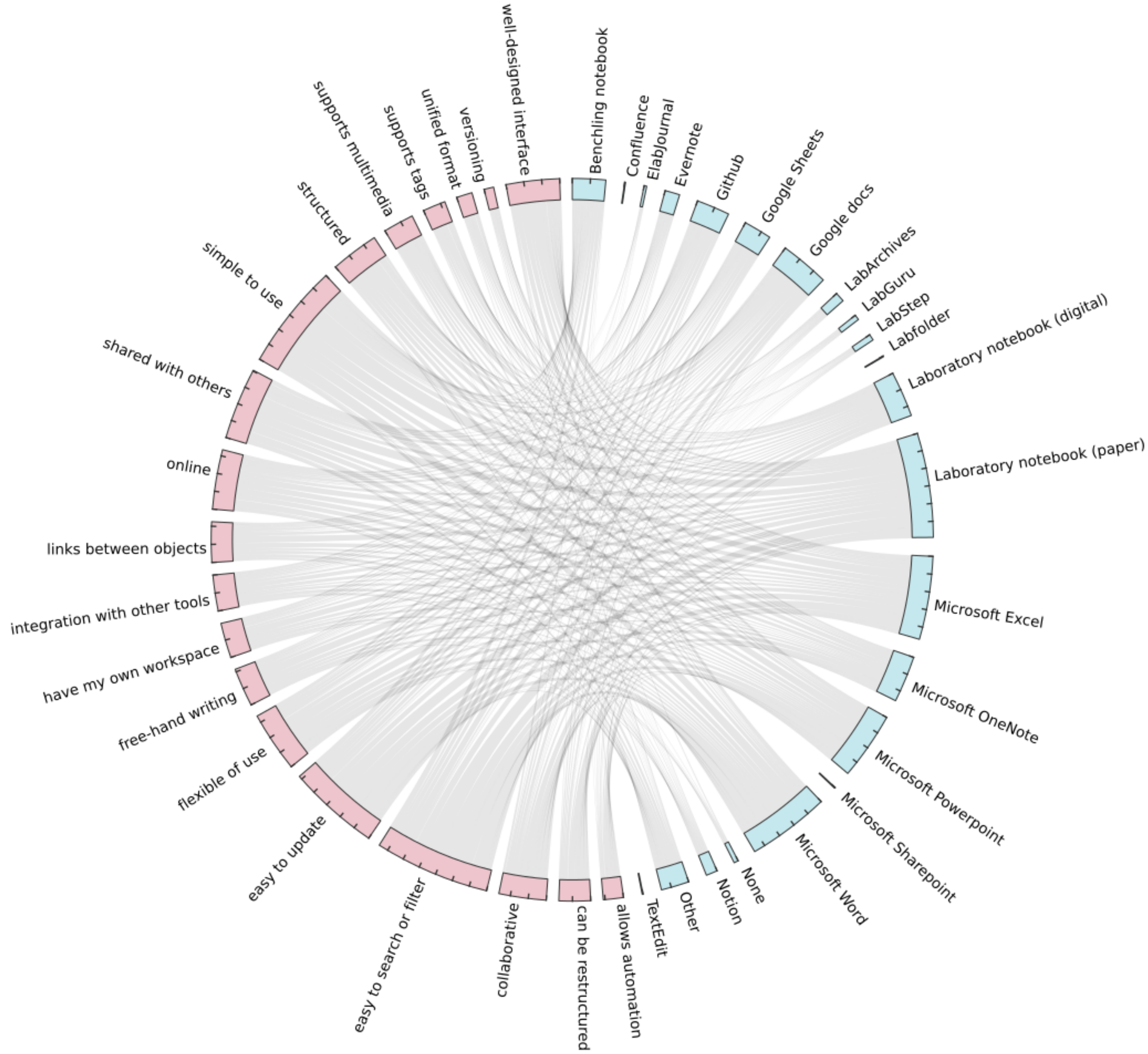


Note-taking solutions (blue arcs) vs desired features (red arcs). Arc length depicts the number of respondents, with ticks denoting 20 respondents. Link thickness shows the fraction of respondents using a solution and valuing a criterion. The chords connected to the arc 'Confluence' are highlighted in color.

ElabJournal

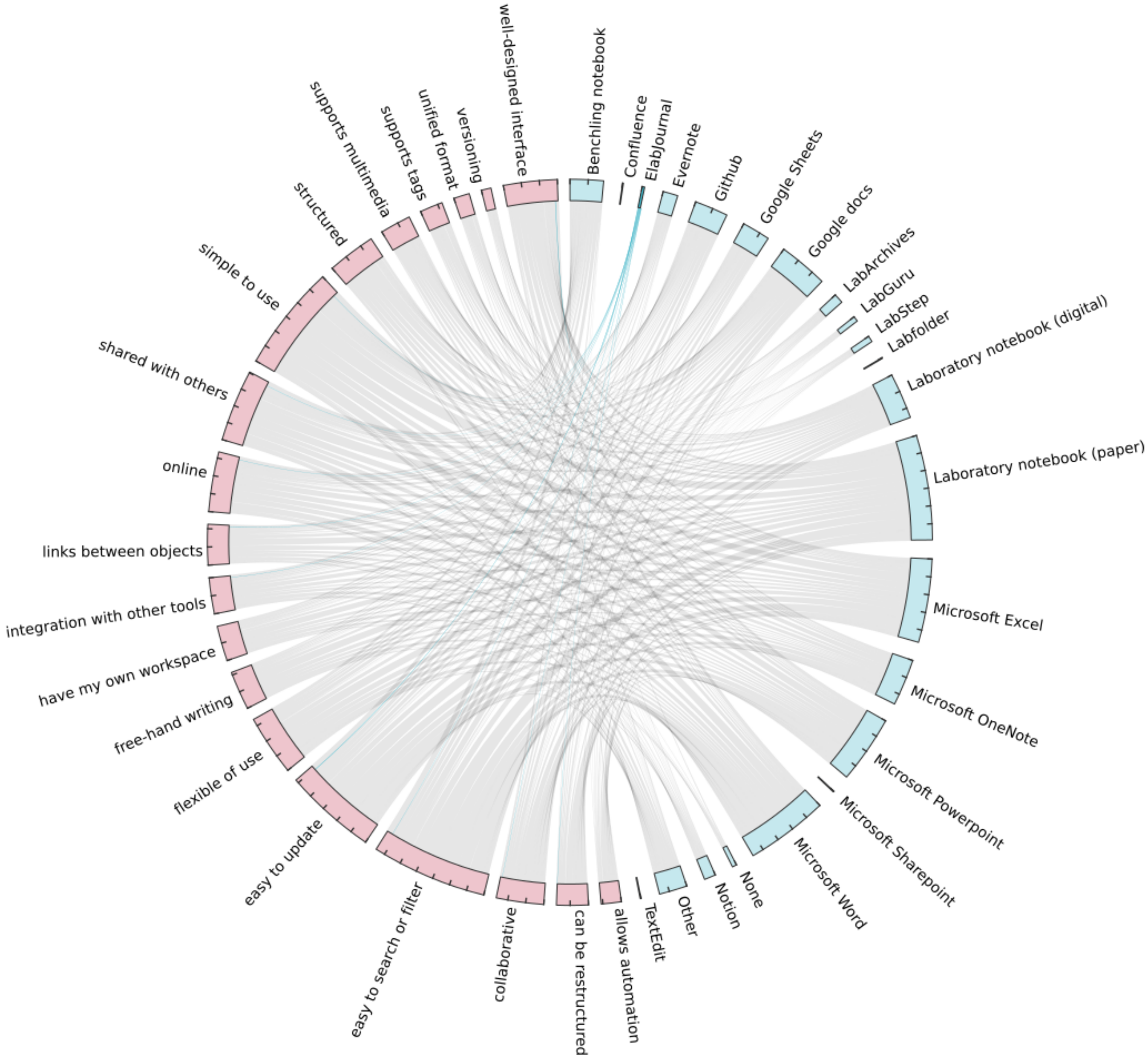


Note-taking solutions (blue arcs) vs desired features (red arcs). Arc length depicts the number of respondents, with ticks denoting 20 respondents. Link thickness shows the fraction of respondents using a solution and valuing a criterion. The chords connected to the arc 'ElabJournal' are highlighted in color.

Evernote

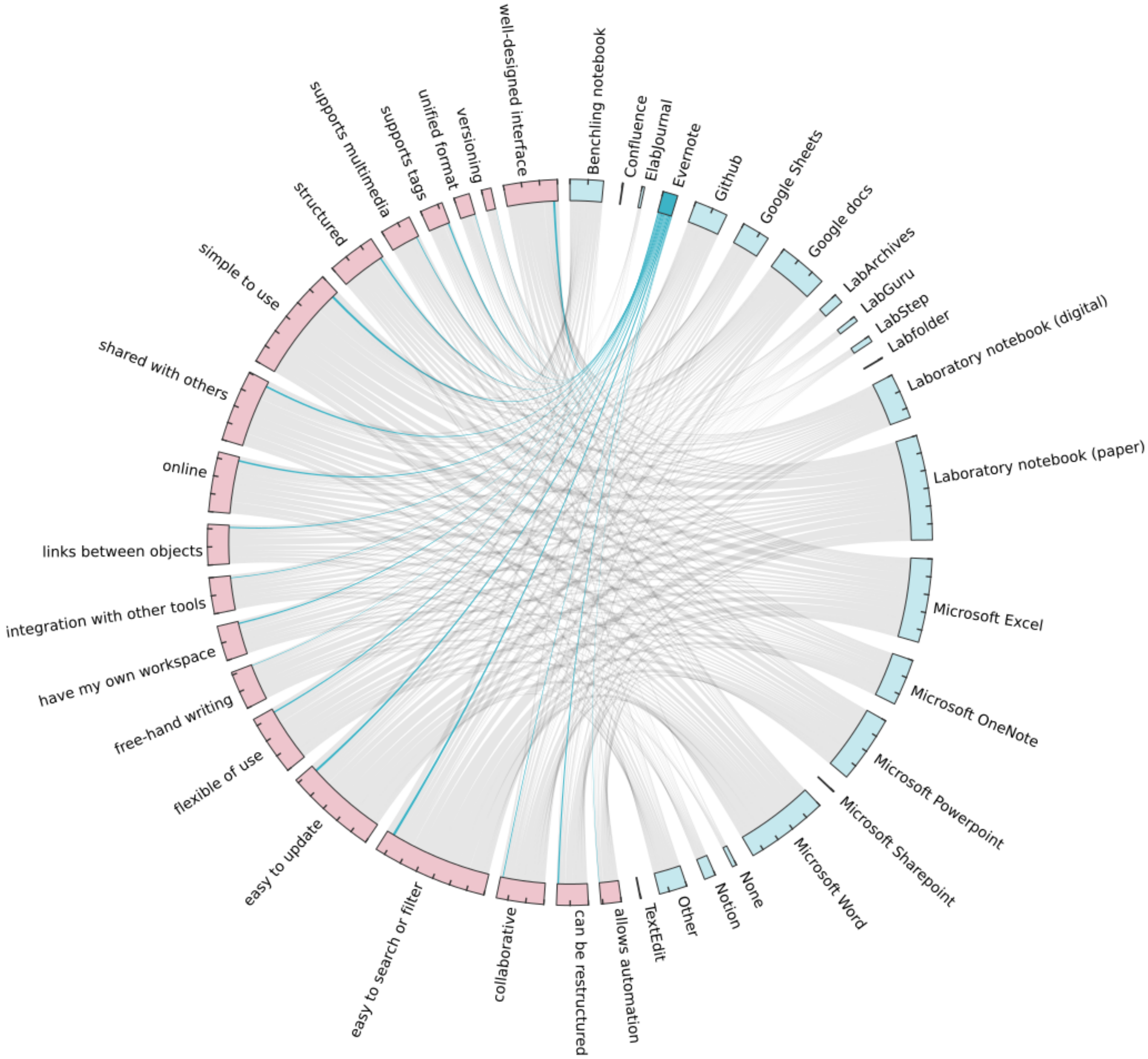


Note-taking solutions (blue arcs) vs desired features (red arcs). Arc length depicts the number of respondents, with ticks denoting 20 respondents. Link thickness shows the fraction of respondents using a solution and valuing a criterion. The chords connected to the arc 'Evernote' are highlighted in color.

Github

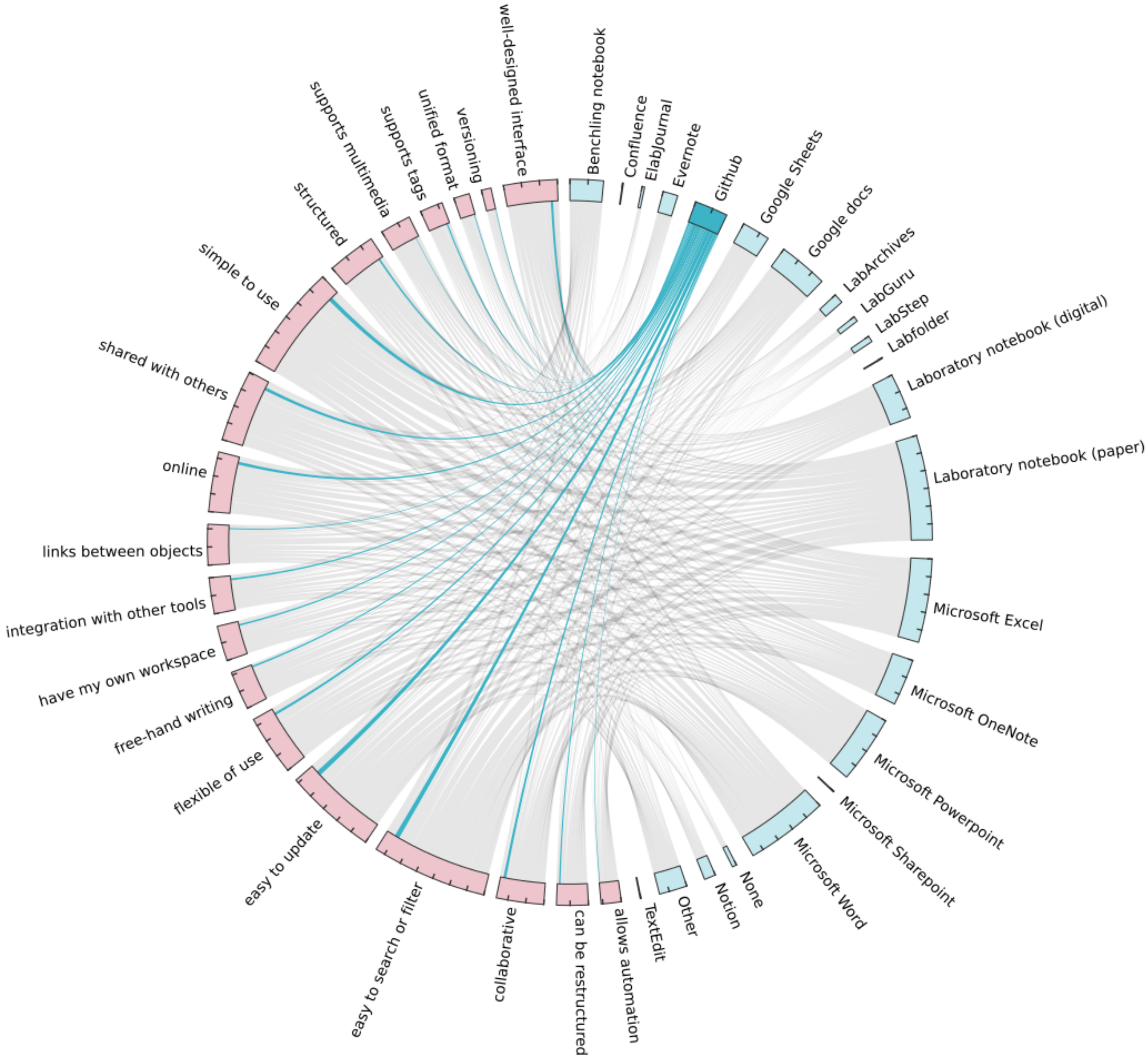


Note-taking solutions (blue arcs) vs desired features (red arcs). Arc length depicts the number of respondents, with ticks denoting 20 respondents. Link thickness shows the fraction of respondents using a solution and valuing a criterion. The chords connected to the arc 'Github' are highlighted in color.

Google Sheets

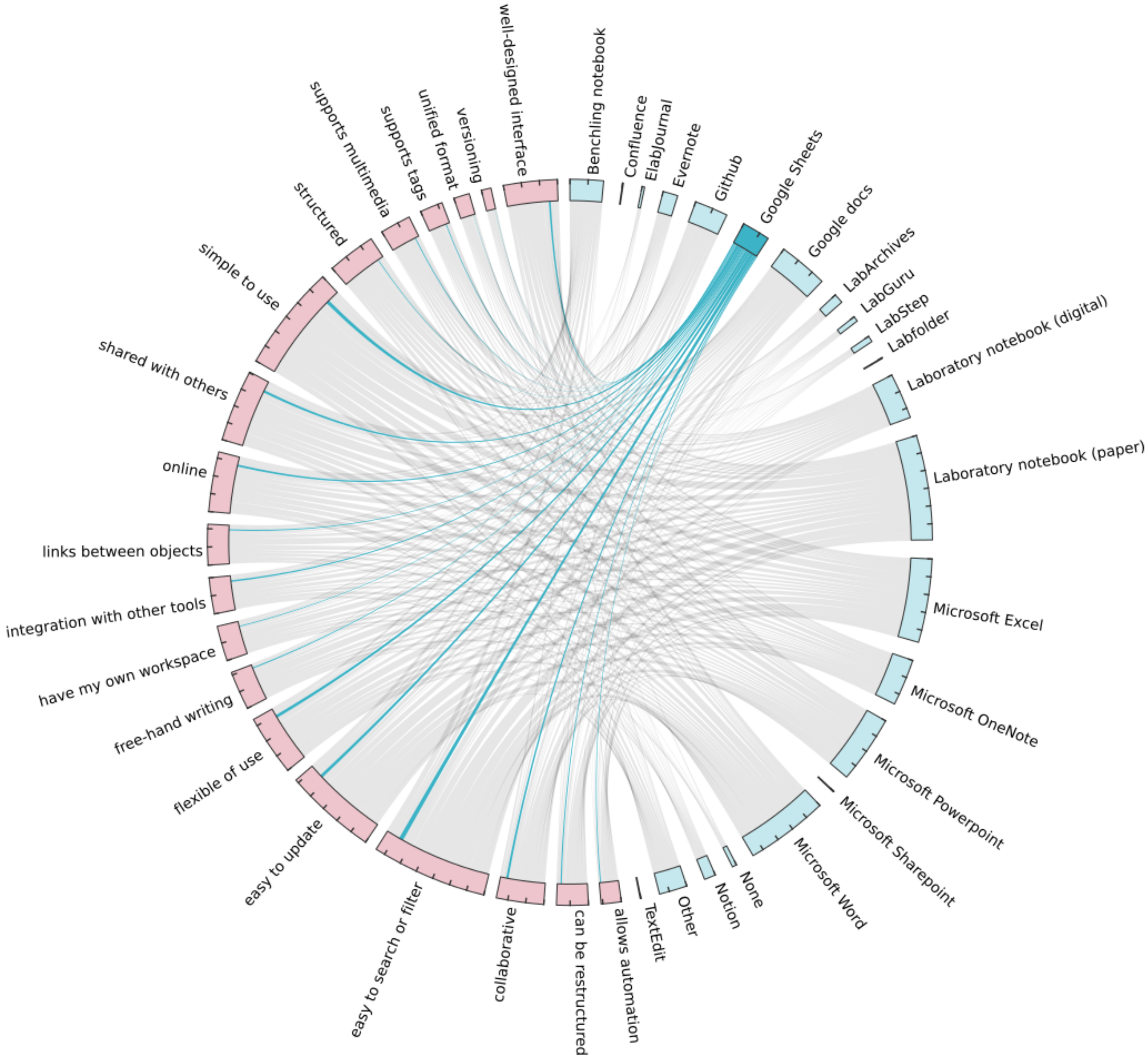


Note-taking solutions (blue arcs) vs desired features (red arcs). Arc length depicts the number of respondents, with ticks denoting 20 respondents. Link thickness shows the fraction of respondents using a solution and valuing a criterion. The chords connected to the arc 'Google Sheets' are highlighted in color.

Google docs

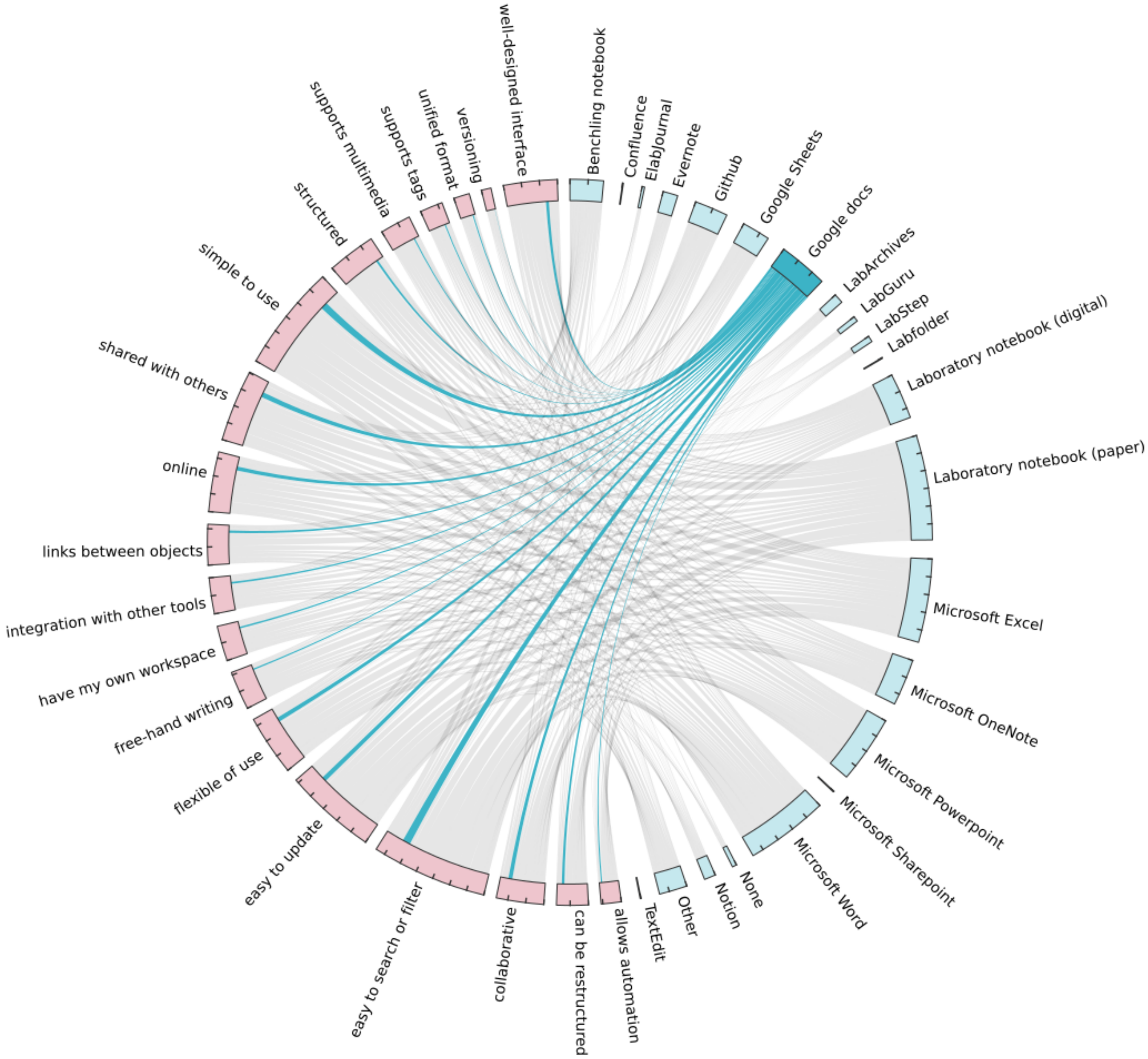


Note-taking solutions (blue arcs) vs desired features (red arcs). Arc length depicts the number of respondents, with ticks denoting 20 respondents. Link thickness shows the fraction of respondents using a solution and valuing a criterion. The chords connected to the arc 'Google docs' are highlighted in color.

LabArchives

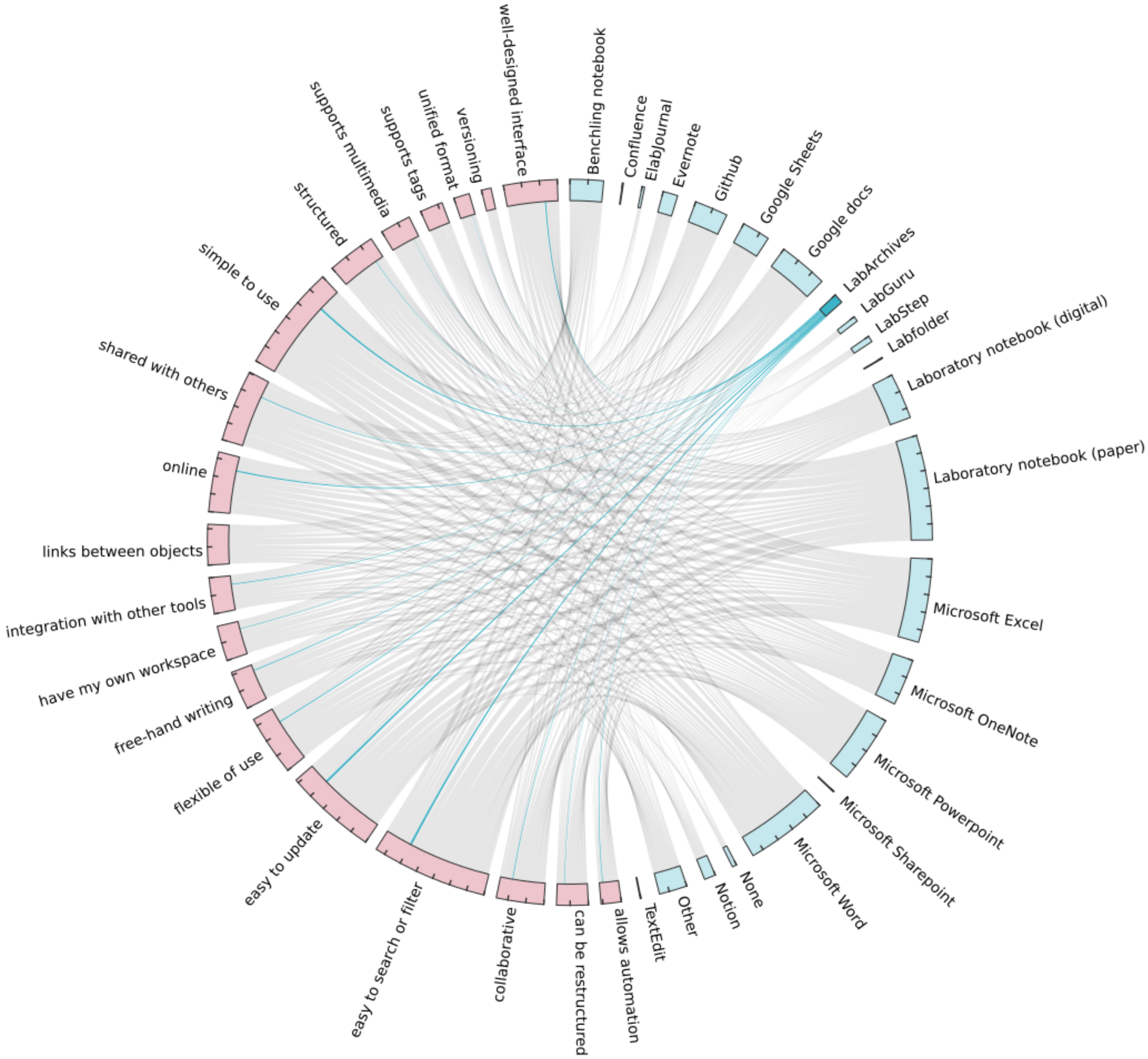


Note-taking solutions (blue arcs) vs desired features (red arcs). Arc length depicts the number of respondents, with ticks denoting 20 respondents. Link thickness shows the fraction of respondents using a solution and valuing a criterion. The chords connected to the arc 'LabArchives' are highlighted in color.

LabGuru

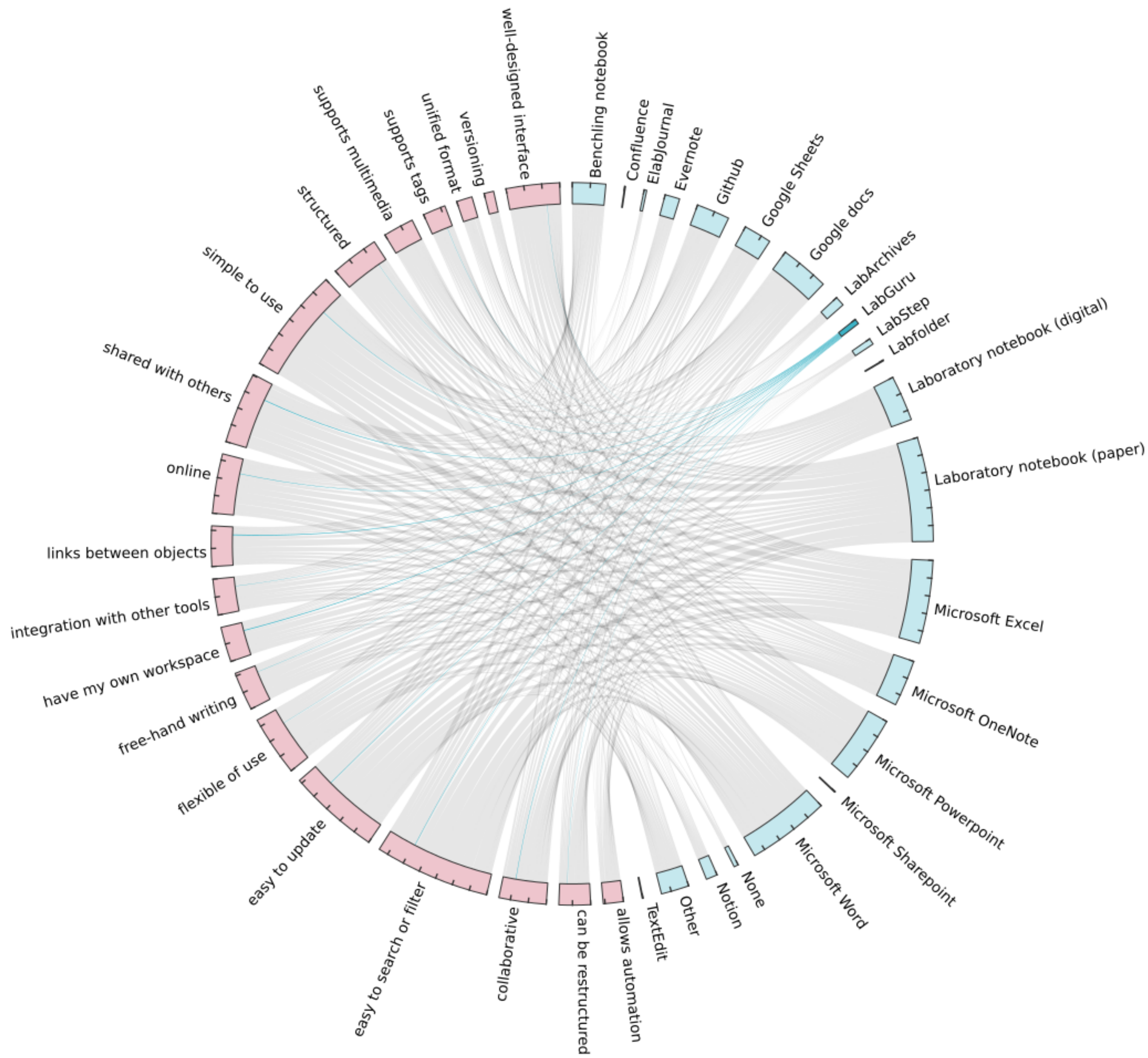


Note-taking solutions (blue arcs) vs desired features (red arcs). Arc length depicts the number of respondents, with ticks denoting 20 respondents. Link thickness shows the fraction of respondents using a solution and valuing a criterion. The chords connected to the arc 'LabGuru' are highlighted in color.

LabStep

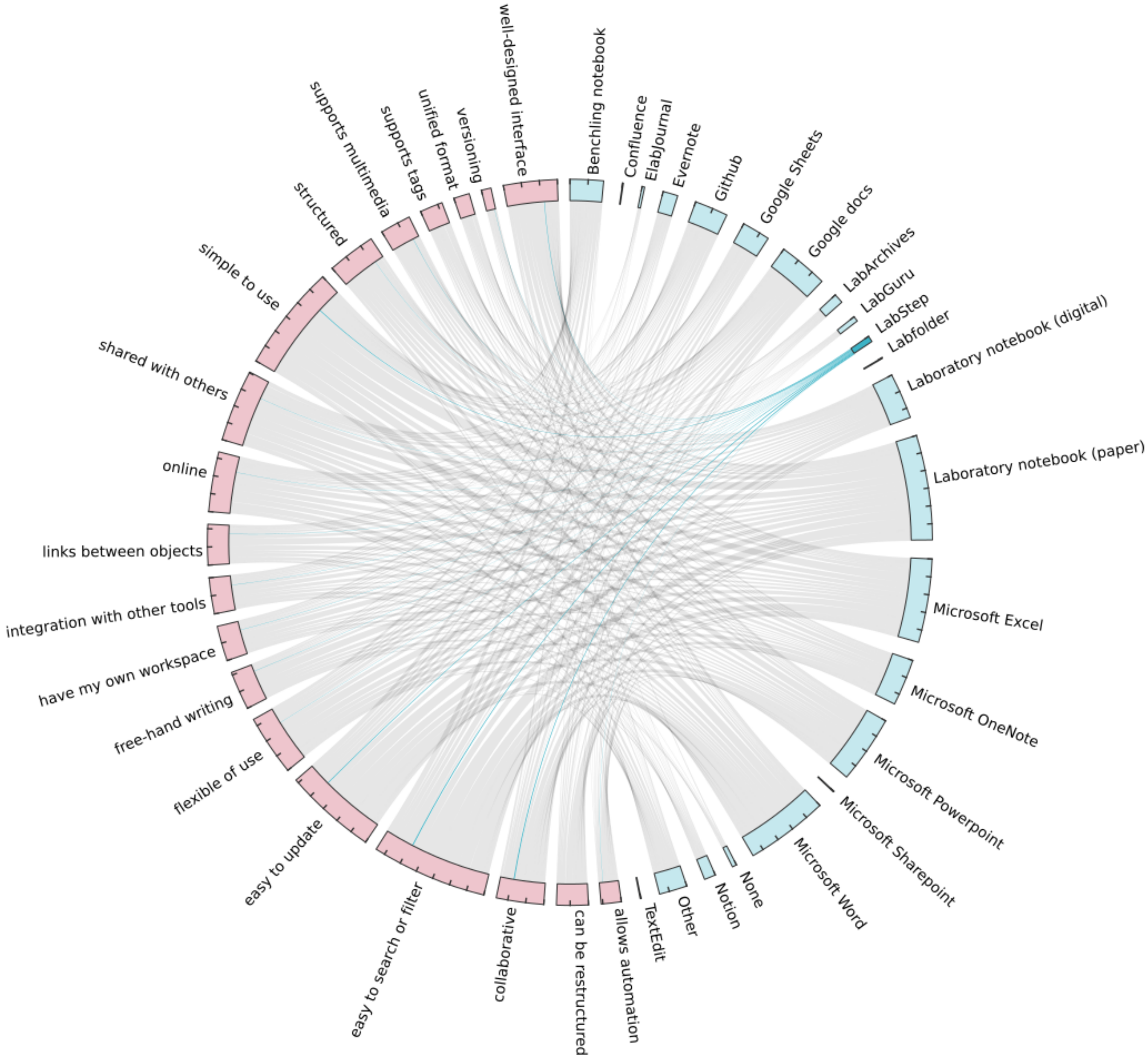


Note-taking solutions (blue arcs) vs desired features (red arcs). Arc length depicts the number of respondents, with ticks denoting 20 respondents. Link thickness shows the fraction of respondents using a solution and valuing a criterion. The chords connected to the arc 'LabStep' are highlighted in color.

Labfolder

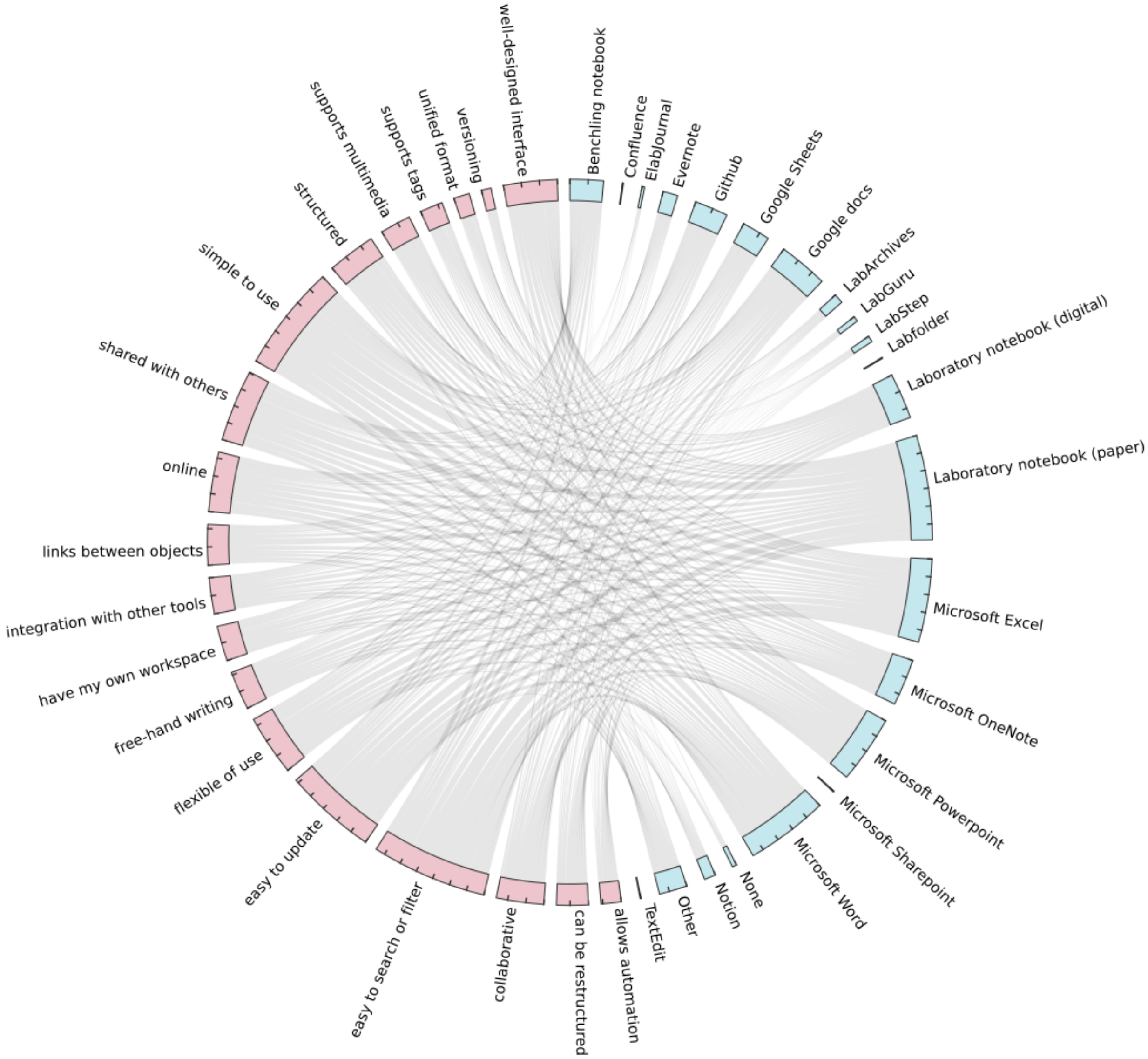


Note-taking solutions (blue arcs) vs desired features (red arcs). Arc length depicts the number of respondents, with ticks denoting 20 respondents. Link thickness shows the fraction of respondents using a solution and valuing a criterion. The chords connected to the arc 'Labfolder' are highlighted in color.

Laboratory notebook (digital)

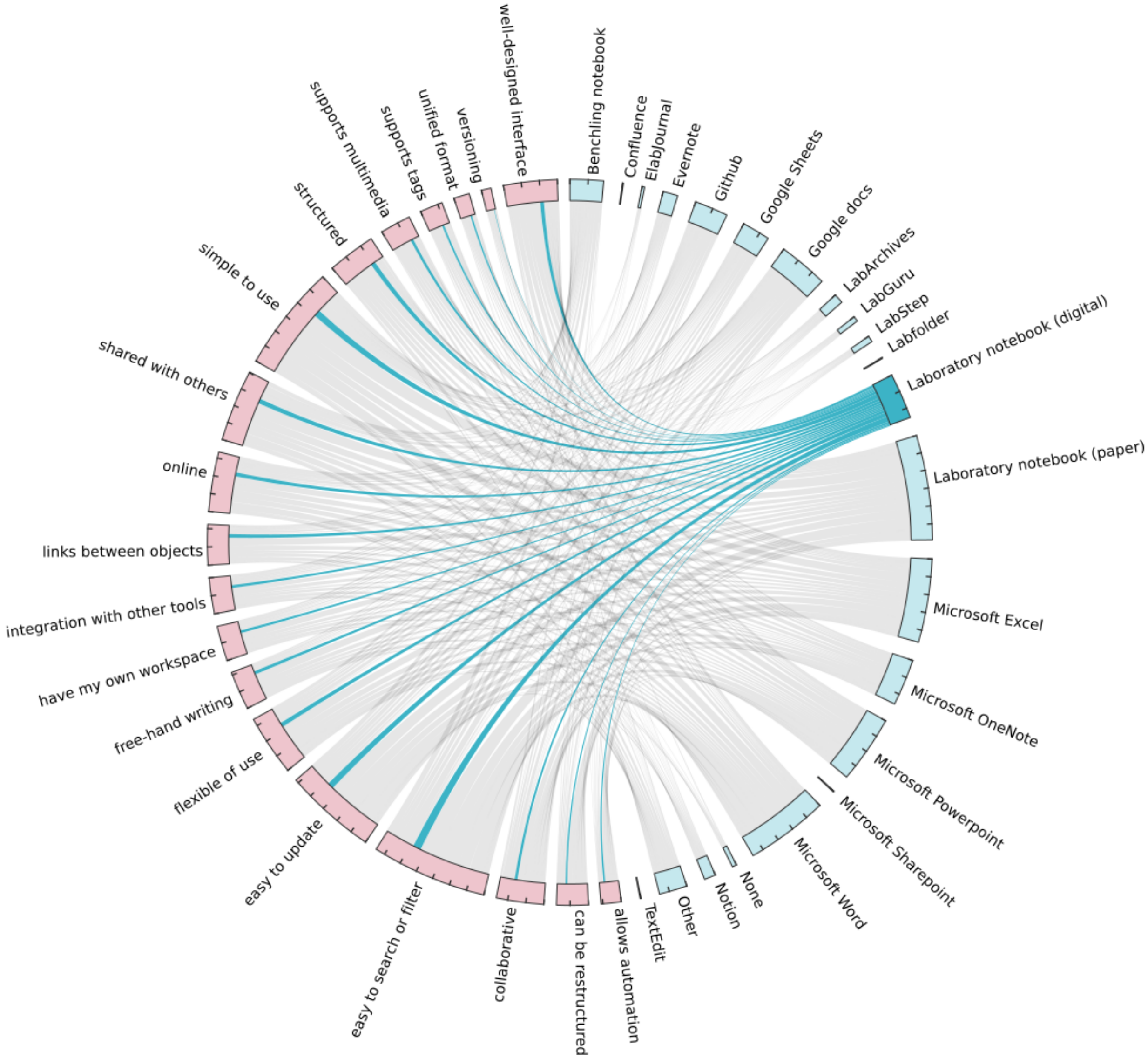


Note-taking solutions (blue arcs) vs desired features (red arcs). Arc length depicts the number of respondents, with ticks denoting 20 respondents. Link thickness shows the fraction of respondents using a solution and valuing a criterion. The chords connected to the arc 'Laboratory notebook (digital)' are highlighted in color.

Laboratory notebook (paper)

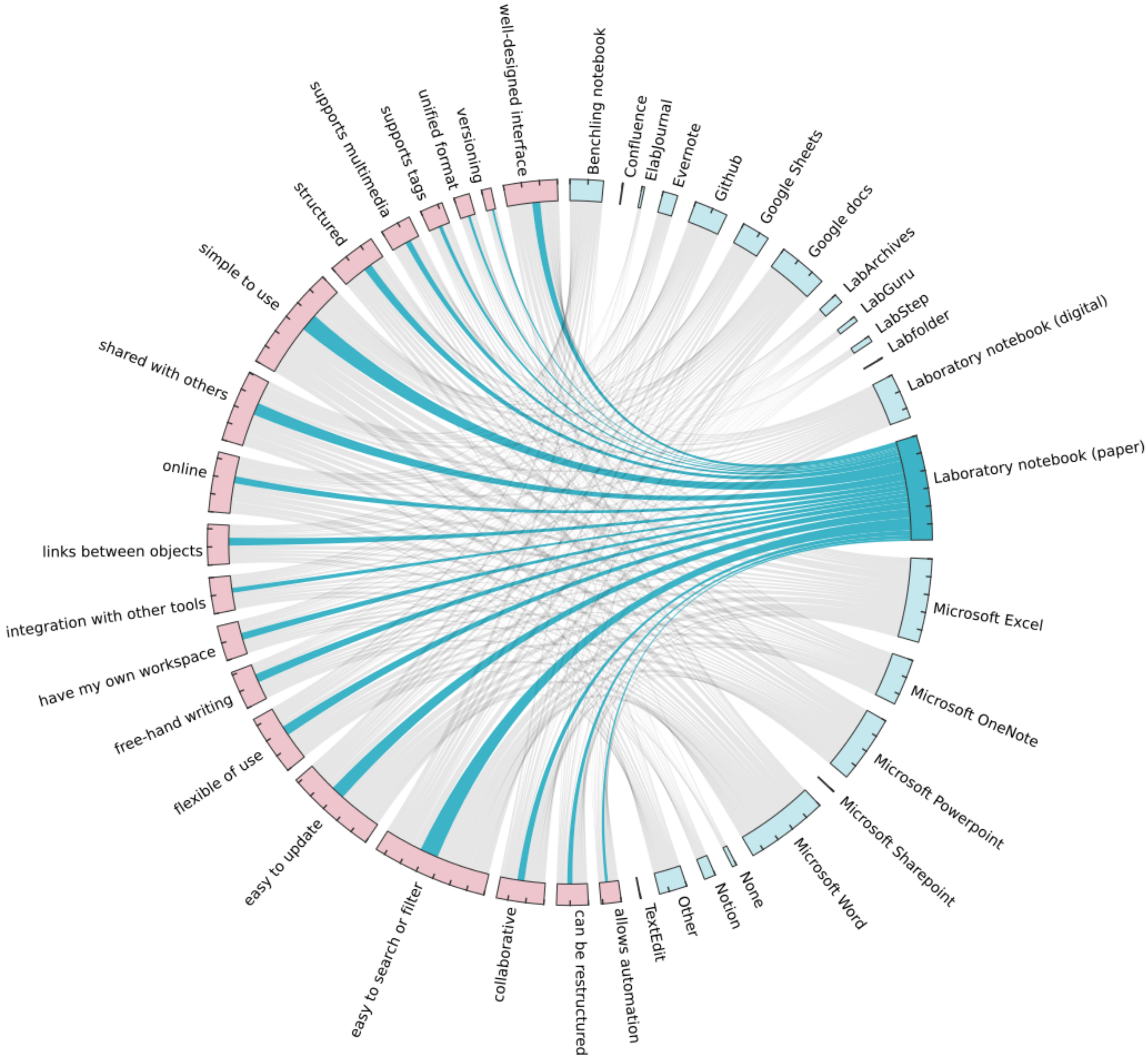


Note-taking solutions (blue arcs) vs desired features (red arcs). Arc length depicts the number of respondents, with ticks denoting 20 respondents. Link thickness shows the fraction of respondents using a solution and valuing a criterion. The chords connected to the arc 'Laboratory notebook (paper)' are highlighted in color.

Microsoft Excel

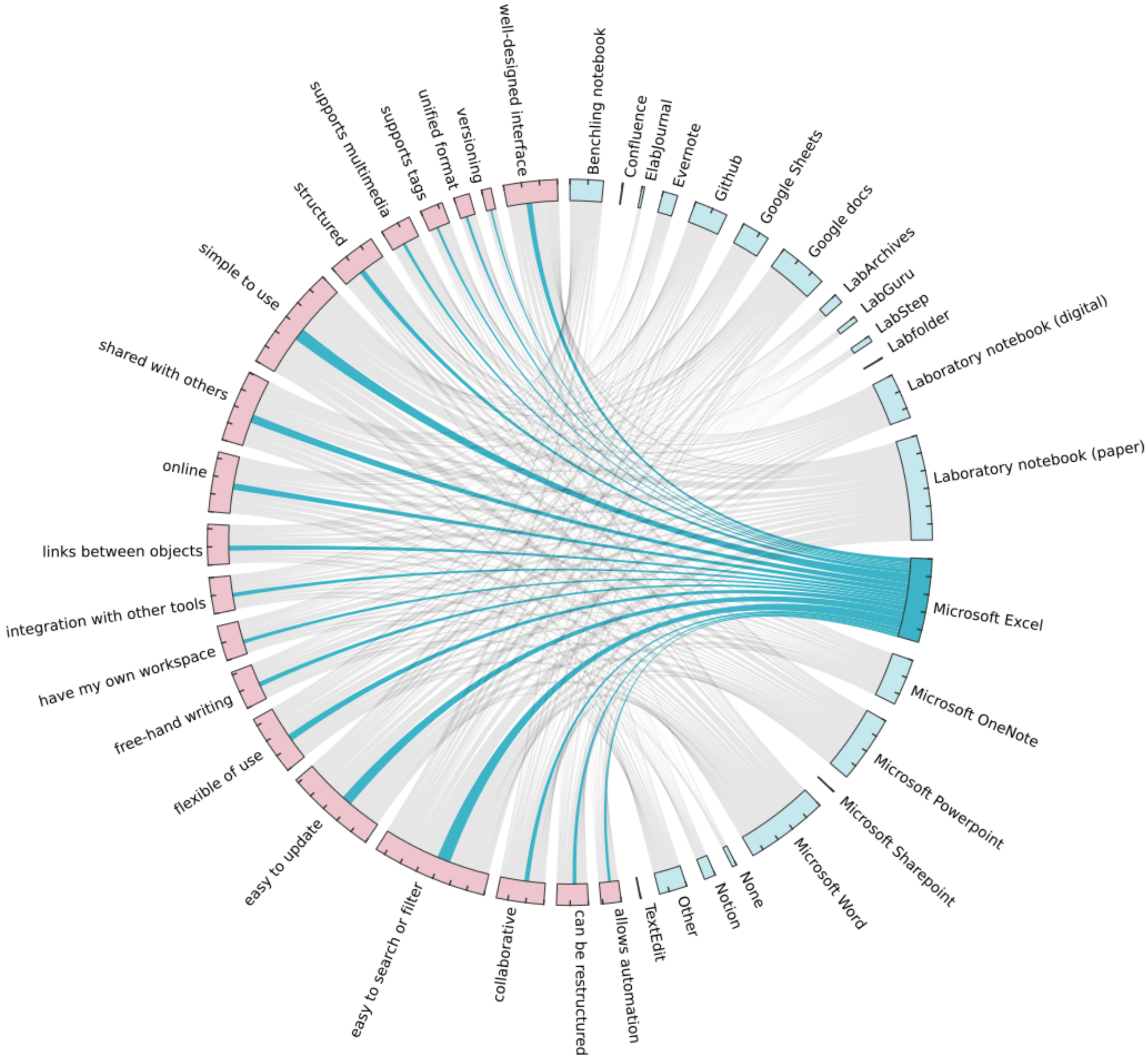


Note-taking solutions (blue arcs) vs desired features (red arcs). Arc length depicts the number of respondents, with ticks denoting 20 respondents. Link thickness shows the fraction of respondents using a solution and valuing a criterion. The chords connected to the arc 'Microsoft Excel' are highlighted in color.

Microsoft OneNote

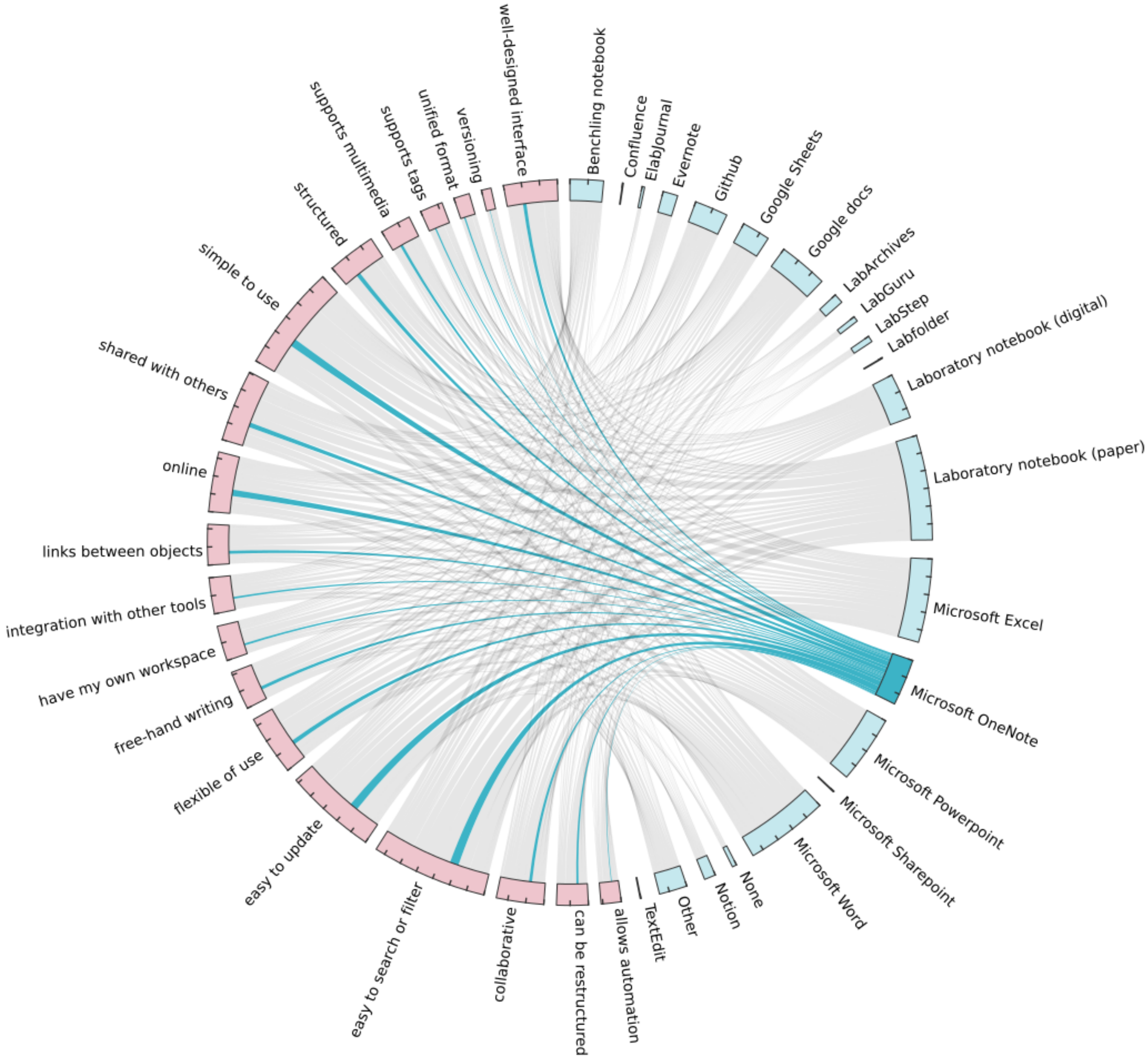


Note-taking solutions (blue arcs) vs desired features (red arcs). Arc length depicts the number of respondents, with ticks denoting 20 respondents. Link thickness shows the fraction of respondents using a solution and valuing a criterion. The chords connected to the arc 'Microsoft OneNote' are highlighted in color.

Microsoft Powerpoint

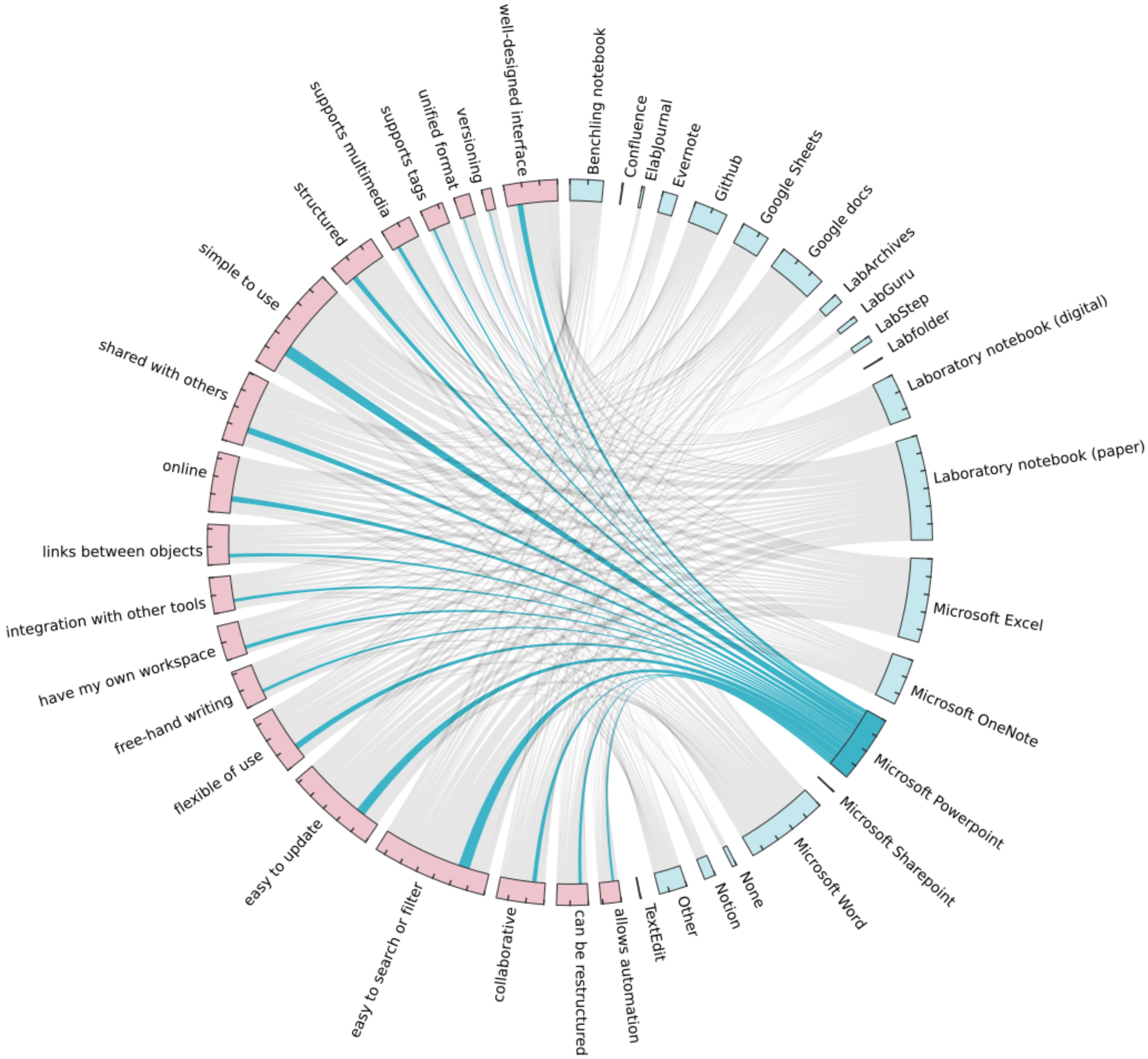


Note-taking solutions (blue arcs) vs desired features (red arcs). Arc length depicts the number of respondents, with ticks denoting 20 respondents. Link thickness shows the fraction of respondents using a solution and valuing a criterion. The chords connected to the arc 'Microsoft Powerpoint' are highlighted in color.

Microsoft Sharepoint

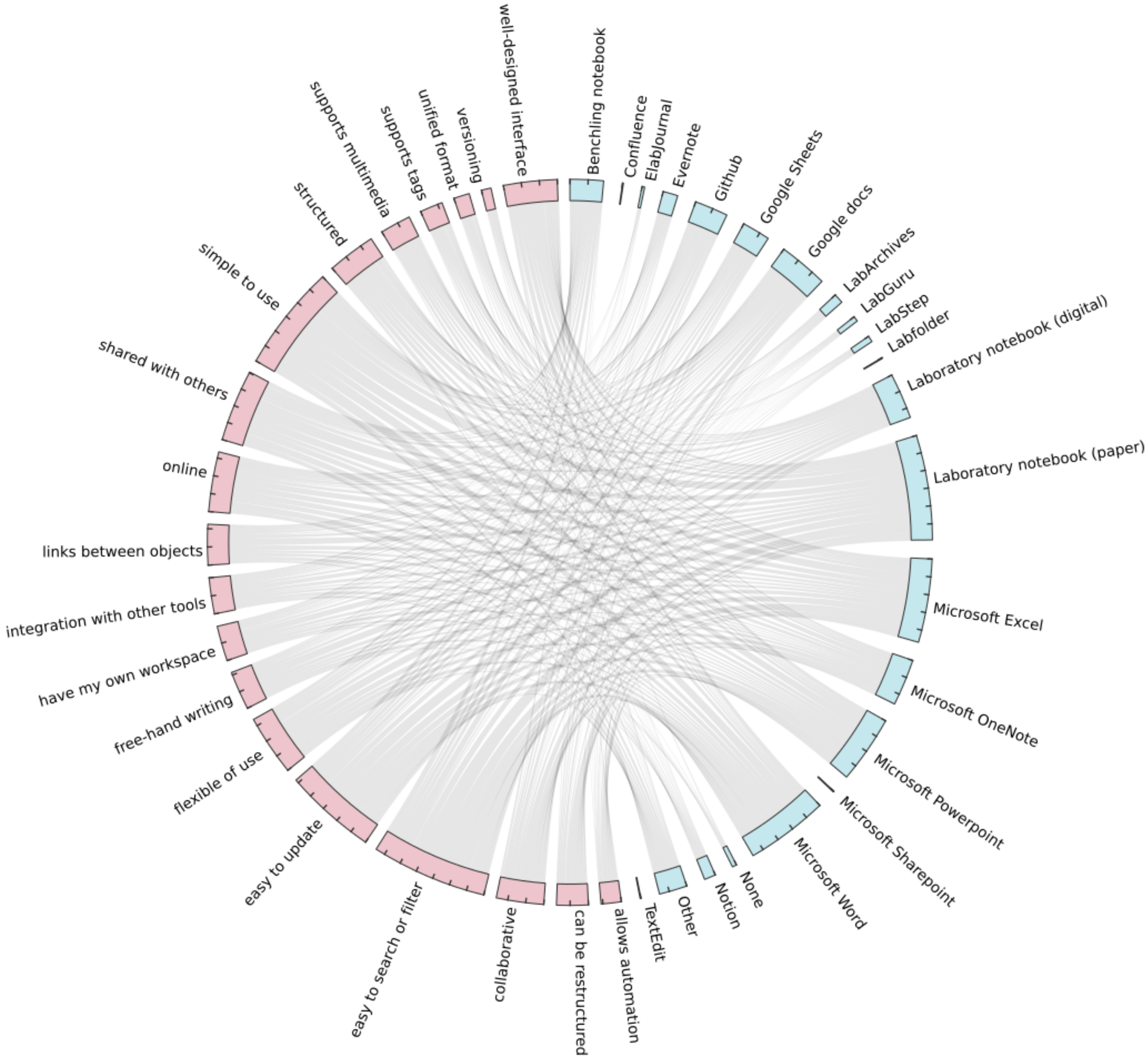


Note-taking solutions (blue arcs) vs desired features (red arcs). Arc length depicts the number of respondents, with ticks denoting 20 respondents. Link thickness shows the fraction of respondents using a solution and valuing a criterion. The chords connected to the arc 'Microsoft Sharepoint' are highlighted in color.

Microsoft Word

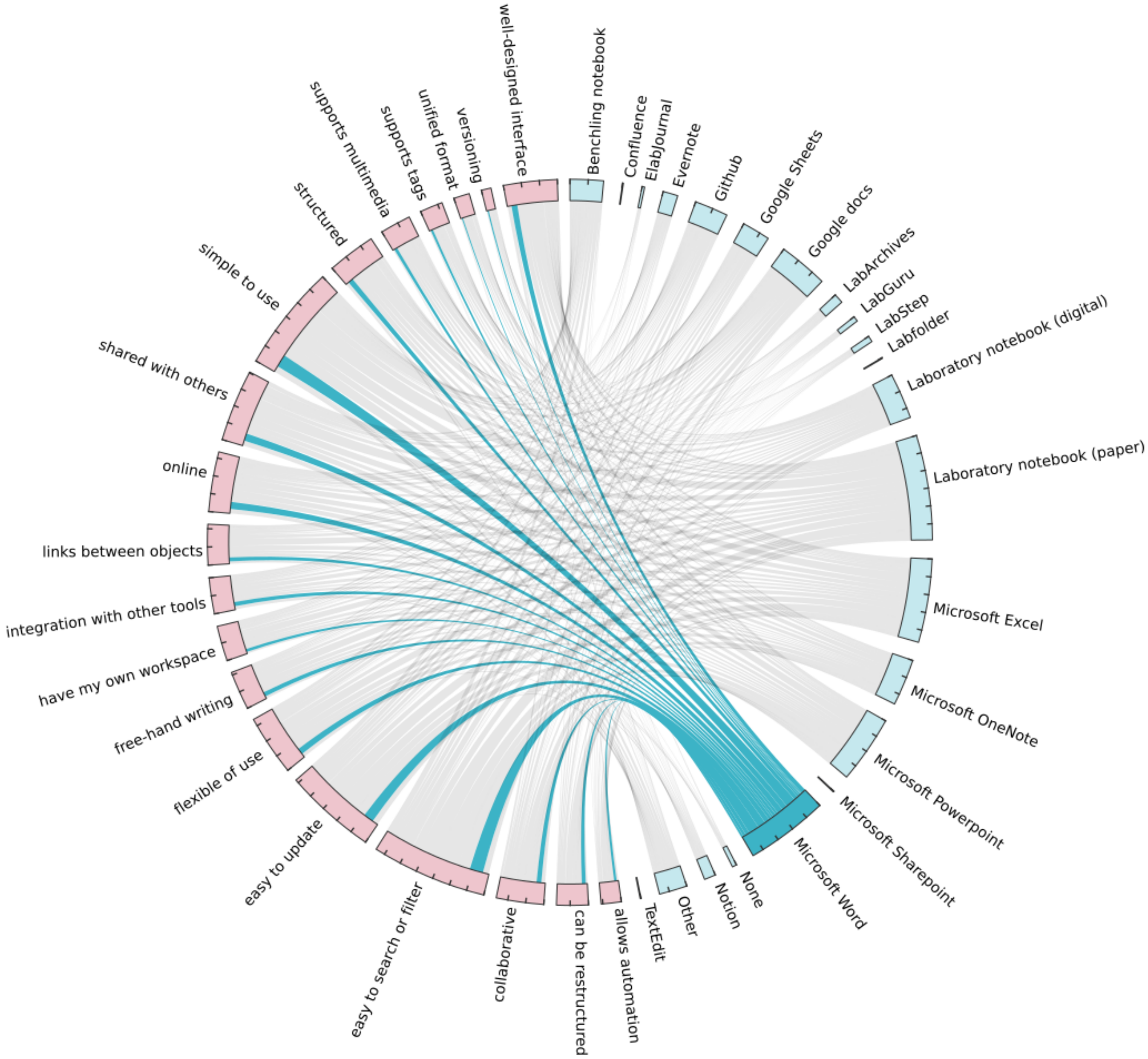


Note-taking solutions (blue arcs) vs desired features (red arcs). Arc length depicts the number of respondents, with ticks denoting 20 respondents. Link thickness shows the fraction of respondents using a solution and valuing a criterion. The chords connected to the arc 'Microsoft Word' are highlighted in color.

None

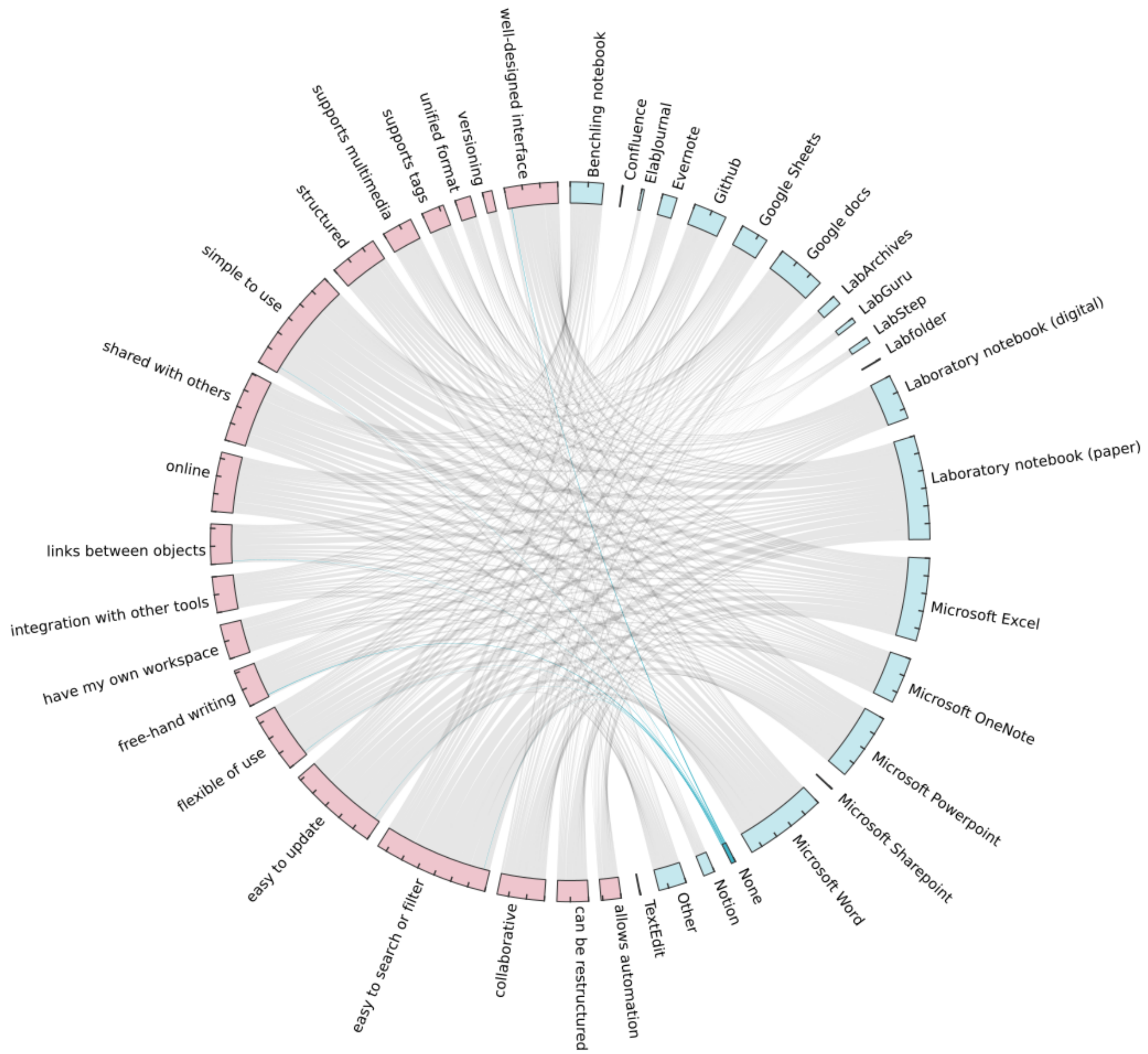


Note-taking solutions (blue arcs) vs desired features (red arcs). Arc length depicts the number of respondents, with ticks denoting 20 respondents. Link thickness shows the fraction of respondents using a solution and valuing a criterion. The chords connected to the arc 'None' are highlighted in color.

Notion

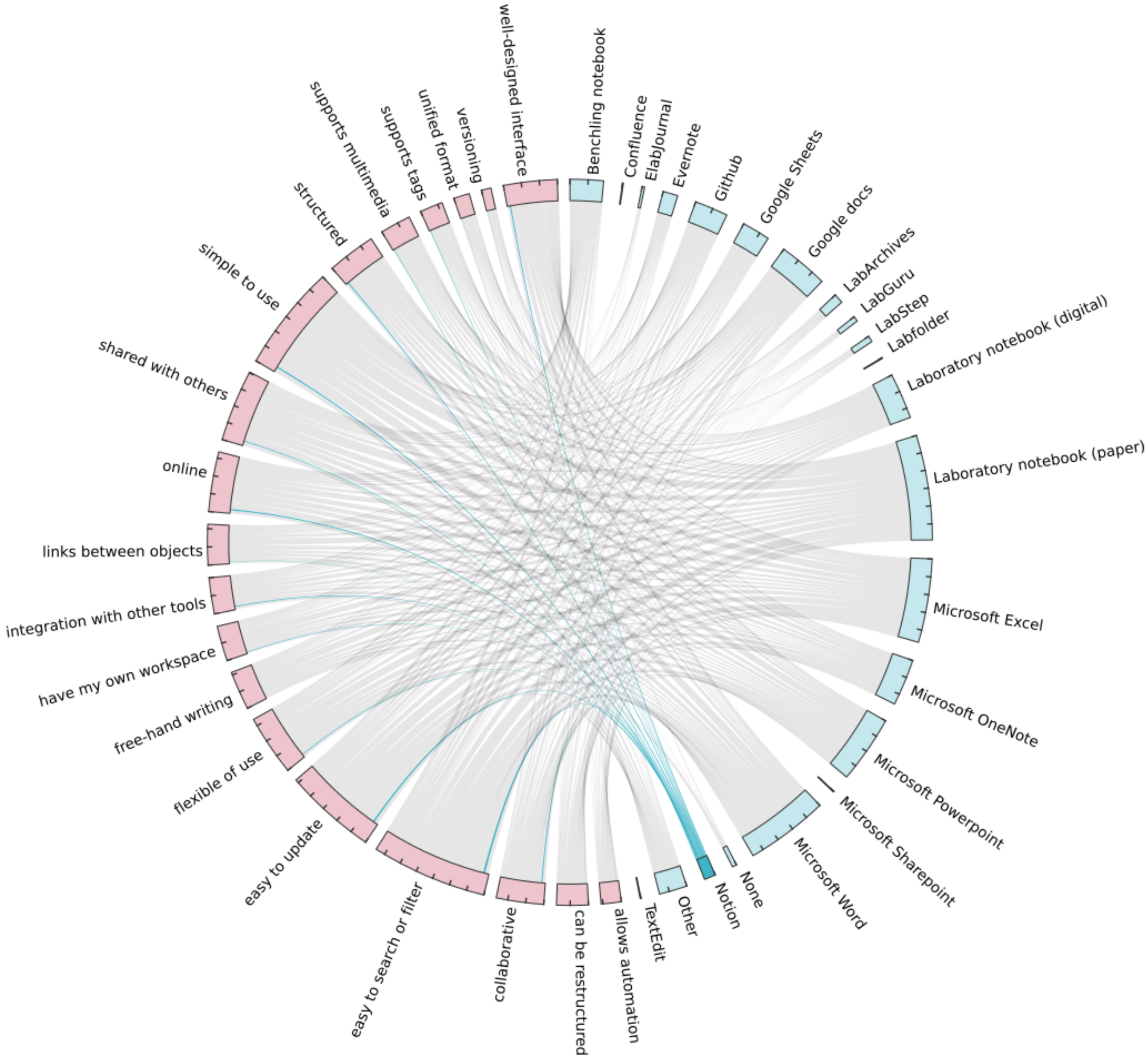


Note-taking solutions (blue arcs) vs desired features (red arcs). Arc length depicts the number of respondents, with ticks denoting 20 respondents. Link thickness shows the fraction of respondents using a solution and valuing a criterion. The chords connected to the arc 'Notion' are highlighted in color.

Other

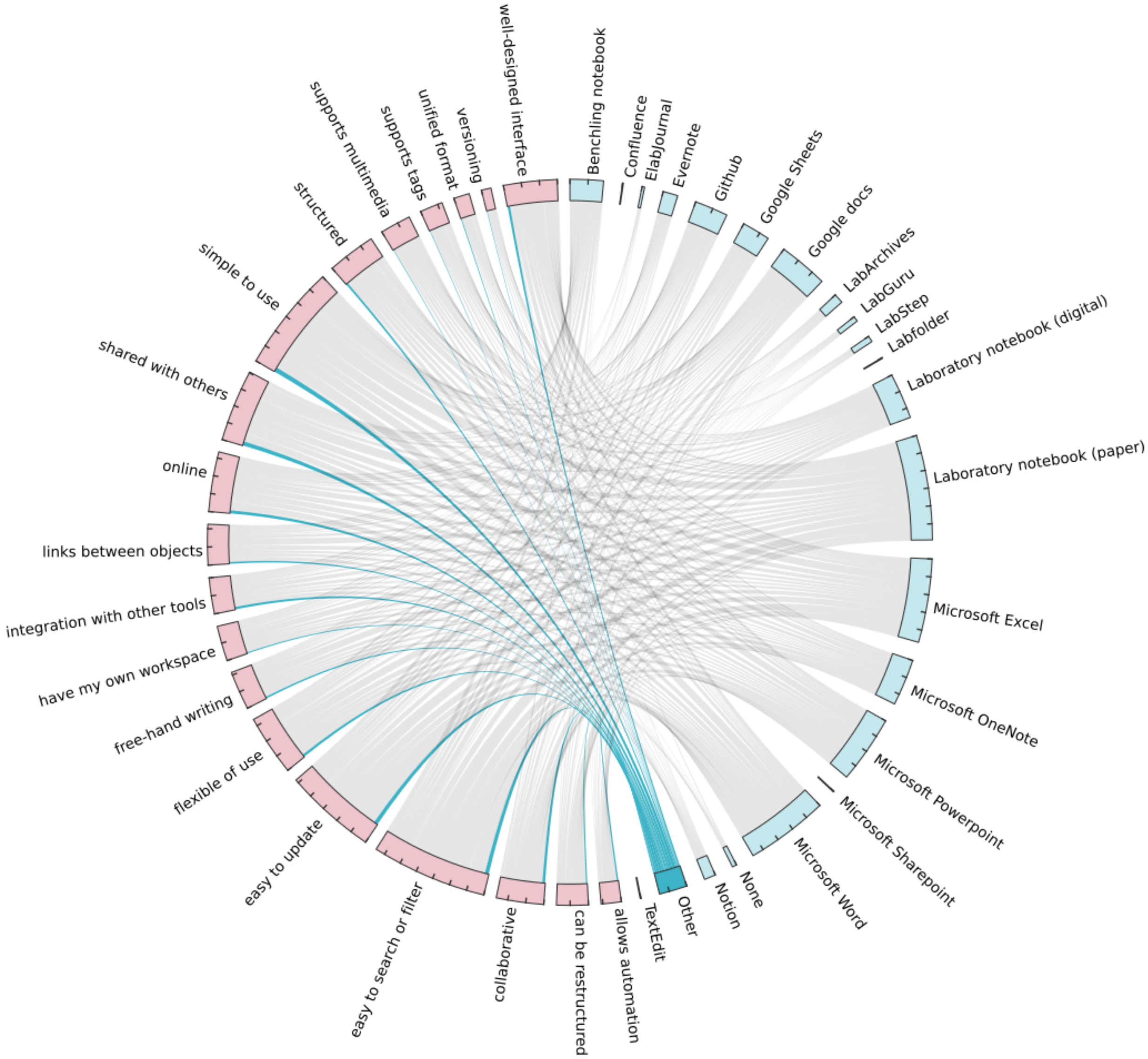


Note-taking solutions (blue arcs) vs desired features (red arcs). Arc length depicts the number of respondents, with ticks denoting 20 respondents. Link thickness shows the fraction of respondents using a solution and valuing a criterion. The chords connected to the arc 'Other' are highlighted in color.

TextEdit

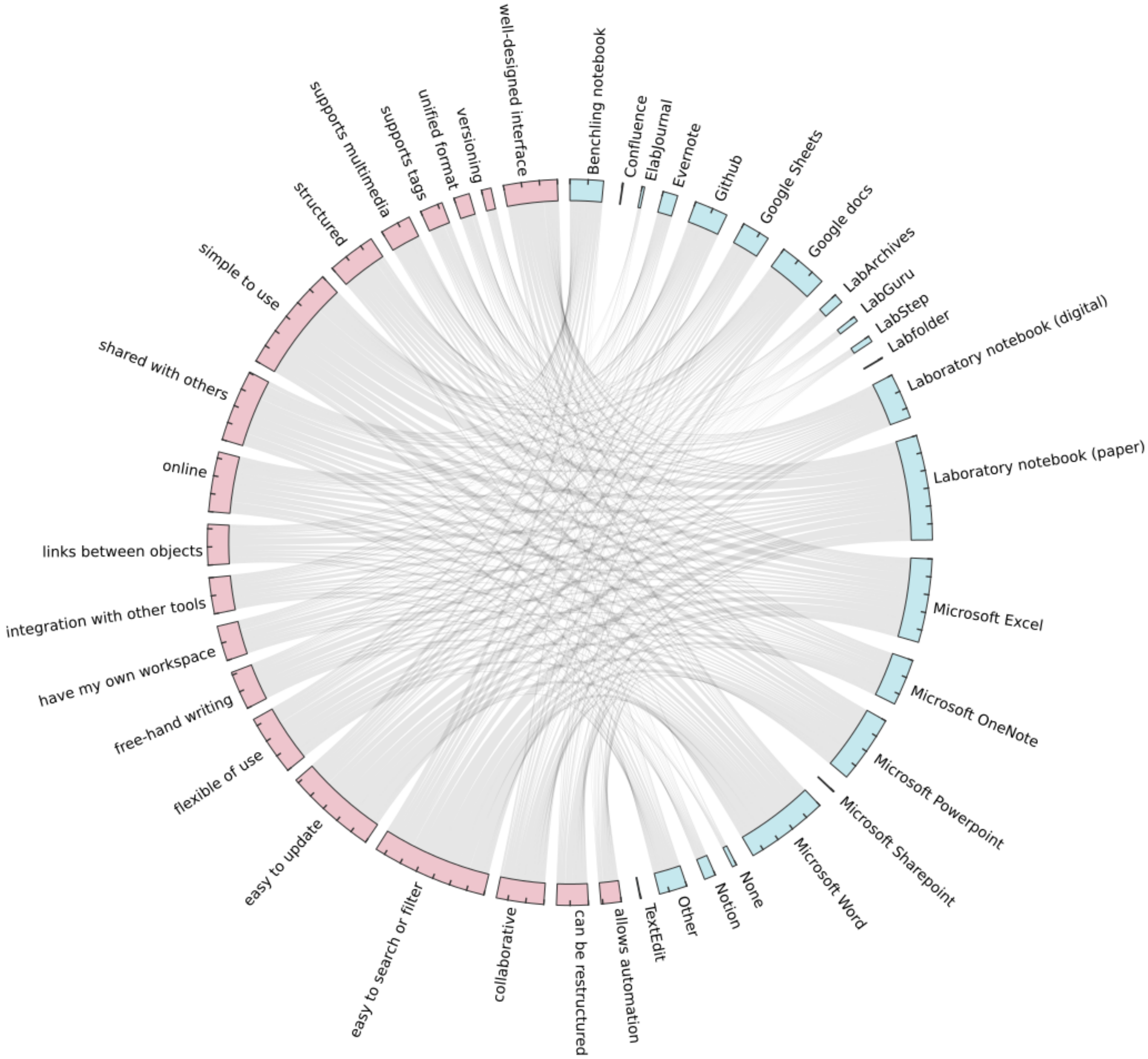


Note-taking solutions (blue arcs) vs desired features (red arcs). Arc length depicts the number of respondents, with ticks denoting 20 respondents. Link thickness shows the fraction of respondents using a solution and valuing a criterion. The chords connected to the arc 'TextEdit' are highlighted in color.

allows automation

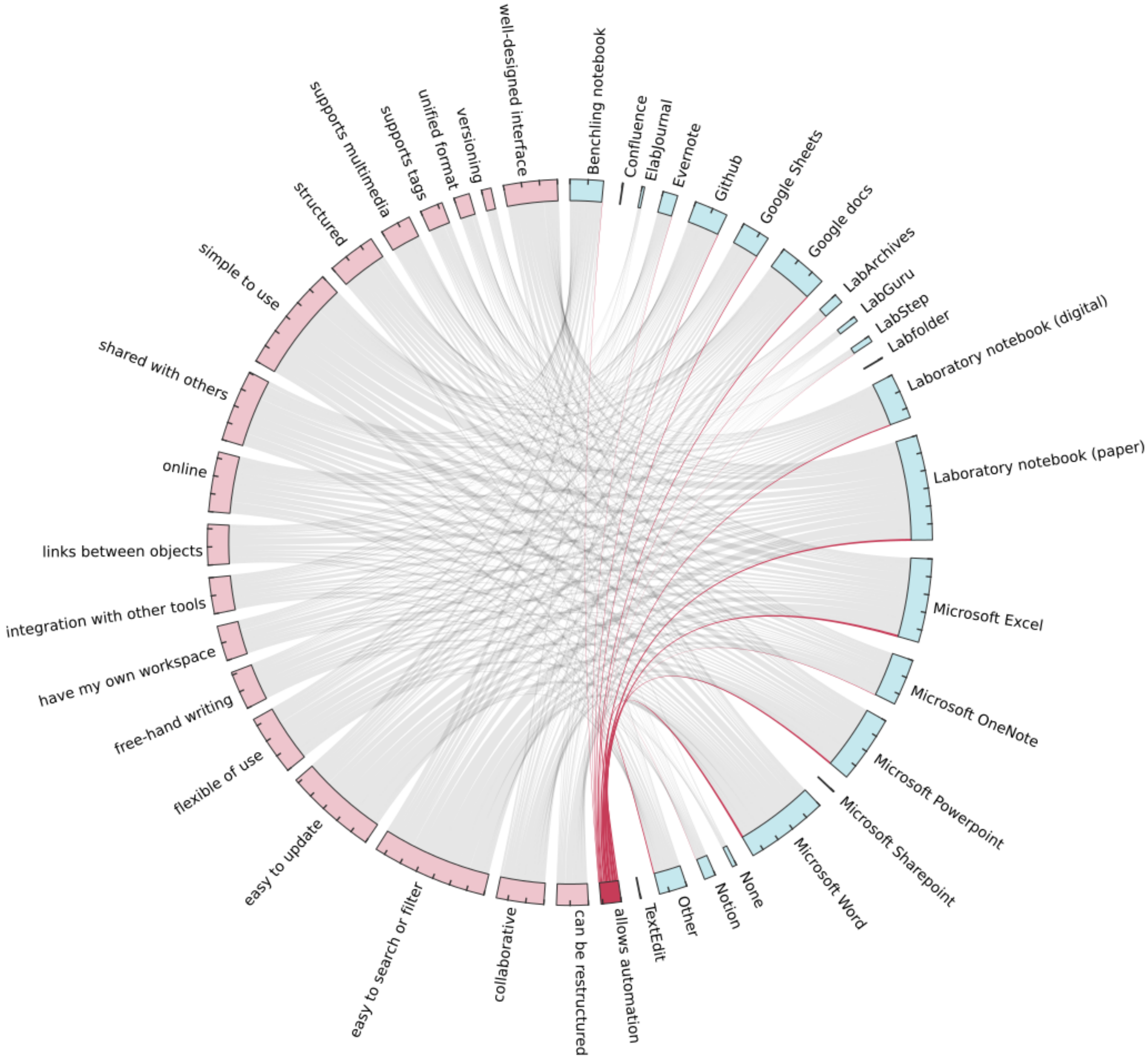


Note-taking solutions (blue arcs) vs desired features (red arcs). Arc length depicts the number of respondents, with ticks denoting 20 respondents. Link thickness shows the fraction of respondents using a solution and valuing a criterion. The chords connected to the arc 'allows automation' are highlighted in color.

can be restructured

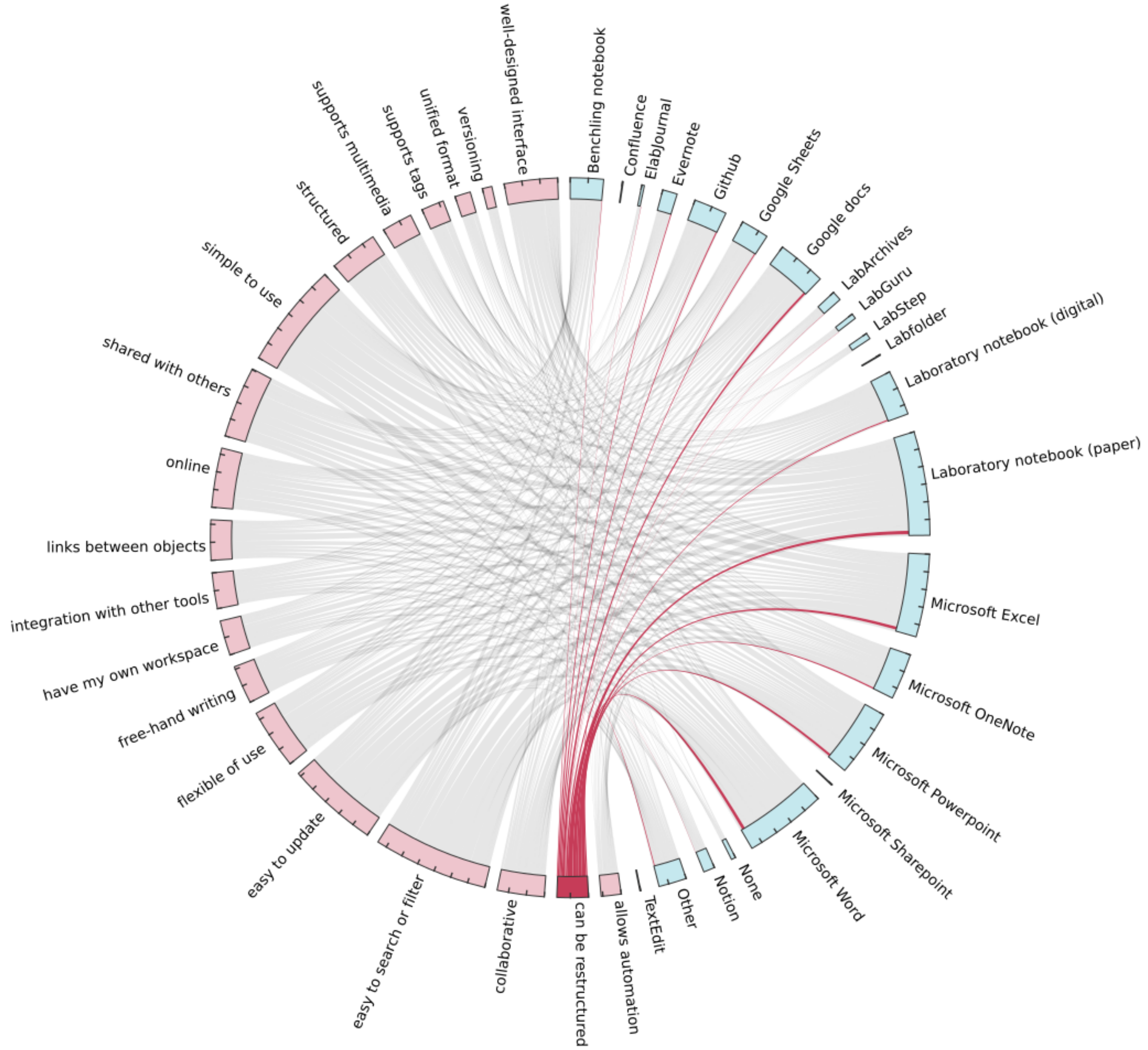


Note-taking solutions (blue arcs) vs desired features (red arcs). Arc length depicts the number of respondents, with ticks denoting 20 respondents. Link thickness shows the fraction of respondents using a solution and valuing a criterion. The chords connected to the arc 'can be restructured' are highlighted in color.

collaborative

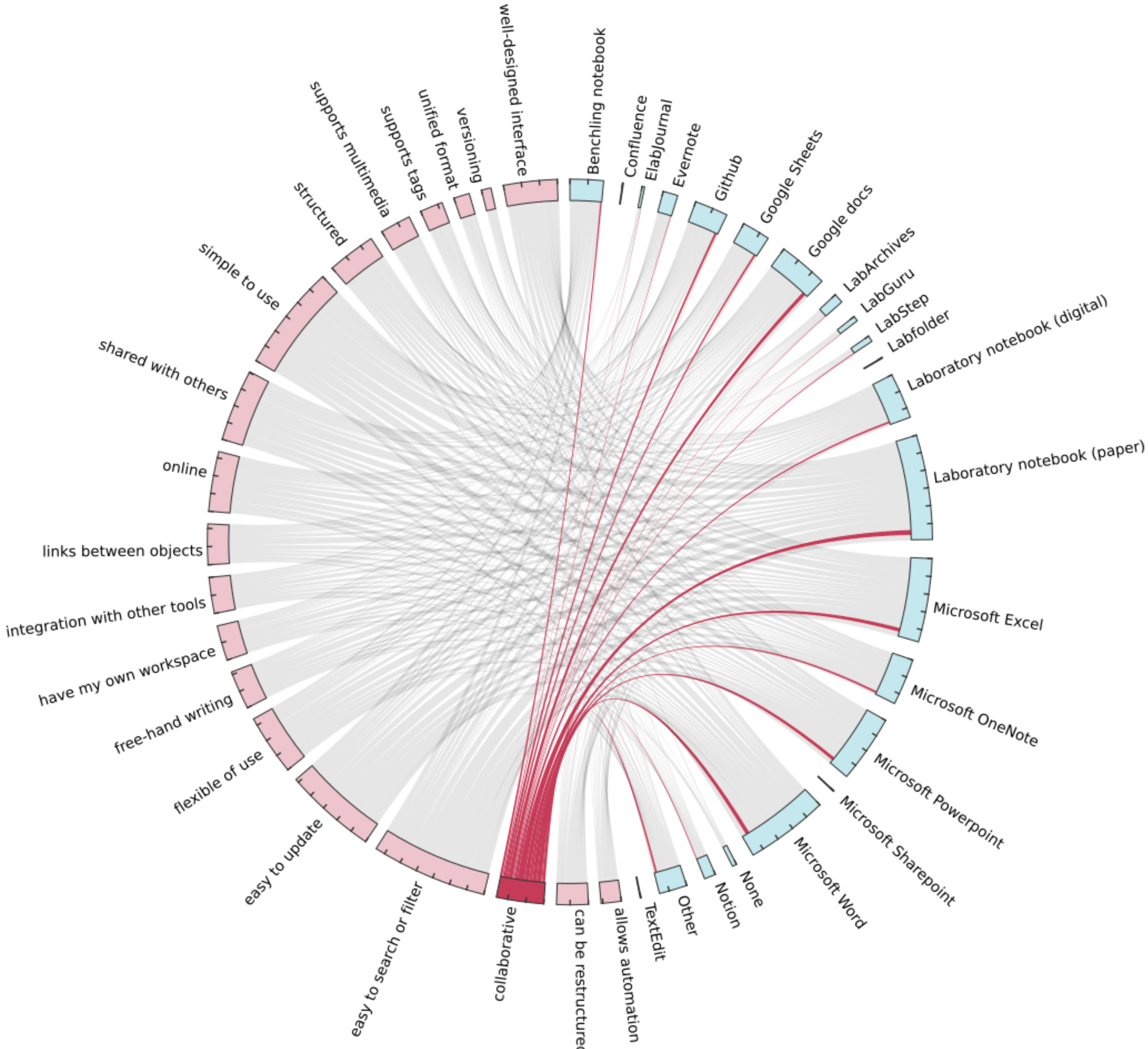


Note-taking solutions (blue arcs) vs desired features (red arcs). Arc length depicts the number of respondents, with ticks denoting 20 respondents. Link thickness shows the fraction of respondents using a solution and valuing a criterion. The chords connected to the arc 'collaborative' are highlighted in color.

easy to search or filter

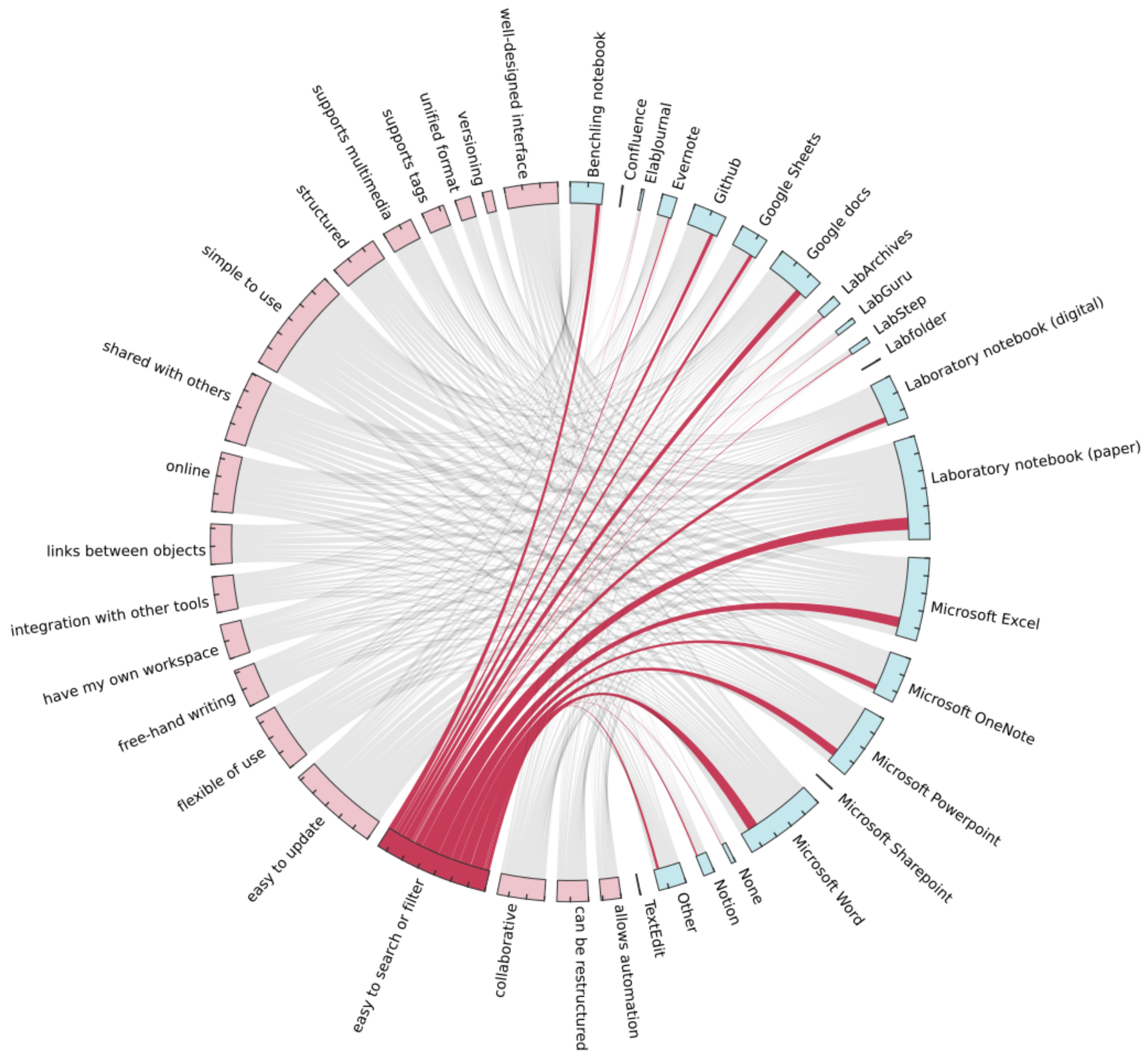


Note-taking solutions (blue arcs) vs desired features (red arcs). Arc length depicts the number of respondents, with ticks denoting 20 respondents. Link thickness shows the fraction of respondents using a solution and valuing a criterion. The chords connected to the arc 'easy to search or filter' are highlighted in color.

easy to update

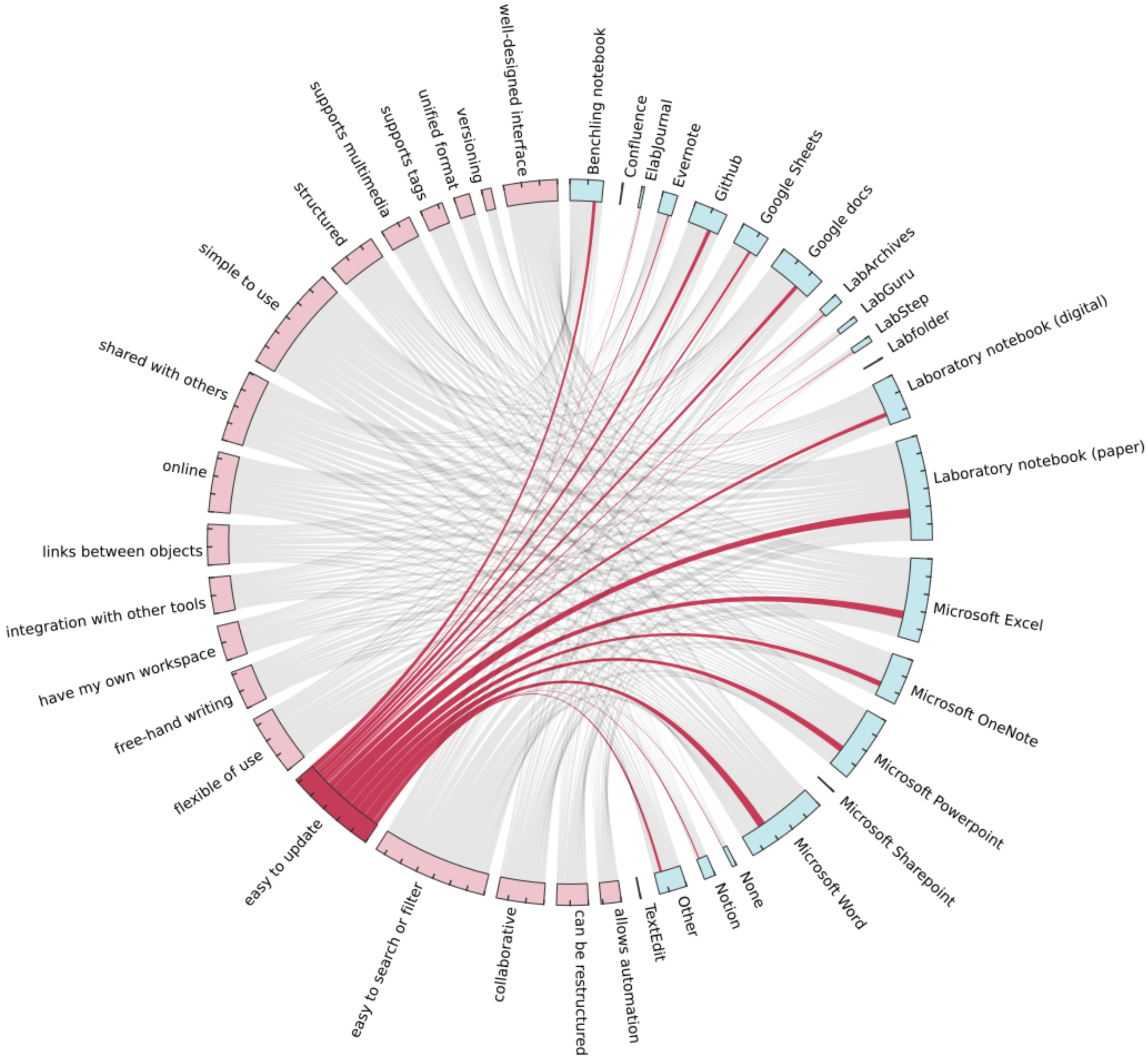


Note-taking solutions (blue arcs) vs desired features (red arcs). Arc length depicts the number of respondents, with ticks denoting 20 respondents. Link thickness shows the fraction of respondents using a solution and valuing a criterion. The chords connected to the arc 'easy to update' are highlighted in color.

flexible of use

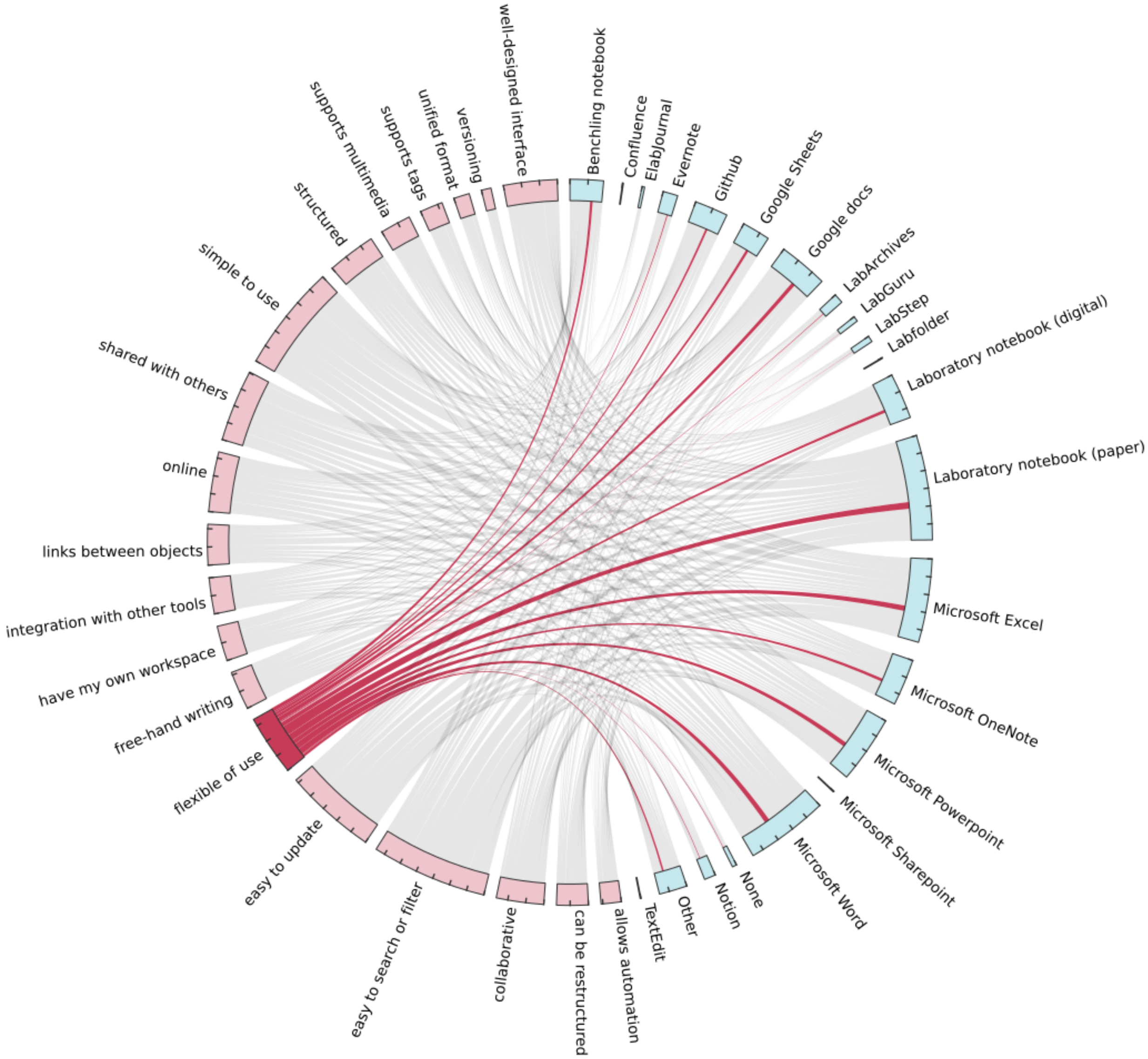


Note-taking solutions (blue arcs) vs desired features (red arcs). Arc length depicts the number of respondents, with ticks denoting 20 respondents. Link thickness shows the fraction of respondents using a solution and valuing a criterion. The chords connected to the arc 'flexible of use' are highlighted in color.

free-hand writing

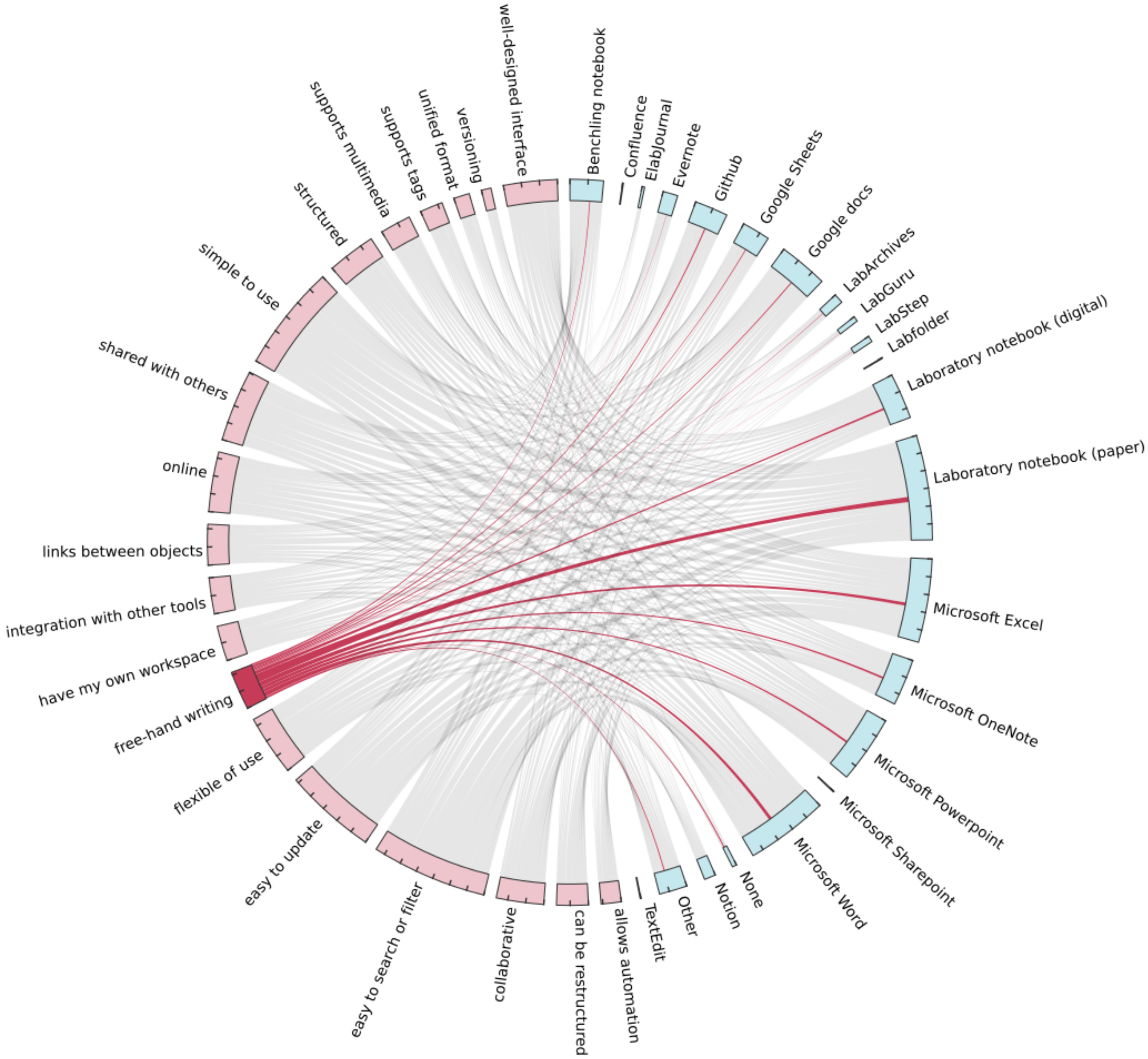


Note-taking solutions (blue arcs) vs desired features (red arcs). Arc length depicts the number of respondents, with ticks denoting 20 respondents. Link thickness shows the fraction of respondents using a solution and valuing a criterion. The chords connected to the arc 'free-hand writing' are highlighted in color.

have my own workspace

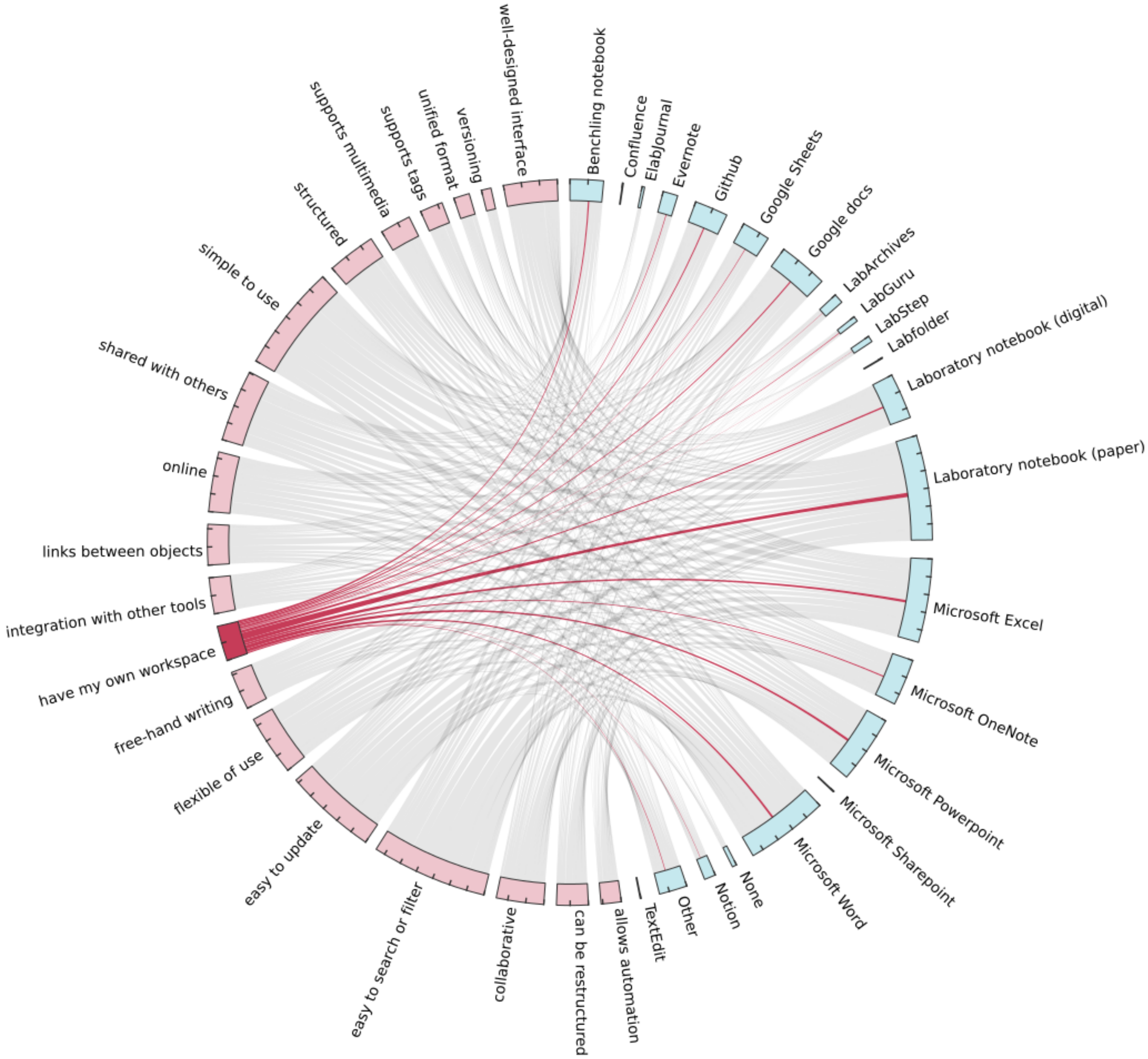


Note-taking solutions (blue arcs) vs desired features (red arcs). Arc length depicts the number of respondents, with ticks denoting 20 respondents. Link thickness shows the fraction of respondents using a solution and valuing a criterion. The chords connected to the arc 'have my own workspace' are highlighted in color.

integration with other tools

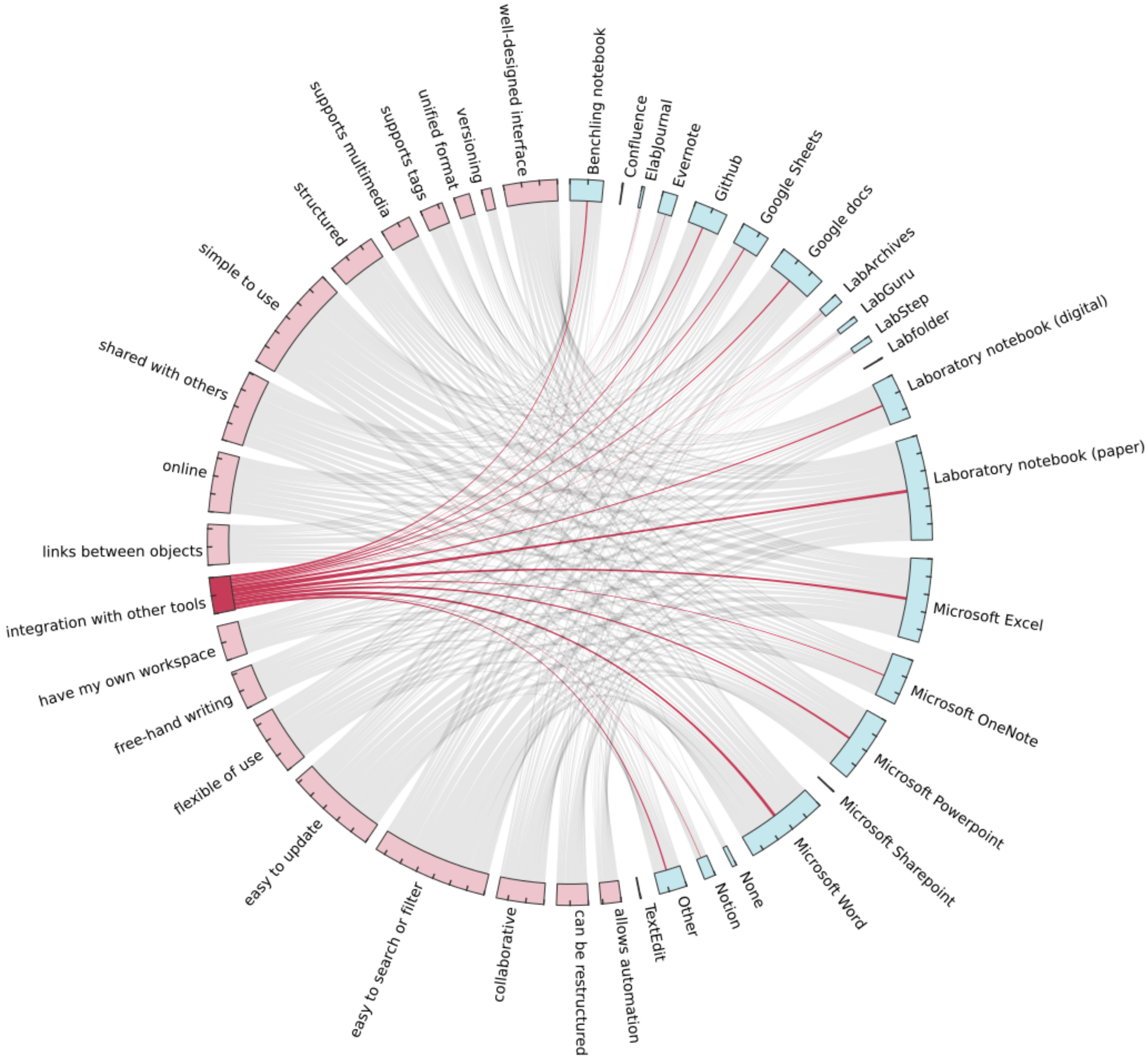


Note-taking solutions (blue arcs) vs desired features (red arcs). Arc length depicts the number of respondents, with ticks denoting 20 respondents. Link thickness shows the fraction of respondents using a solution and valuing a criterion. The chords connected to the arc 'integration with other tools' are highlighted in color.

links between objects

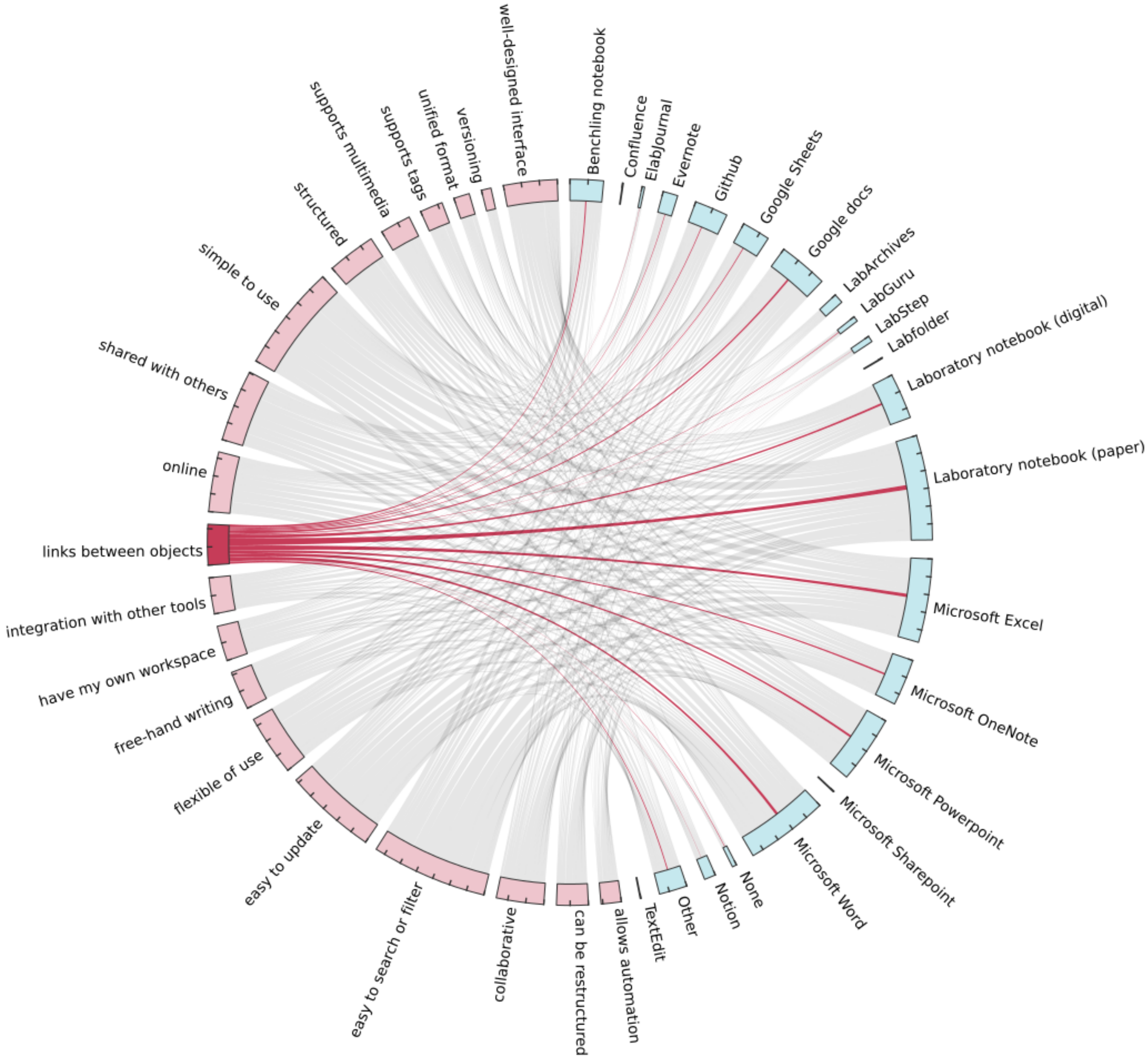


Note-taking solutions (blue arcs) vs desired features (red arcs). Arc length depicts the number of respondents, with ticks denoting 20 respondents. Link thickness shows the fraction of respondents using a solution and valuing a criterion. The chords connected to the arc 'links between objects' are highlighted in color.

online

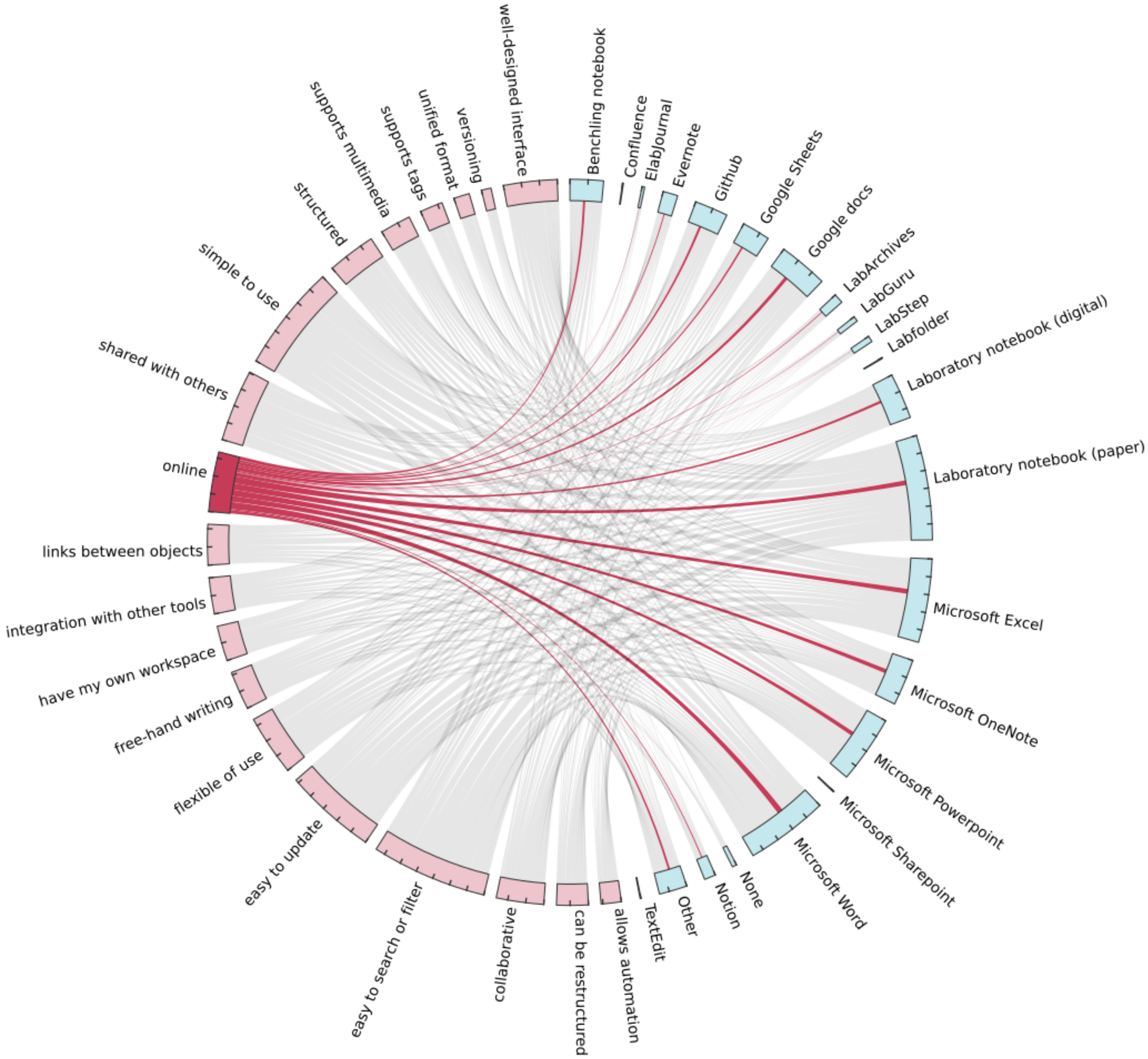


Note-taking solutions (blue arcs) vs desired features (red arcs). Arc length depicts the number of respondents, with ticks denoting 20 respondents. Link thickness shows the fraction of respondents using a solution and valuing a criterion. The chords connected to the arc 'online' are highlighted in color.

shared with others

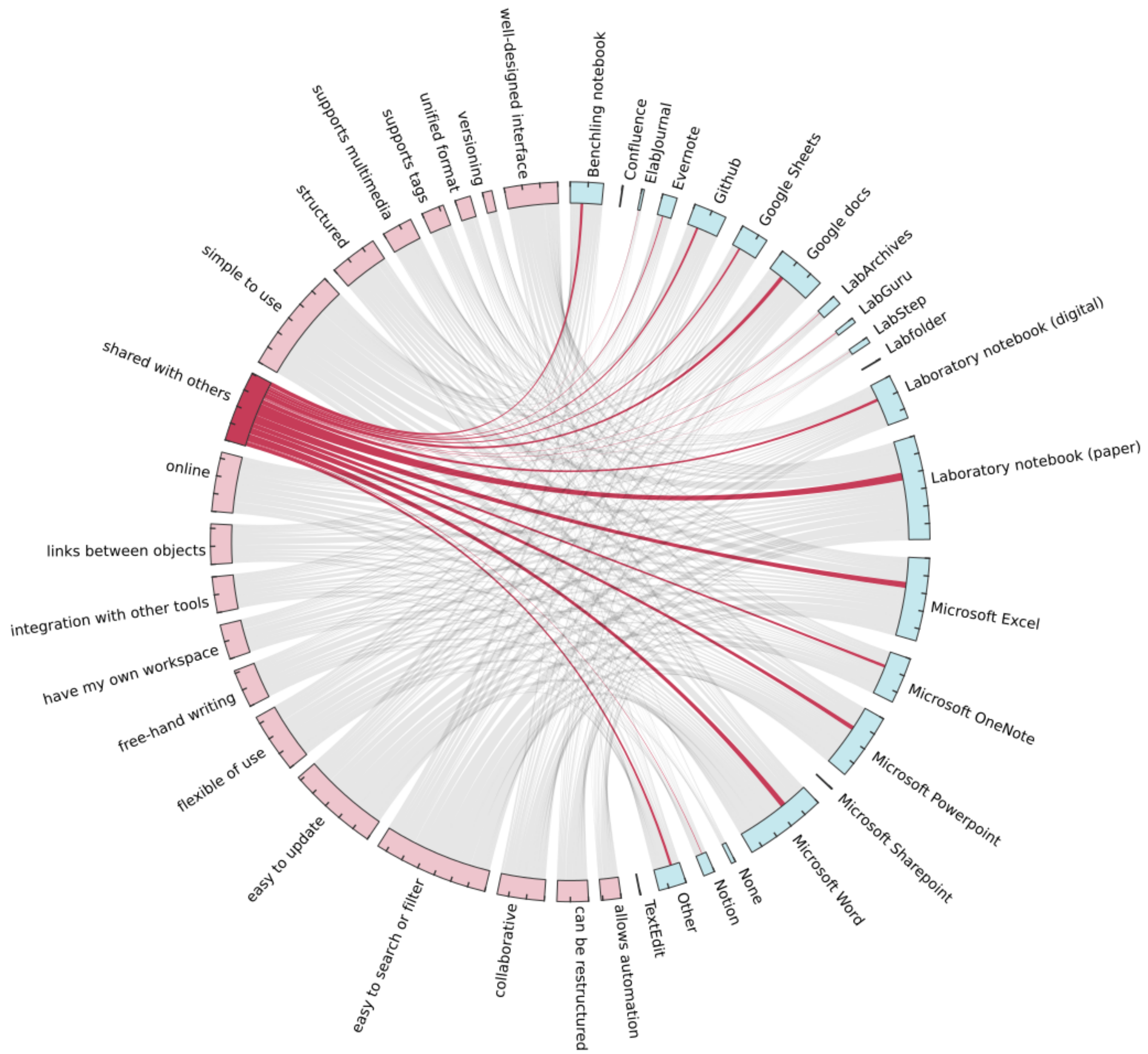


Note-taking solutions (blue arcs) vs desired features (red arcs). Arc length depicts the number of respondents, with ticks denoting 20 respondents. Link thickness shows the fraction of respondents using a solution and valuing a criterion. The chords connected to the arc 'shared with others' are highlighted in color.

simple to use

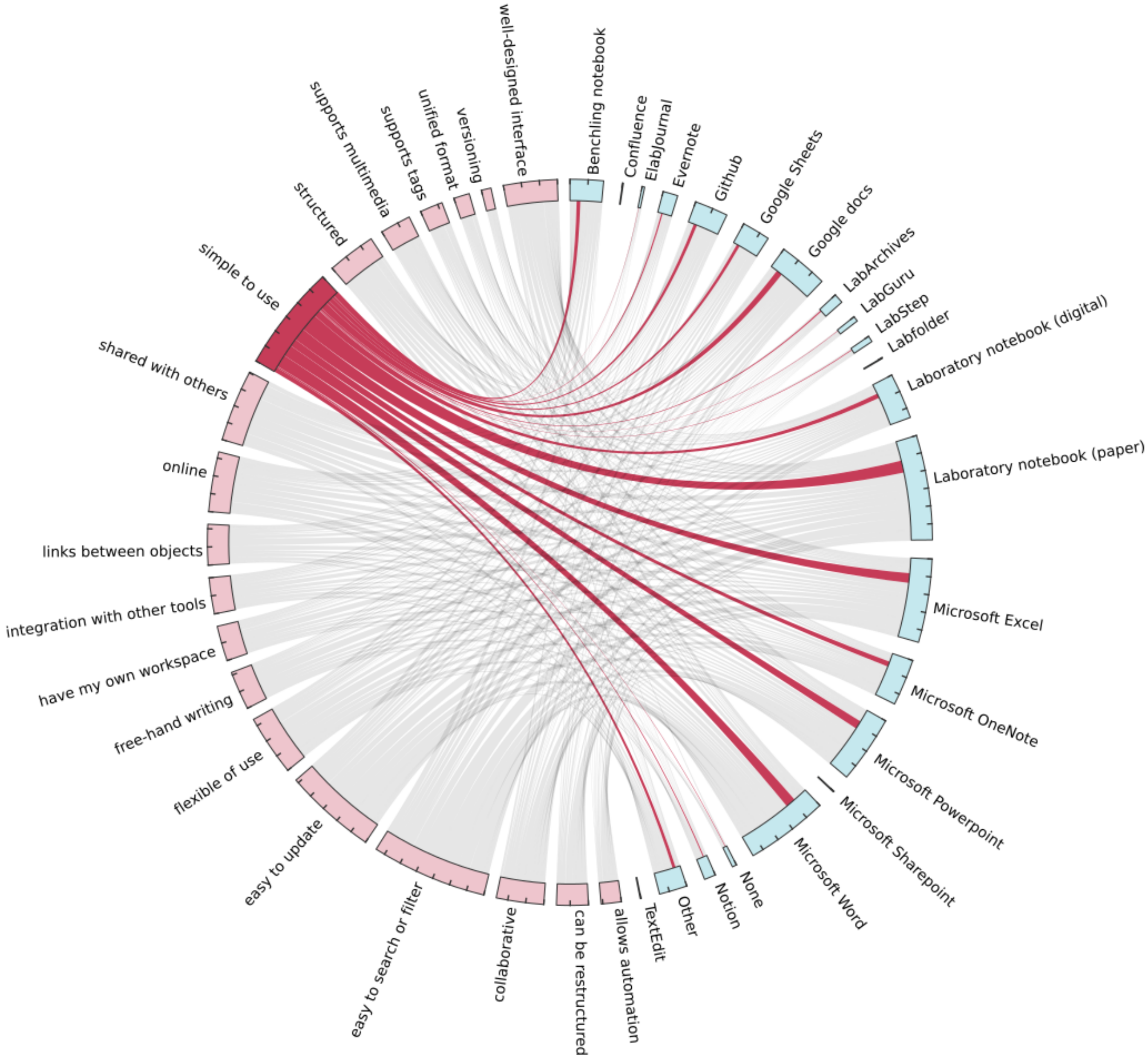


Note-taking solutions (blue arcs) vs desired features (red arcs). Arc length depicts the number of respondents, with ticks denoting 20 respondents. Link thickness shows the fraction of respondents using a solution and valuing a criterion. The chords connected to the arc 'simple to use' are highlighted in color.

structured

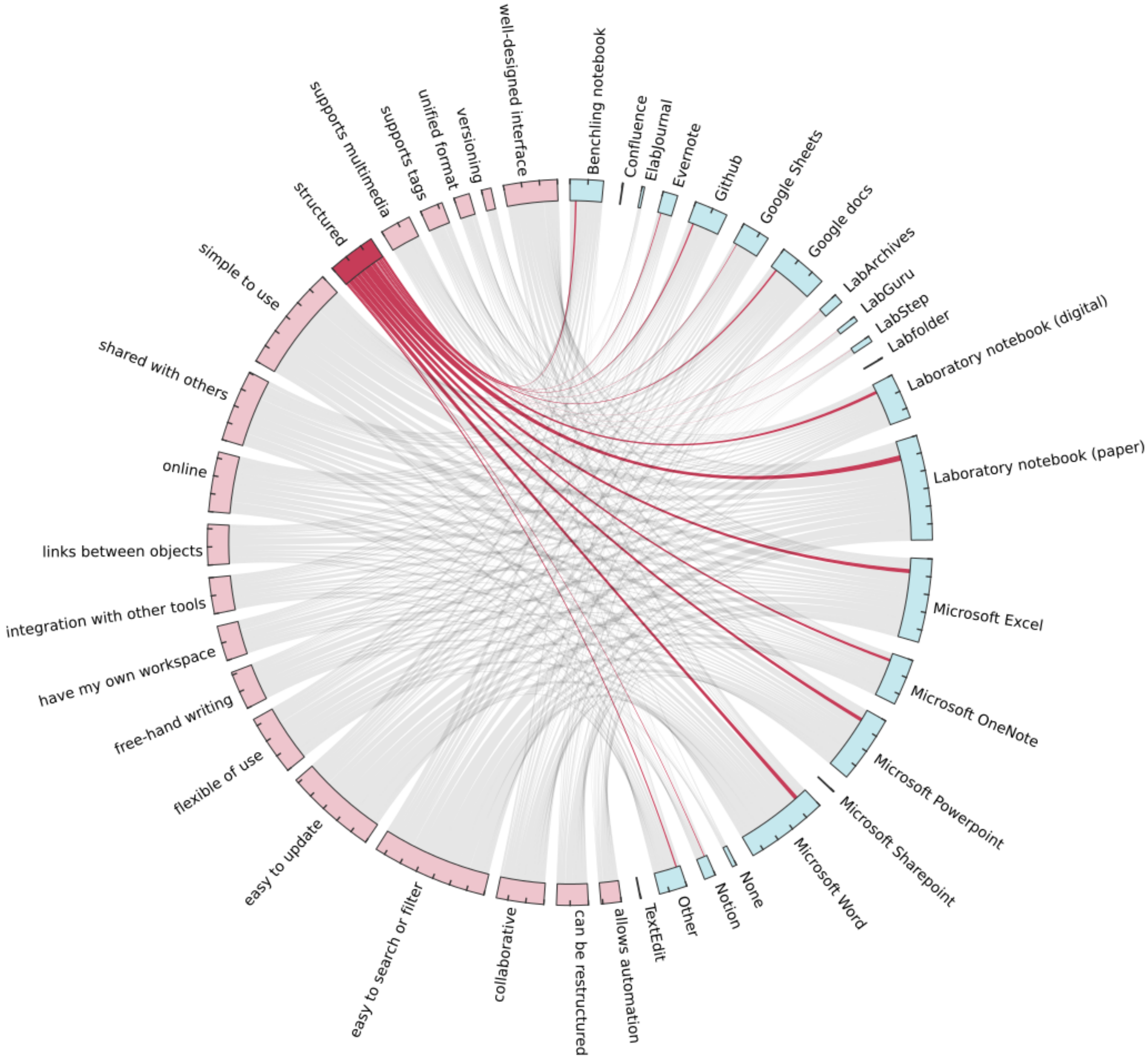


Note-taking solutions (blue arcs) vs desired features (red arcs). Arc length depicts the number of respondents, with ticks denoting 20 respondents. Link thickness shows the fraction of respondents using a solution and valuing a criterion. The chords connected to the arc 'structured' are highlighted in color.

supports multimedia

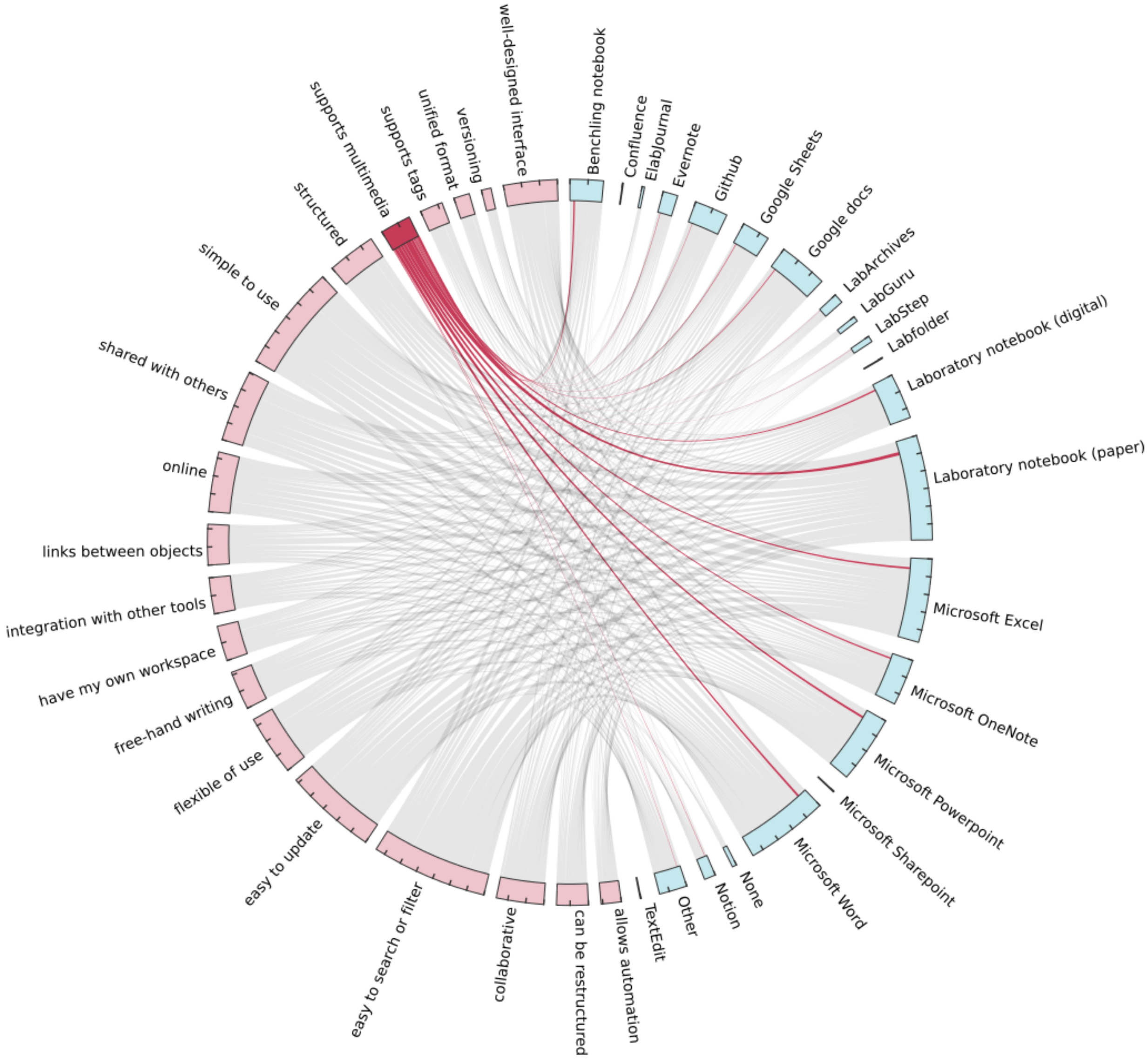


Note-taking solutions (blue arcs) vs desired features (red arcs). Arc length depicts the number of respondents, with ticks denoting 20 respondents. Link thickness shows the fraction of respondents using a solution and valuing a criterion. The chords connected to the arc 'supports multimedia' are highlighted in color.

supports tags

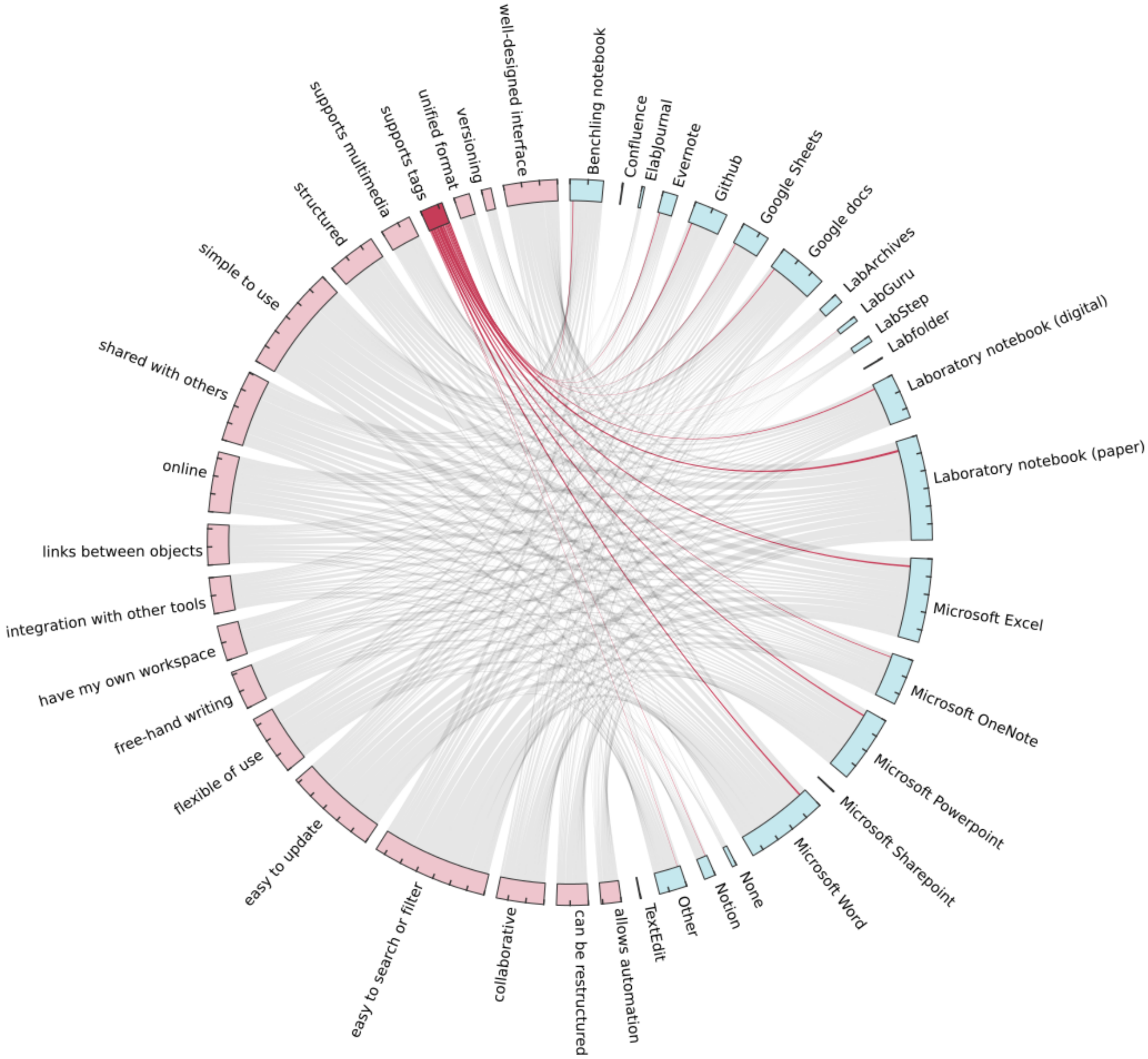


Note-taking solutions (blue arcs) vs desired features (red arcs). Arc length depicts the number of respondents, with ticks denoting 20 respondents. Link thickness shows the fraction of respondents using a solution and valuing a criterion. The chords connected to the arc 'supports tags' are highlighted in color.

unified format

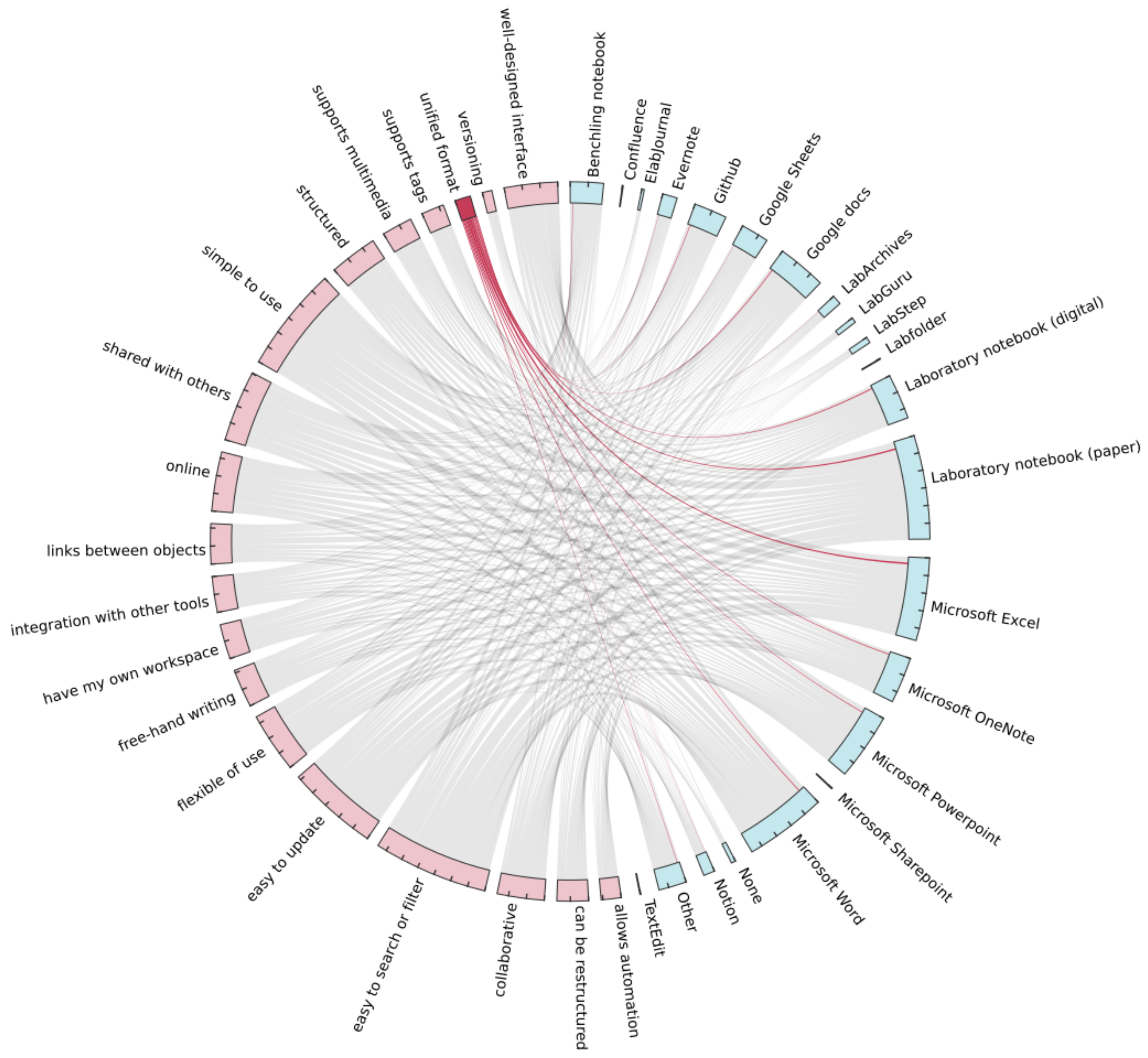


Note-taking solutions (blue arcs) vs desired features (red arcs). Arc length depicts the number of respondents, with ticks denoting 20 respondents. Link thickness shows the fraction of respondents using a solution and valuing a criterion. The chords connected to the arc 'unified format' are highlighted in color.

versioning

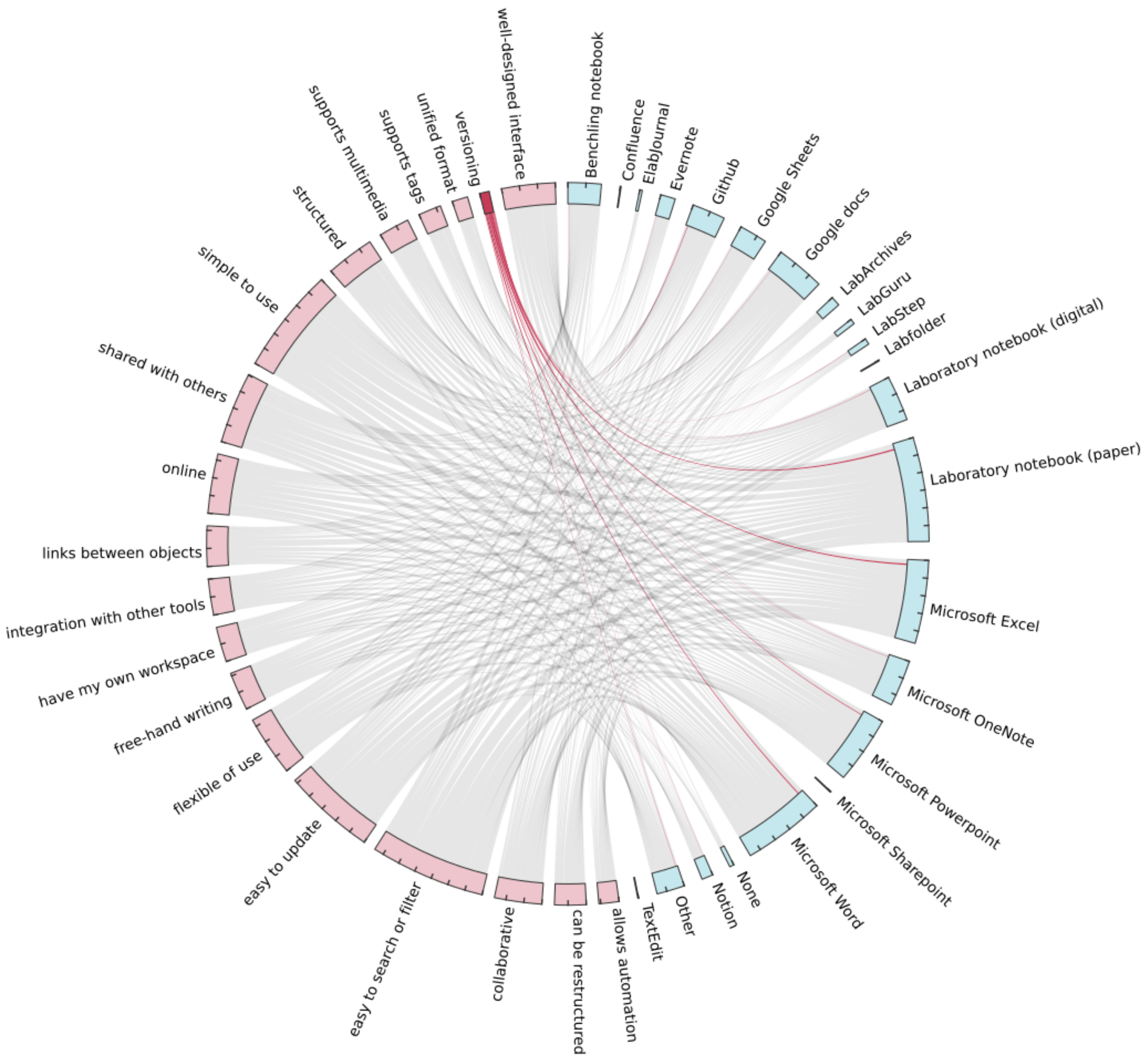


Note-taking solutions (blue arcs) vs desired features (red arcs). Arc length depicts the number of respondents, with ticks denoting 20 respondents. Link thickness shows the fraction of respondents using a solution and valuing a criterion. The chords connected to the arc 'versioning' are highlighted in color.

well-designed interface

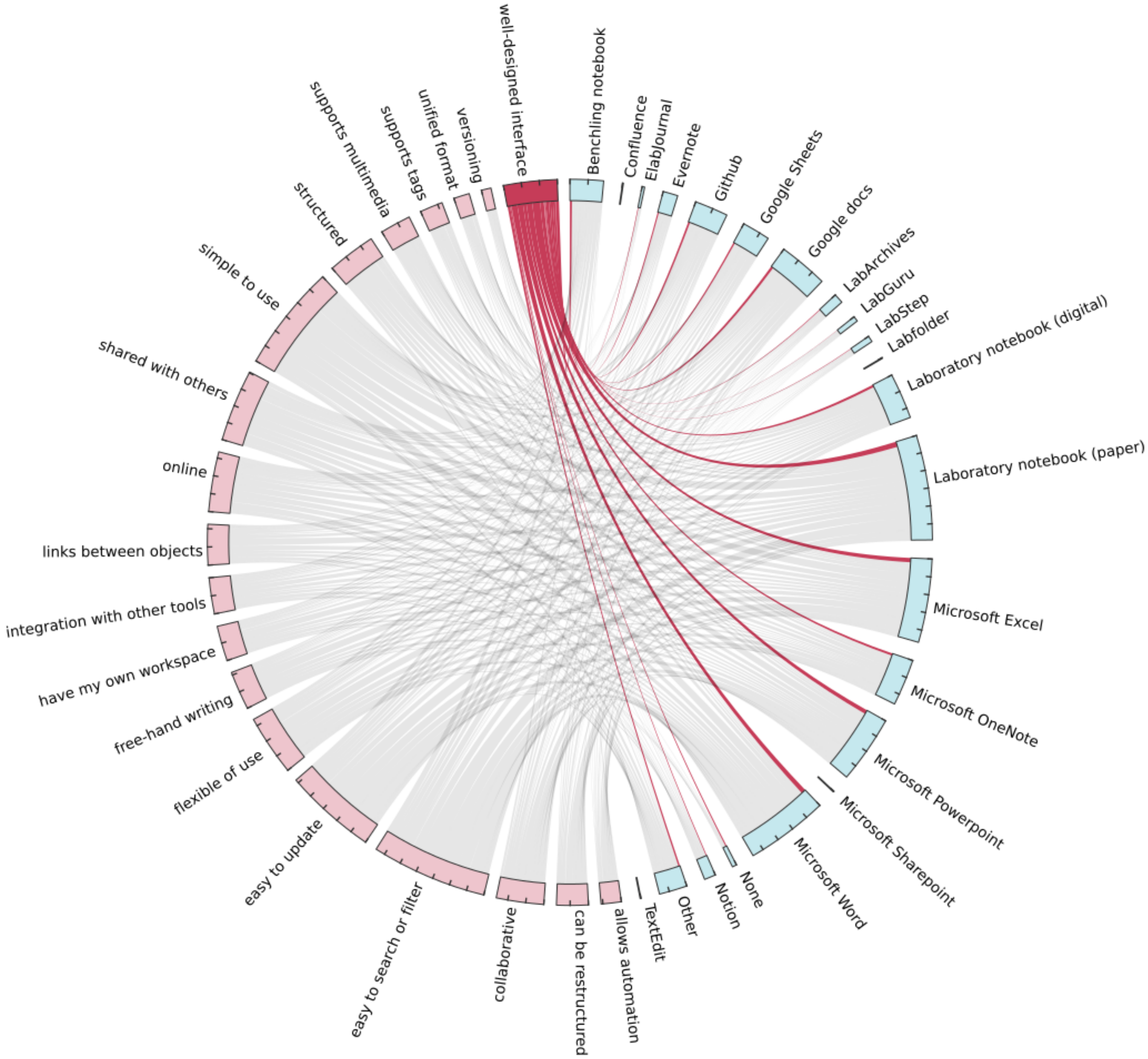


Note-taking solutions (blue arcs) vs desired features (red arcs). Arc length depicts the number of respondents, with ticks denoting 20 respondents. Link thickness shows the fraction of respondents using a solution and valuing a criterion. The chords connected to the arc 'well-designed interface' are highlighted in color.
