## Supplementary Data 1 for "A platform for lab management, note-keeping and automation": Supplementary Data 1 - Sample management.pdf

A-tune

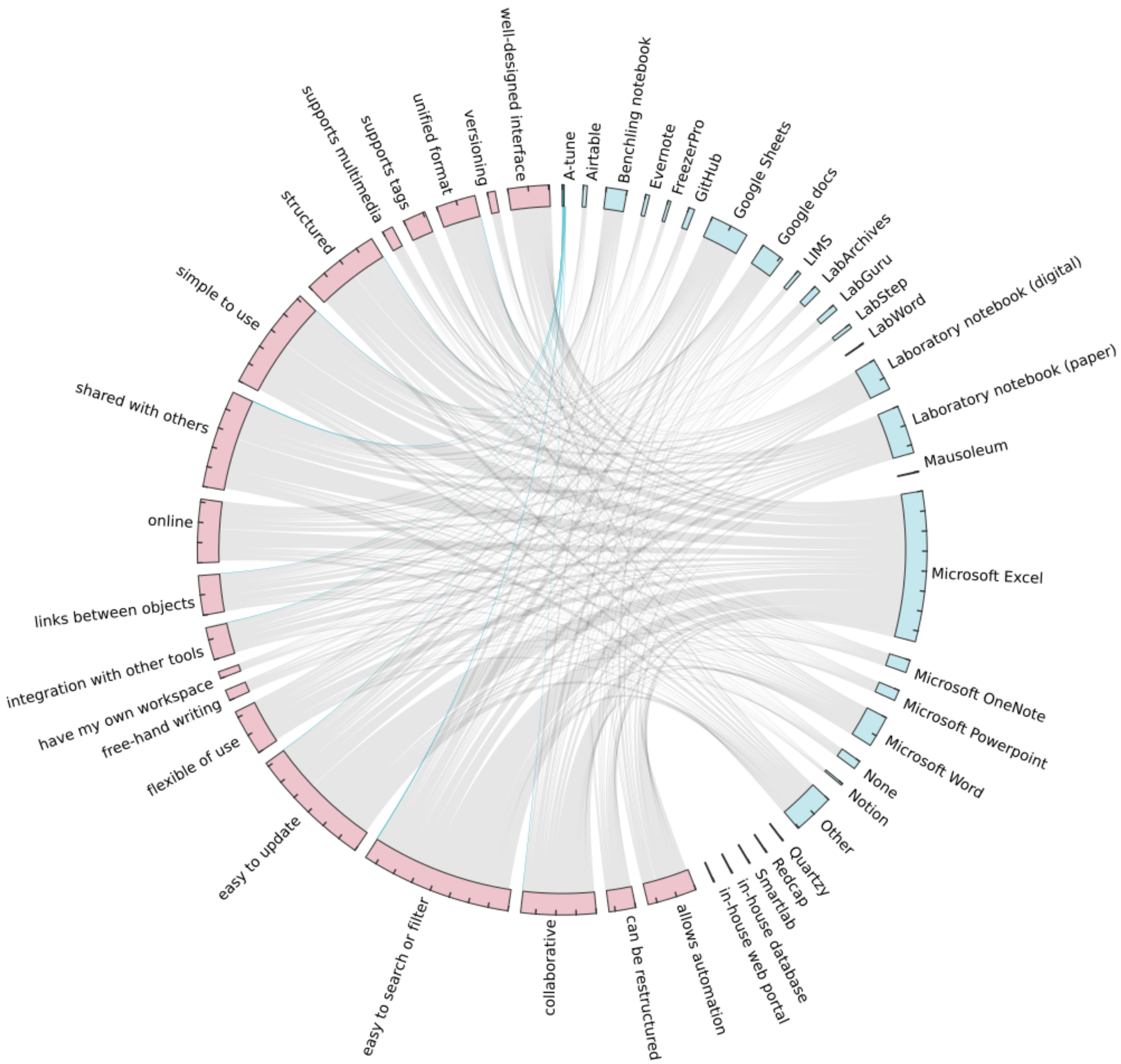


Sample-management solutions (blue arcs) vs desired features (red arcs). Arc length depicts the number of respondents, with ticks denoting 20 respondents. Link thickness shows the fraction of respondents using a solution and valuing a criterion. The chords connected to the arc 'A-tune' are highlighted in color.'

Airtable

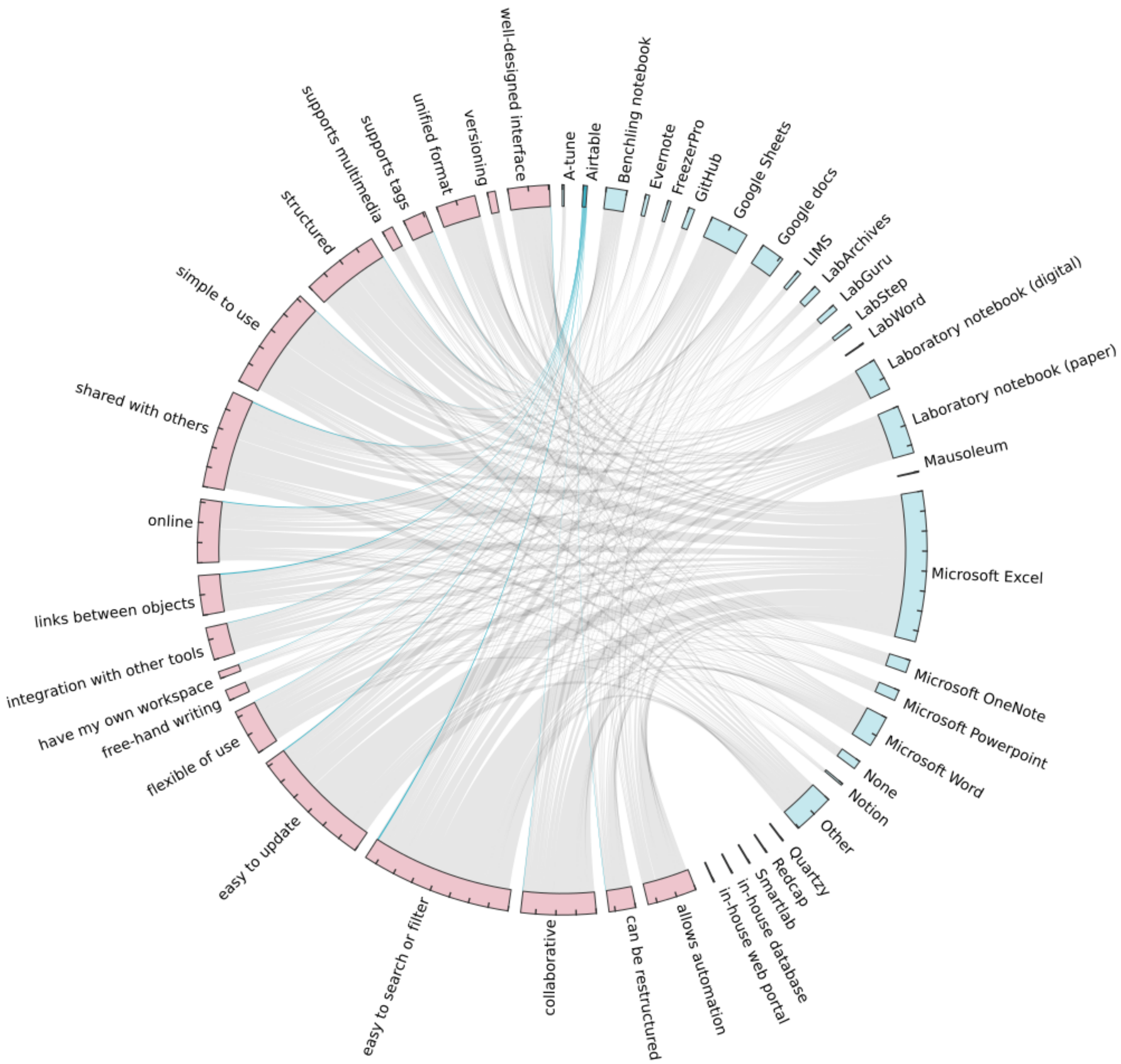


Sample-management solutions (blue arcs) vs desired features (red arcs). Arc length depicts the number of respondents, with ticks denoting 20 respondents. Link thickness shows the fraction of respondents using a solution and valuing a criterion. The chords connected to the arc 'Airtable' are highlighted in color.'

Benchling notebook

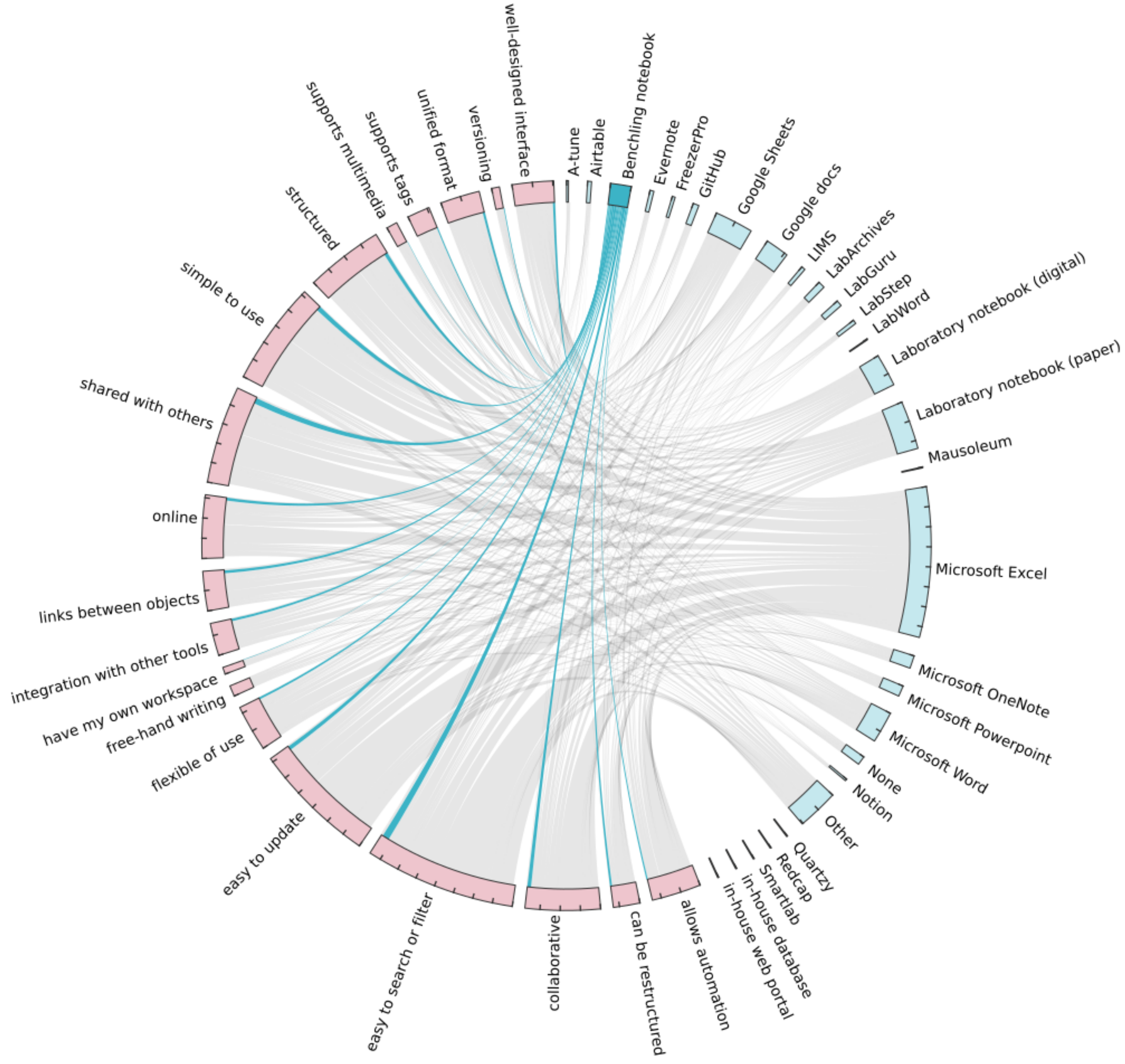


Sample-management solutions (blue arcs) vs desired features (red arcs). Arc length depicts the number of respondents, with ticks denoting 20 respondents. Link thickness shows the fraction of respondents using a solution and valuing a criterion. The chords connected to the arc 'Benchling notebook' are highlighted in color.'

Evernote

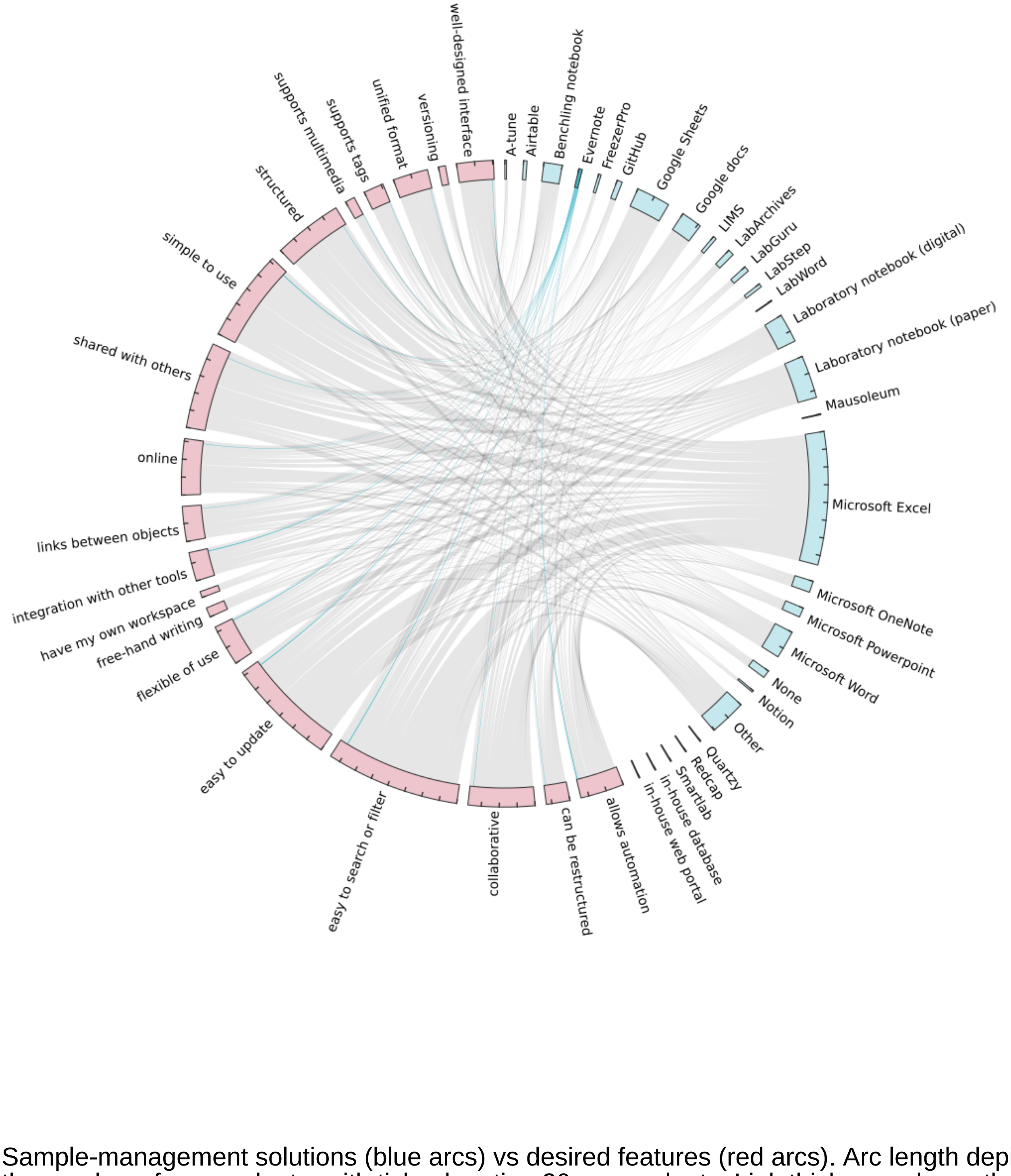


Sample-management solutions (blue arcs) vs desired features (red arcs). Arc length depicts the number of respondents, with ticks denoting 20 respondents. Link thickness shows the fraction of respondents using a solution and valuing a criterion. The chords connected to the arc 'Evernote' are highlighted in color.'

FreezerPro

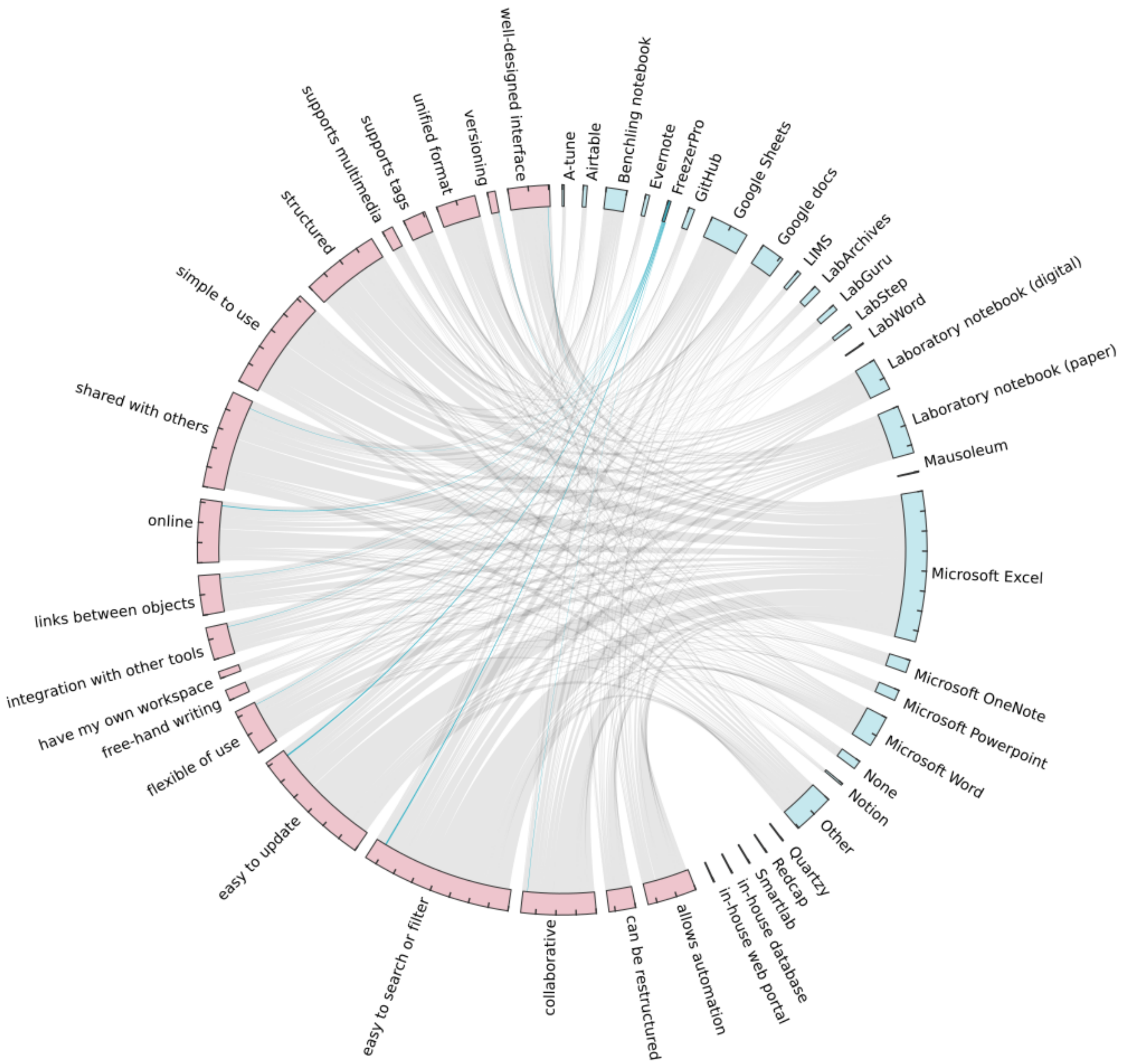


Sample-management solutions (blue arcs) vs desired features (red arcs). Arc length depicts the number of respondents, with ticks denoting 20 respondents. Link thickness shows the fraction of respondents using a solution and valuing a criterion. The chords connected to the arc 'FreezerPro' are highlighted in color.'

GitHub

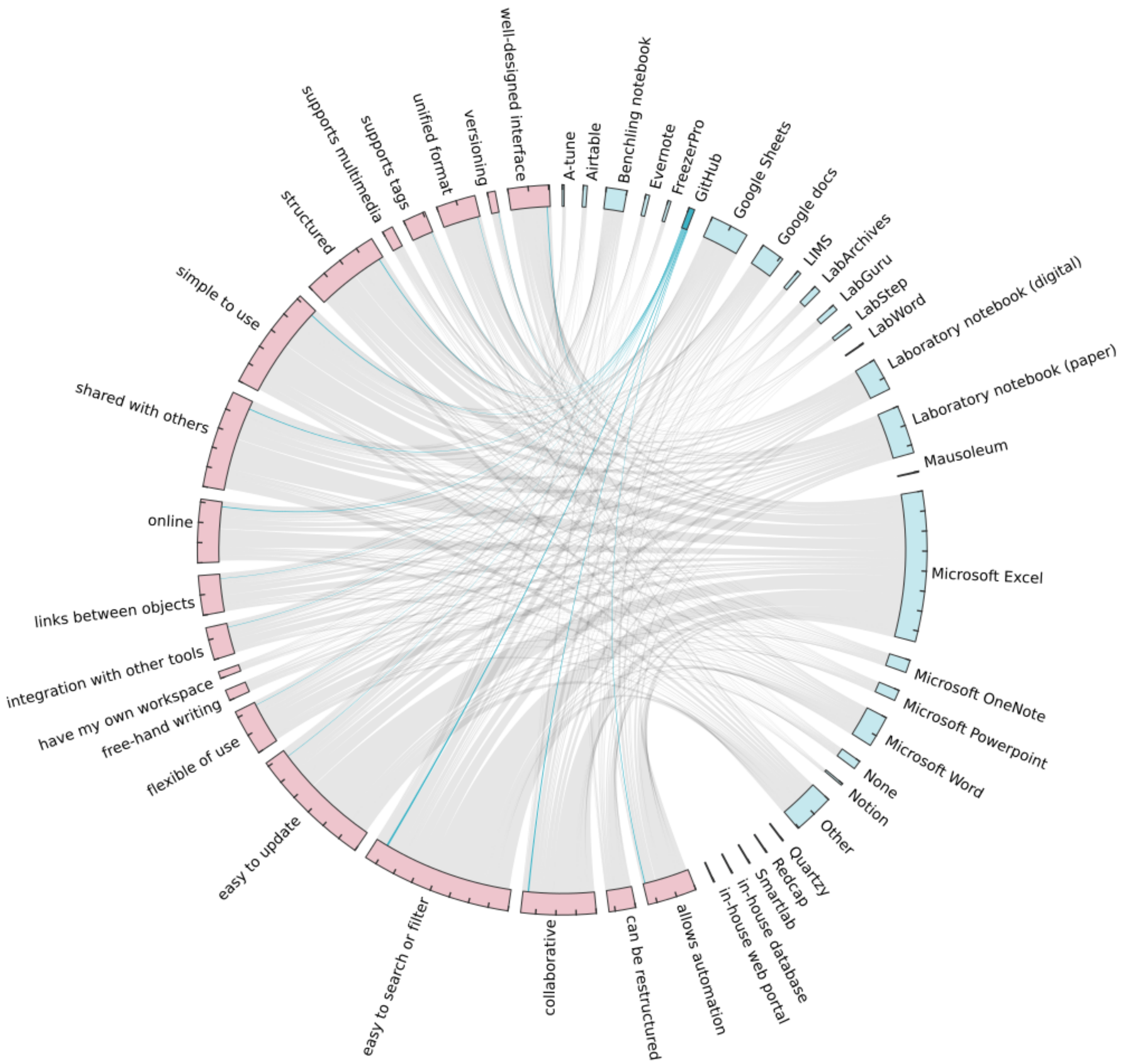


Sample-management solutions (blue arcs) vs desired features (red arcs). Arc length depicts the number of respondents, with ticks denoting 20 respondents. Link thickness shows the fraction of respondents using a solution and valuing a criterion. The chords connected to the arc 'GitHub' are highlighted in color.'

Google Sheets

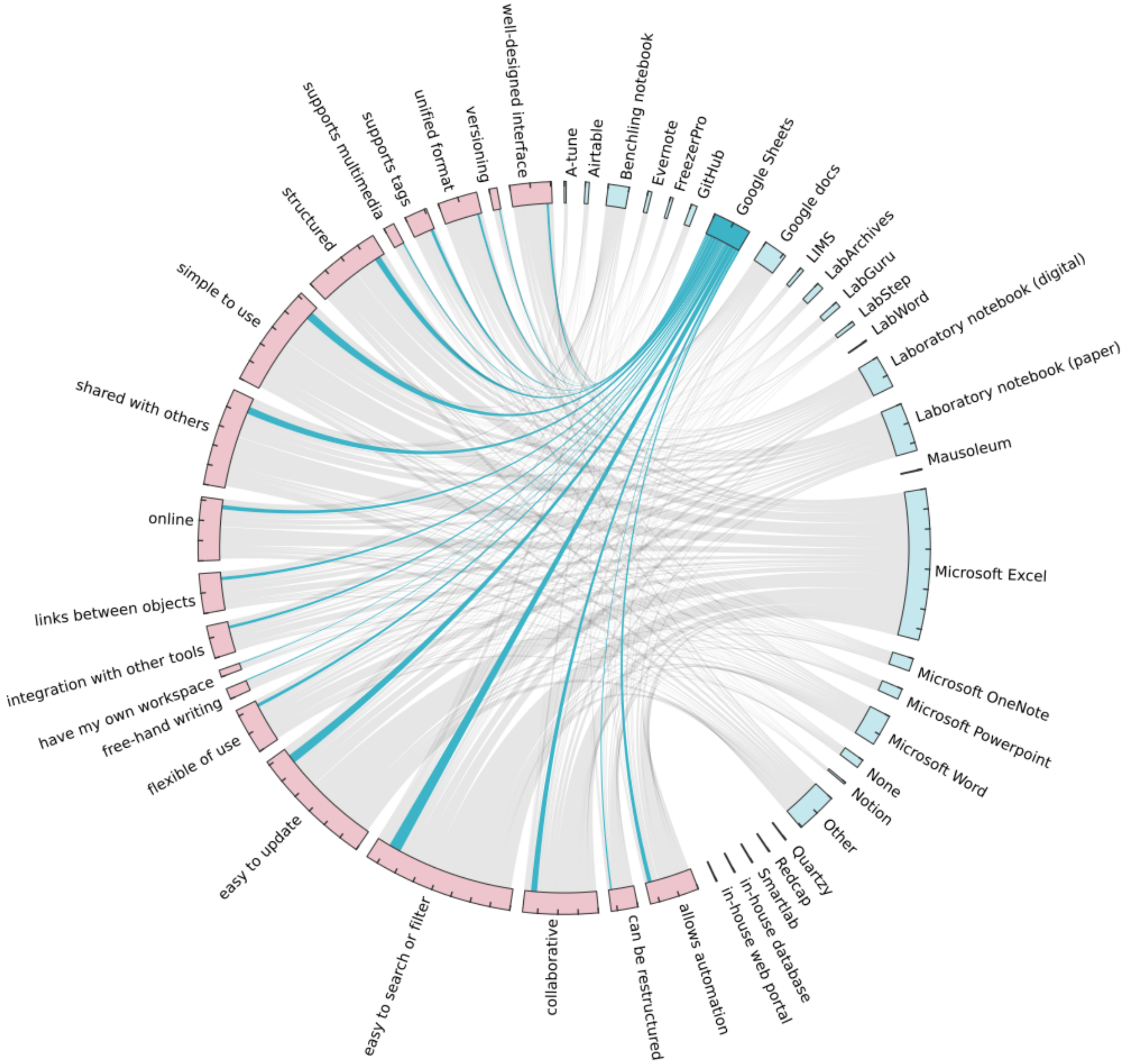


Sample-management solutions (blue arcs) vs desired features (red arcs). Arc length depicts the number of respondents, with ticks denoting 20 respondents. Link thickness shows the fraction of respondents using a solution and valuing a criterion. The chords connected to the arc 'Google Sheets' are highlighted in color.'

Google docs

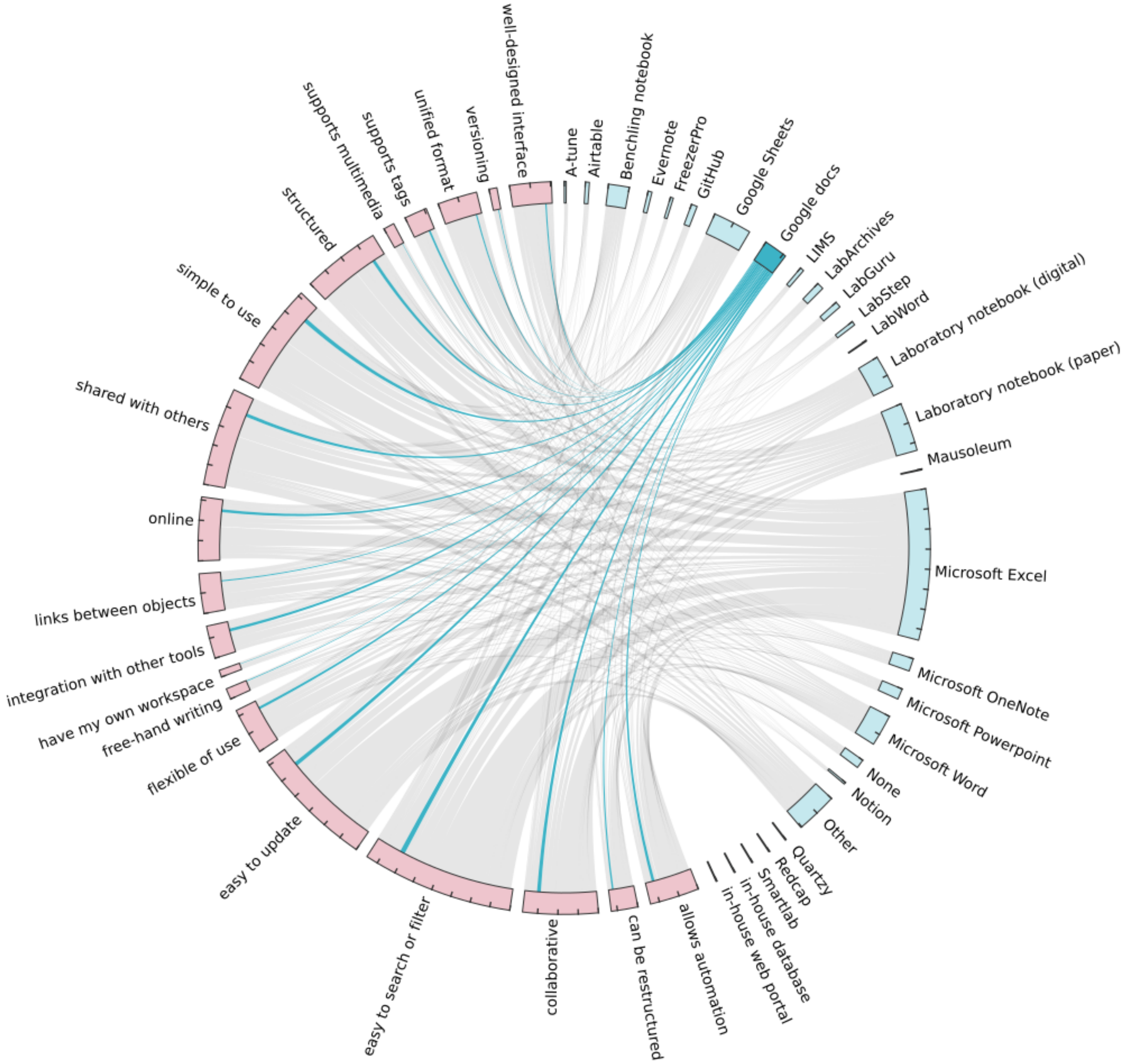


Sample-management solutions (blue arcs) vs desired features (red arcs). Arc length depicts the number of respondents, with ticks denoting 20 respondents. Link thickness shows the fraction of respondents using a solution and valuing a criterion. The chords connected to the arc 'Google docs' are highlighted in color.'

LIMS

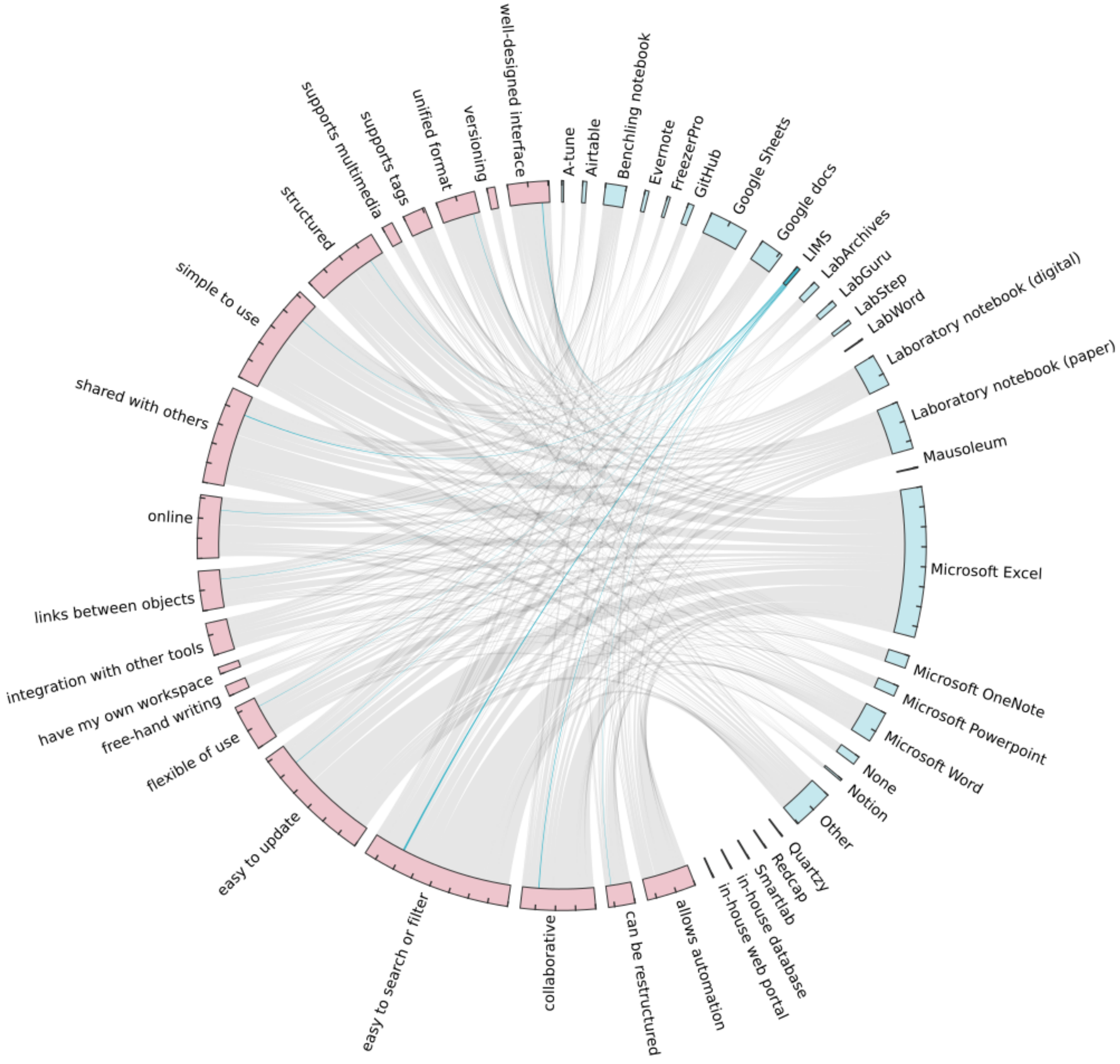


Sample-management solutions (blue arcs) vs desired features (red arcs). Arc length depicts the number of respondents, with ticks denoting 20 respondents. Link thickness shows the fraction of respondents using a solution and valuing a criterion. The chords connected to the arc 'LIMS' are highlighted in color.'

LabArchives

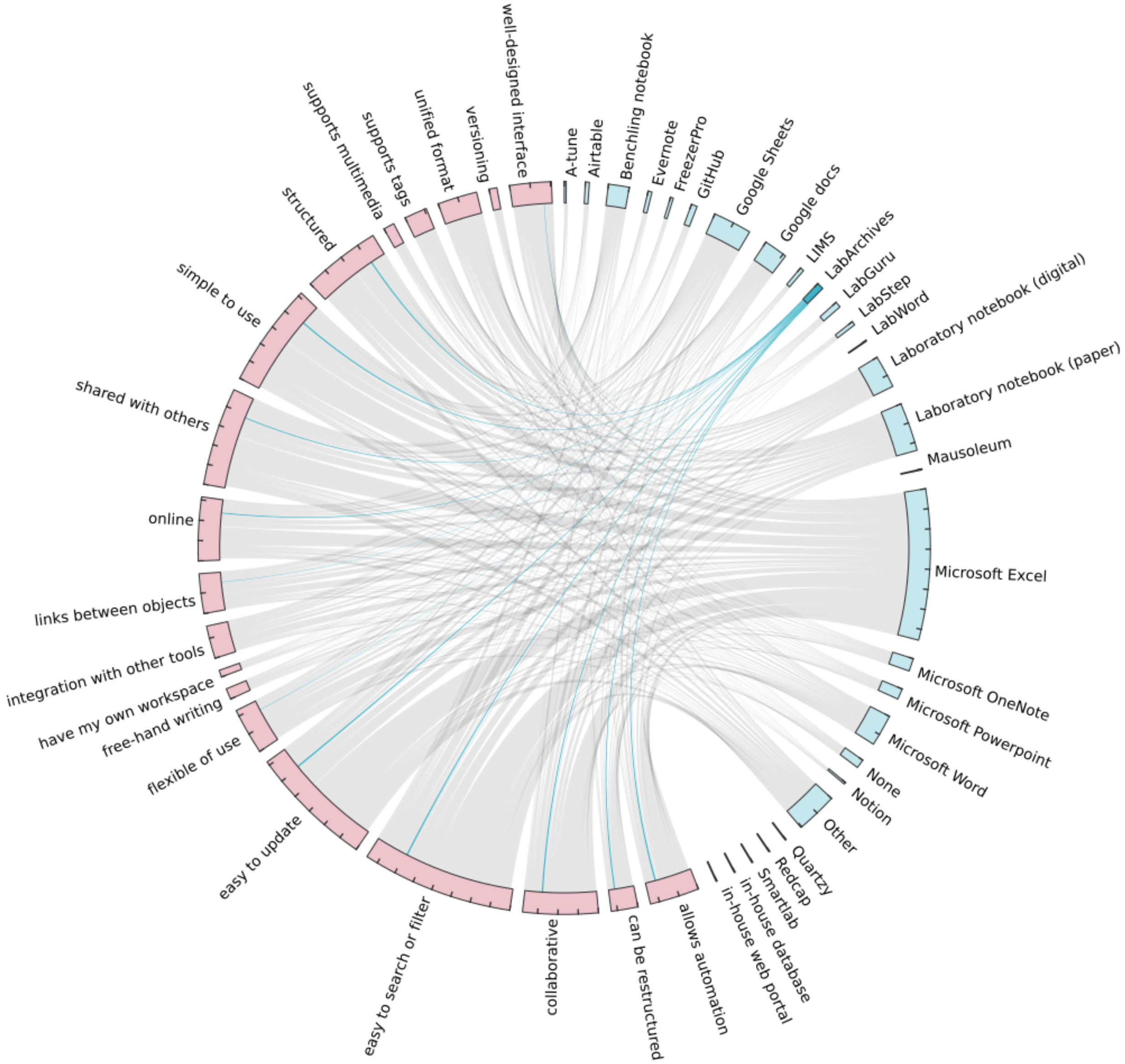


Sample-management solutions (blue arcs) vs desired features (red arcs). Arc length depicts the number of respondents, with ticks denoting 20 respondents. Link thickness shows the fraction of respondents using a solution and valuing a criterion. The chords connected to the arc 'LabArchives' are highlighted in color.'

LabGuru

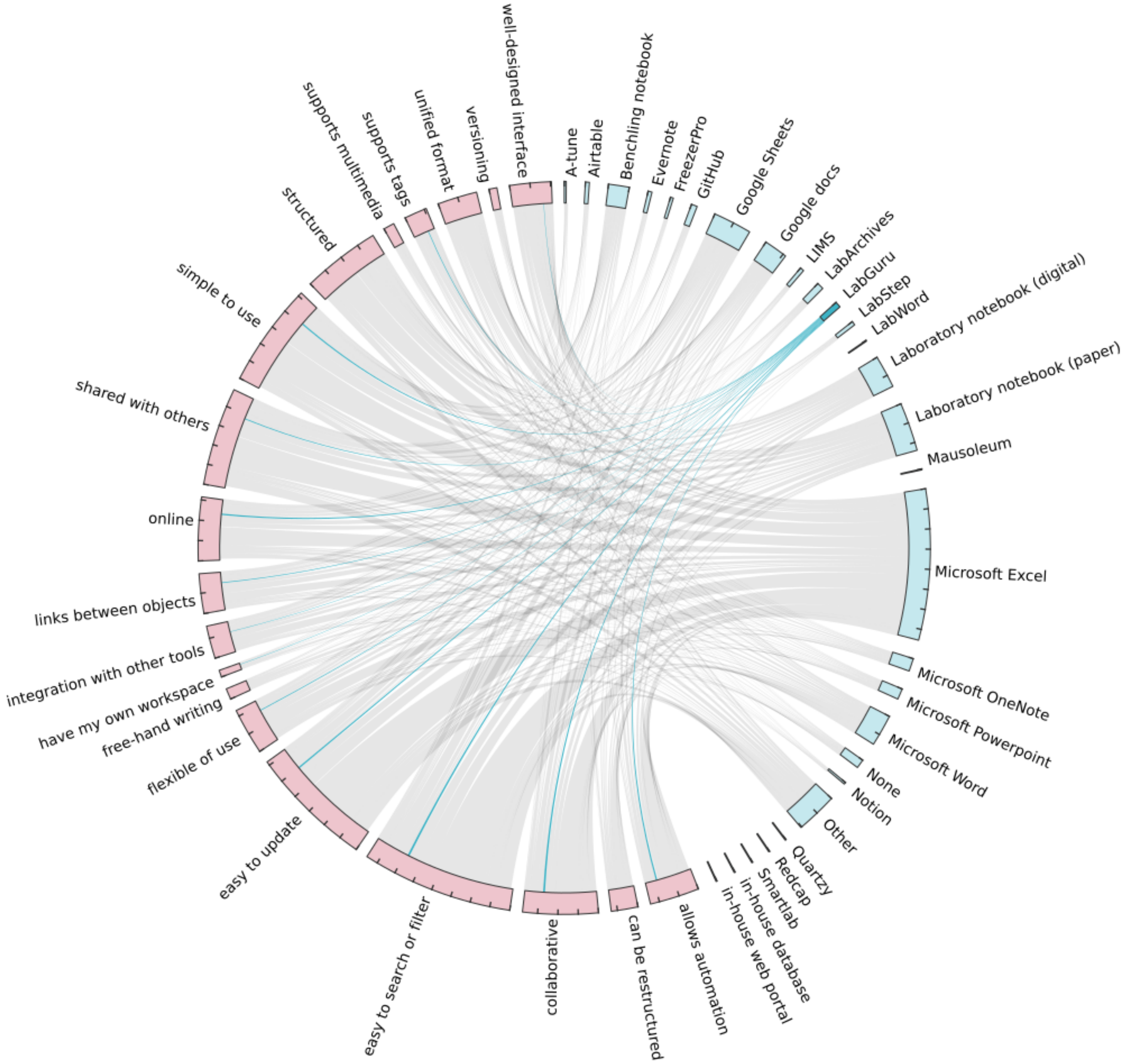


Sample-management solutions (blue arcs) vs desired features (red arcs). Arc length depicts the number of respondents, with ticks denoting 20 respondents. Link thickness shows the fraction of respondents using a solution and valuing a criterion. The chords connected to the arc 'LabGuru' are highlighted in color.'

LabStep

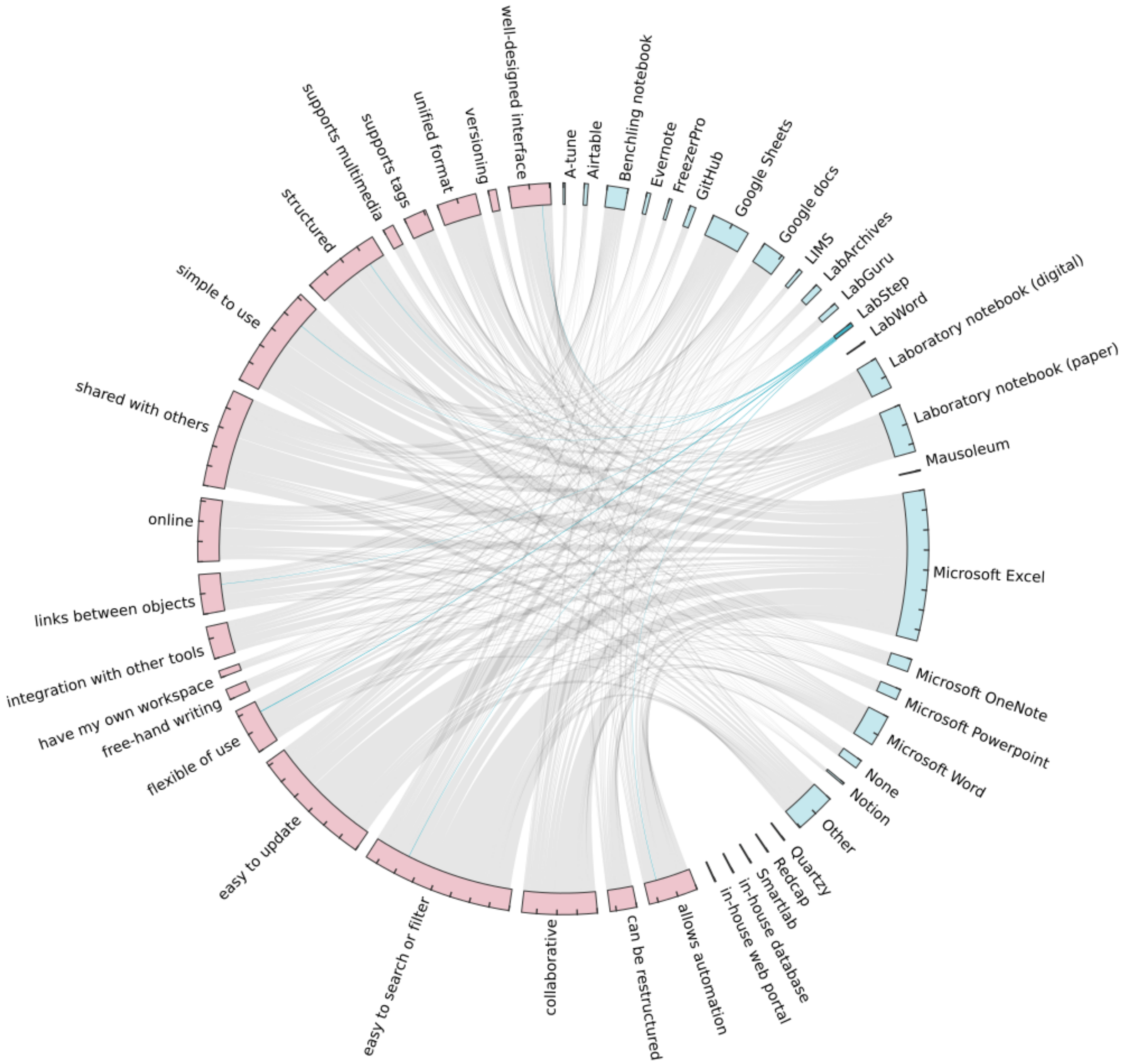


Sample-management solutions (blue arcs) vs desired features (red arcs). Arc length depicts the number of respondents, with ticks denoting 20 respondents. Link thickness shows the fraction of respondents using a solution and valuing a criterion. The chords connected to the arc 'LabStep' are highlighted in color.'

LabWord

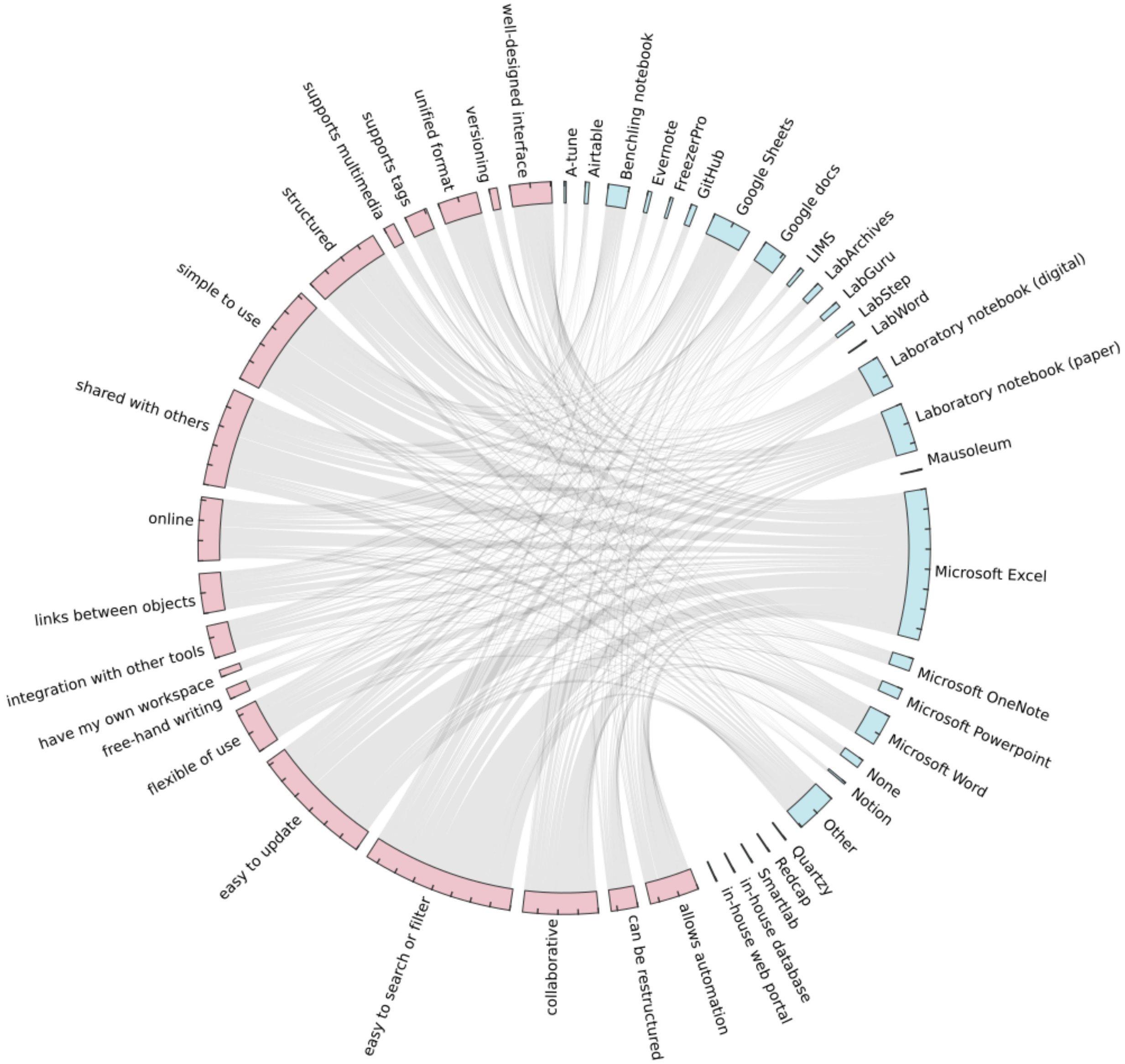


Sample-management solutions (blue arcs) vs desired features (red arcs). Arc length depicts the number of respondents, with ticks denoting 20 respondents. Link thickness shows the fraction of respondents using a solution and valuing a criterion. The chords connected to the arc 'LabWord' are highlighted in color.'

Laboratory notebook (digital)

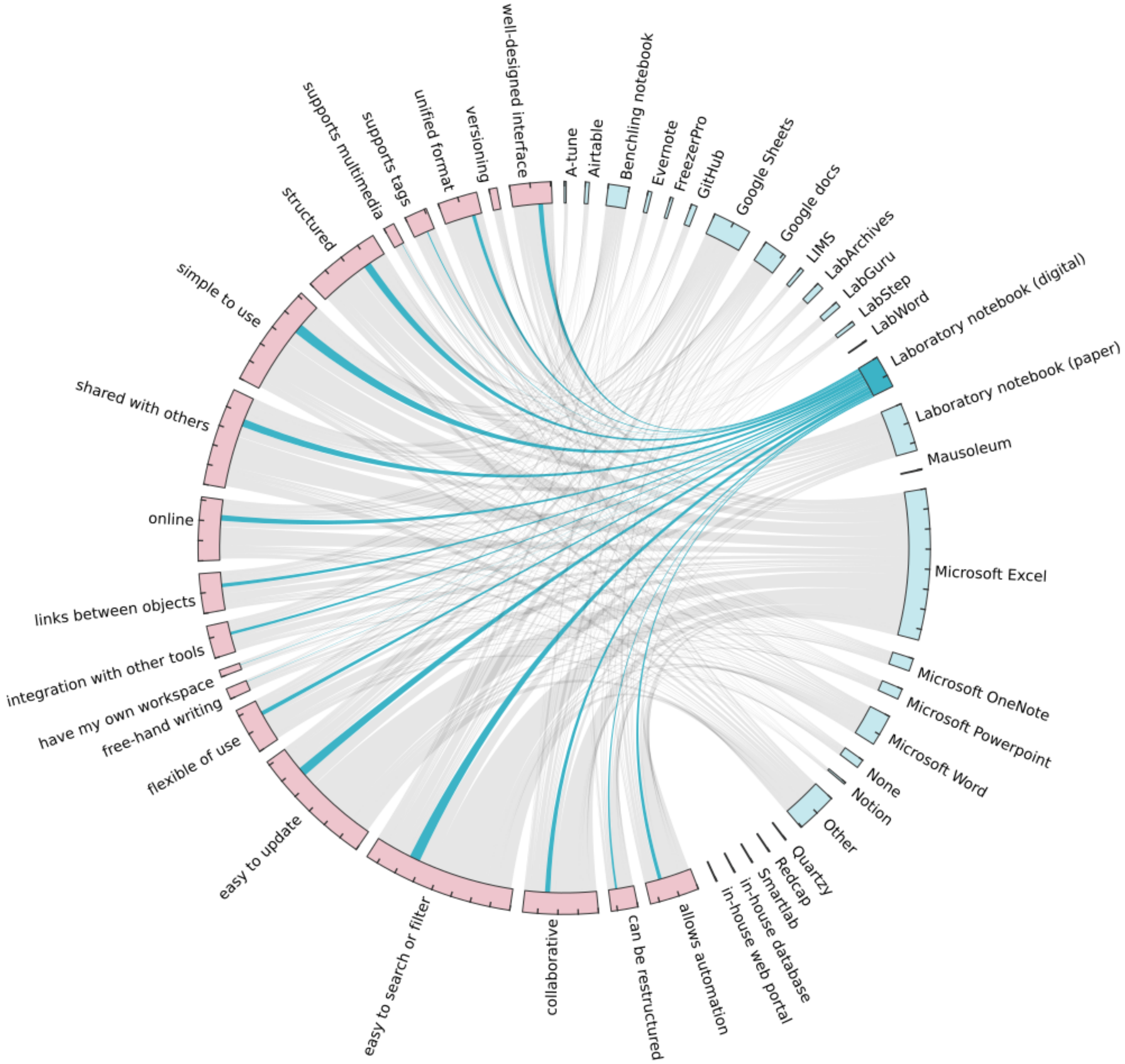


Sample-management solutions (blue arcs) vs desired features (red arcs). Arc length depicts the number of respondents, with ticks denoting 20 respondents. Link thickness shows the fraction of respondents using a solution and valuing a criterion. The chords connected to the arc 'Laboratory notebook (digital)' are highlighted in color.'

Laboratory notebook (paper)

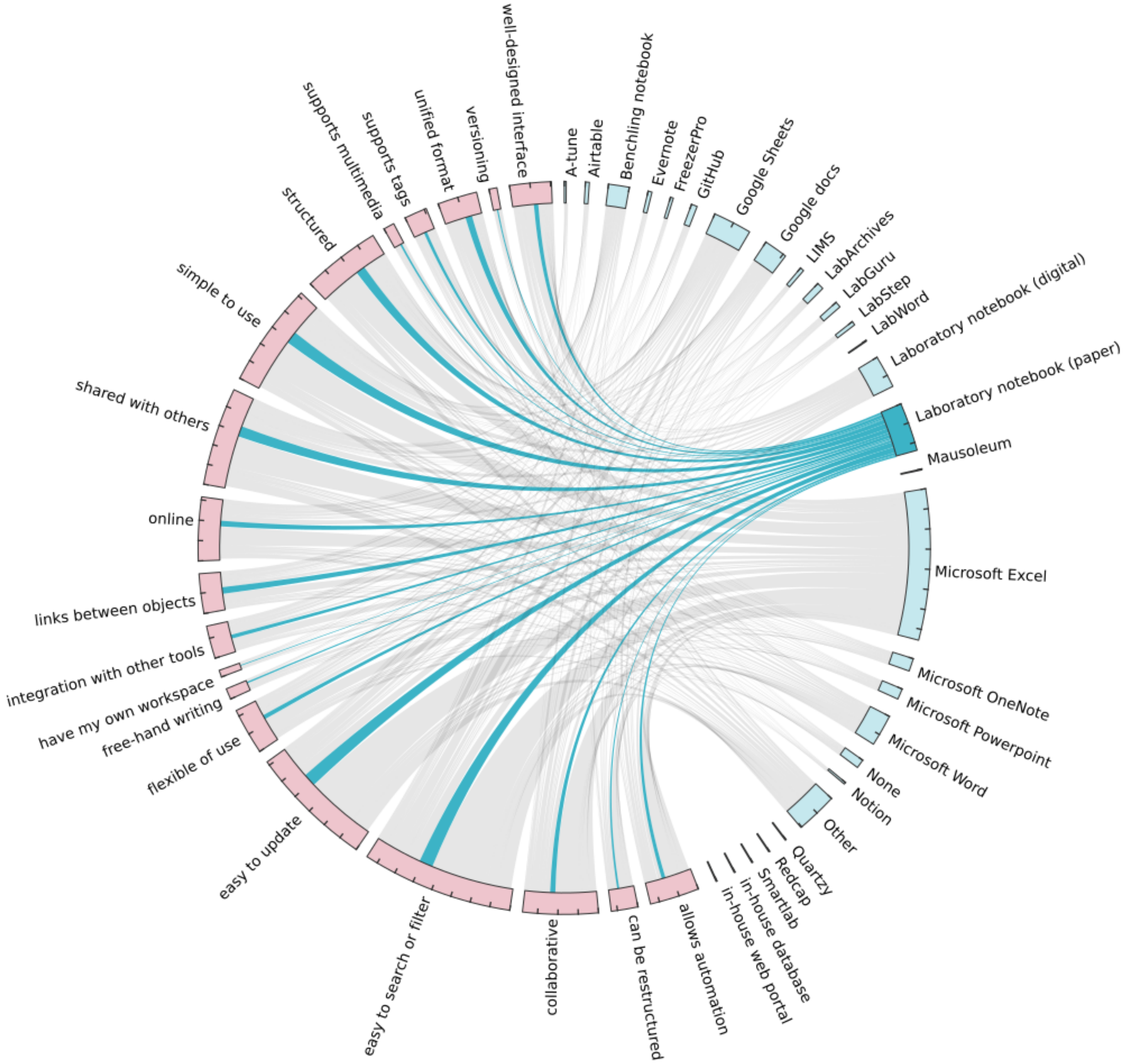


Sample-management solutions (blue arcs) vs desired features (red arcs). Arc length depicts the number of respondents, with ticks denoting 20 respondents. Link thickness shows the fraction of respondents using a solution and valuing a criterion. The chords connected to the arc 'Laboratory notebook (paper)' are highlighted in color.'

Mausoleum

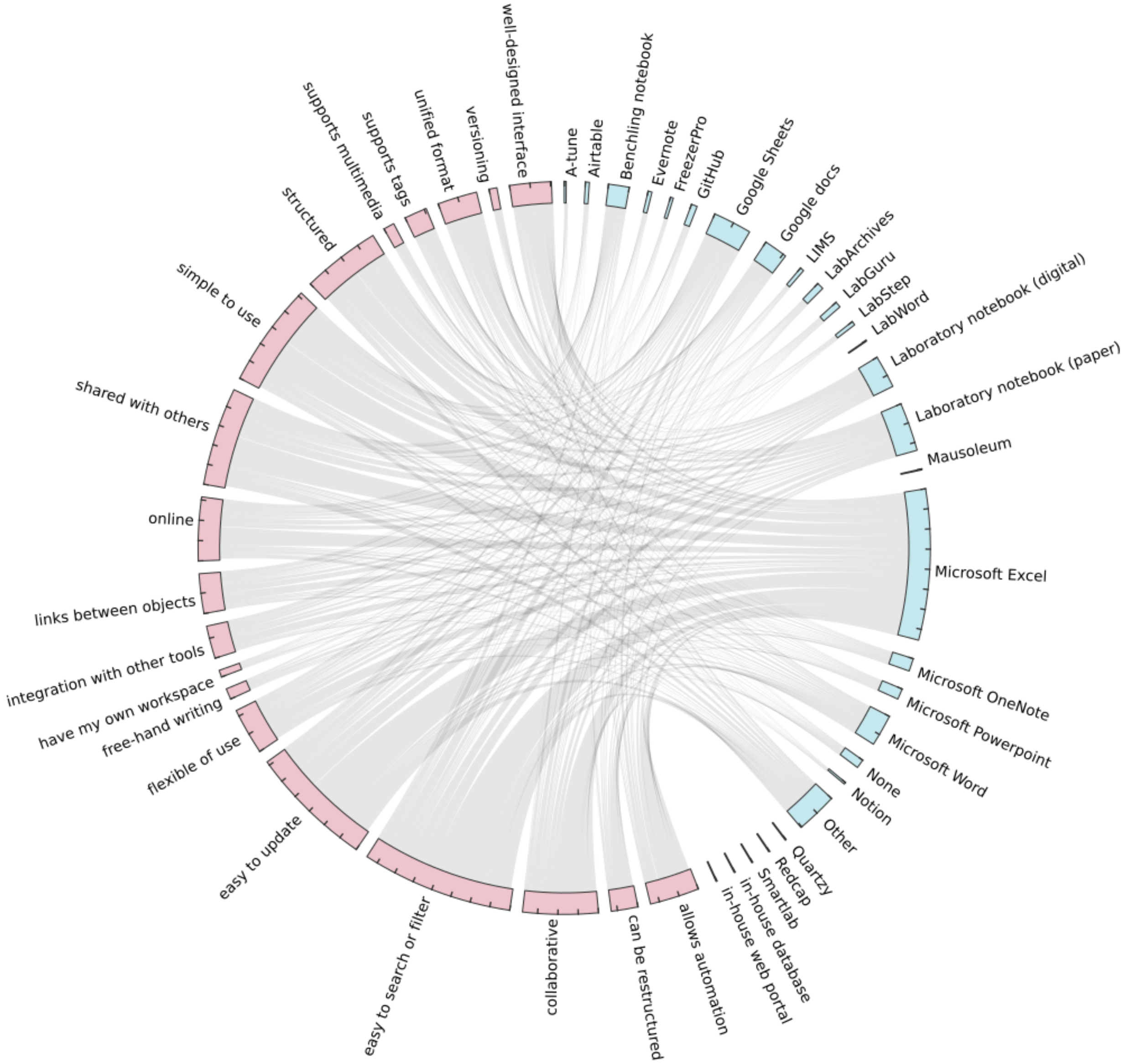


Sample-management solutions (blue arcs) vs desired features (red arcs). Arc length depicts the number of respondents, with ticks denoting 20 respondents. Link thickness shows the fraction of respondents using a solution and valuing a criterion. The chords connected to the arc 'Mausoleum' are highlighted in color.'

Microsoft Excel

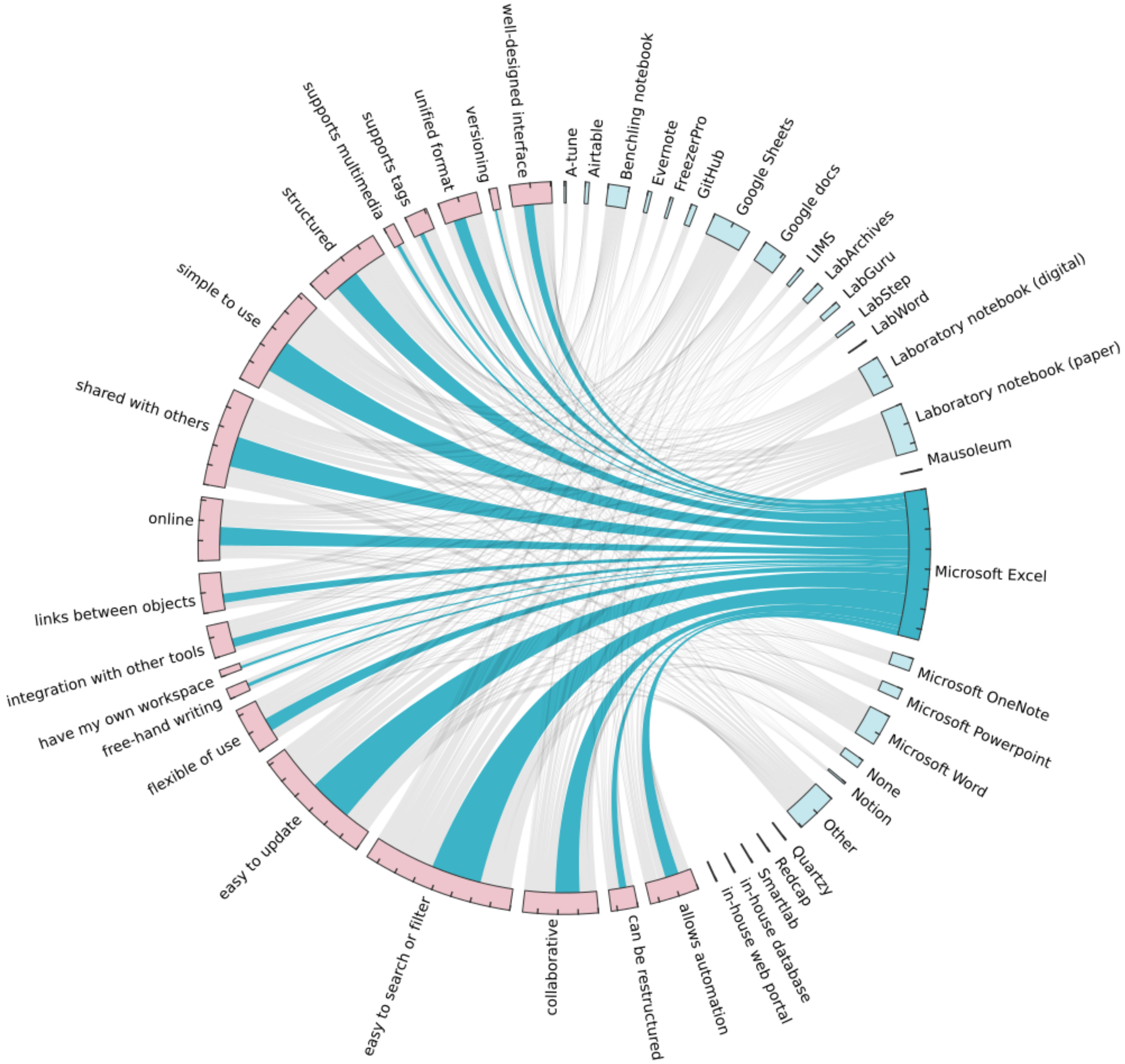


Sample-management solutions (blue arcs) vs desired features (red arcs). Arc length depicts the number of respondents, with ticks denoting 20 respondents. Link thickness shows the fraction of respondents using a solution and valuing a criterion. The chords connected to the arc 'Microsoft Excel' are highlighted in color.'

Microsoft OneNote

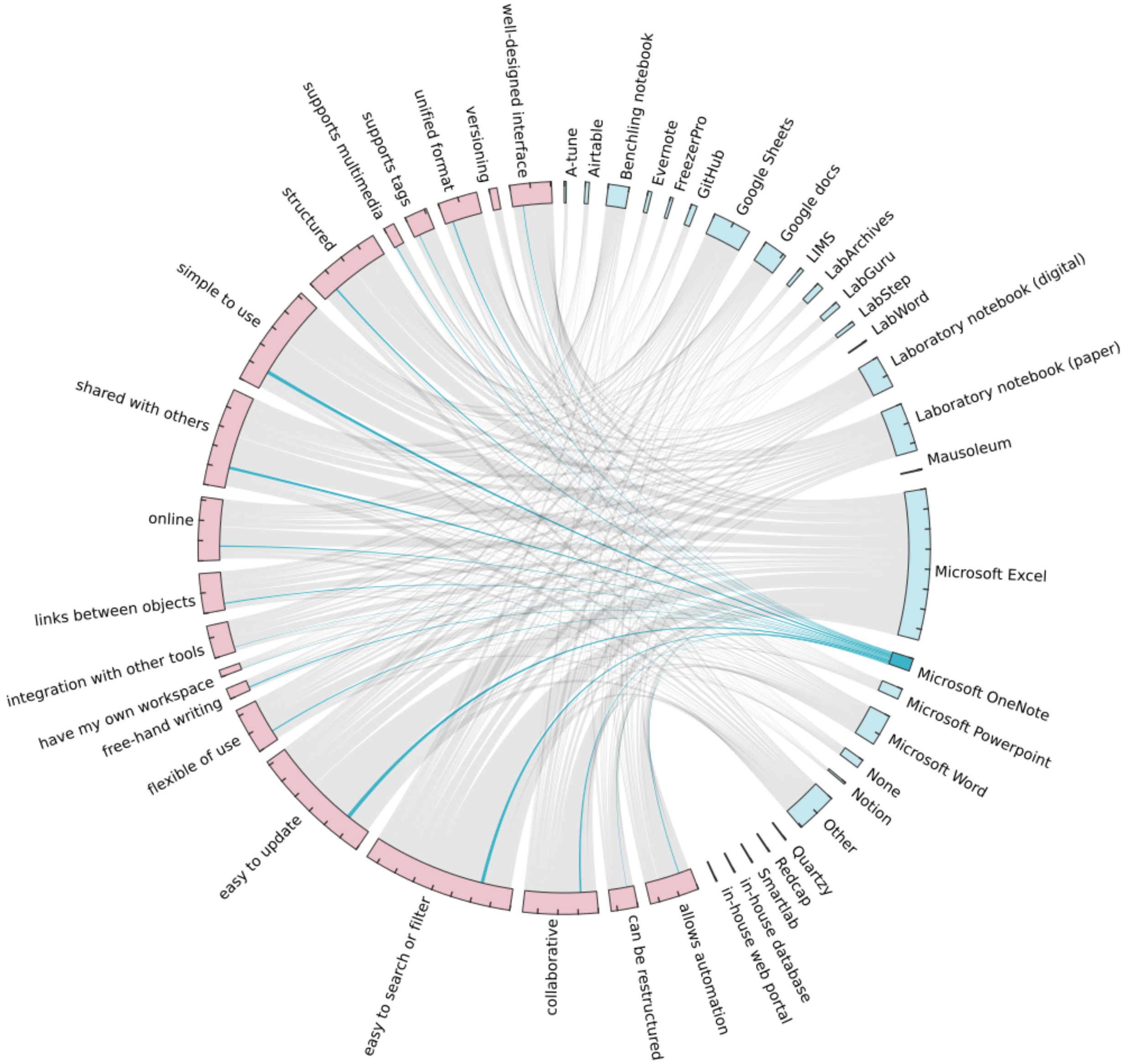


Sample-management solutions (blue arcs) vs desired features (red arcs). Arc length depicts the number of respondents, with ticks denoting 20 respondents. Link thickness shows the fraction of respondents using a solution and valuing a criterion. The chords connected to the arc 'Microsoft OneNote' are highlighted in color.'

Microsoft Powerpoint

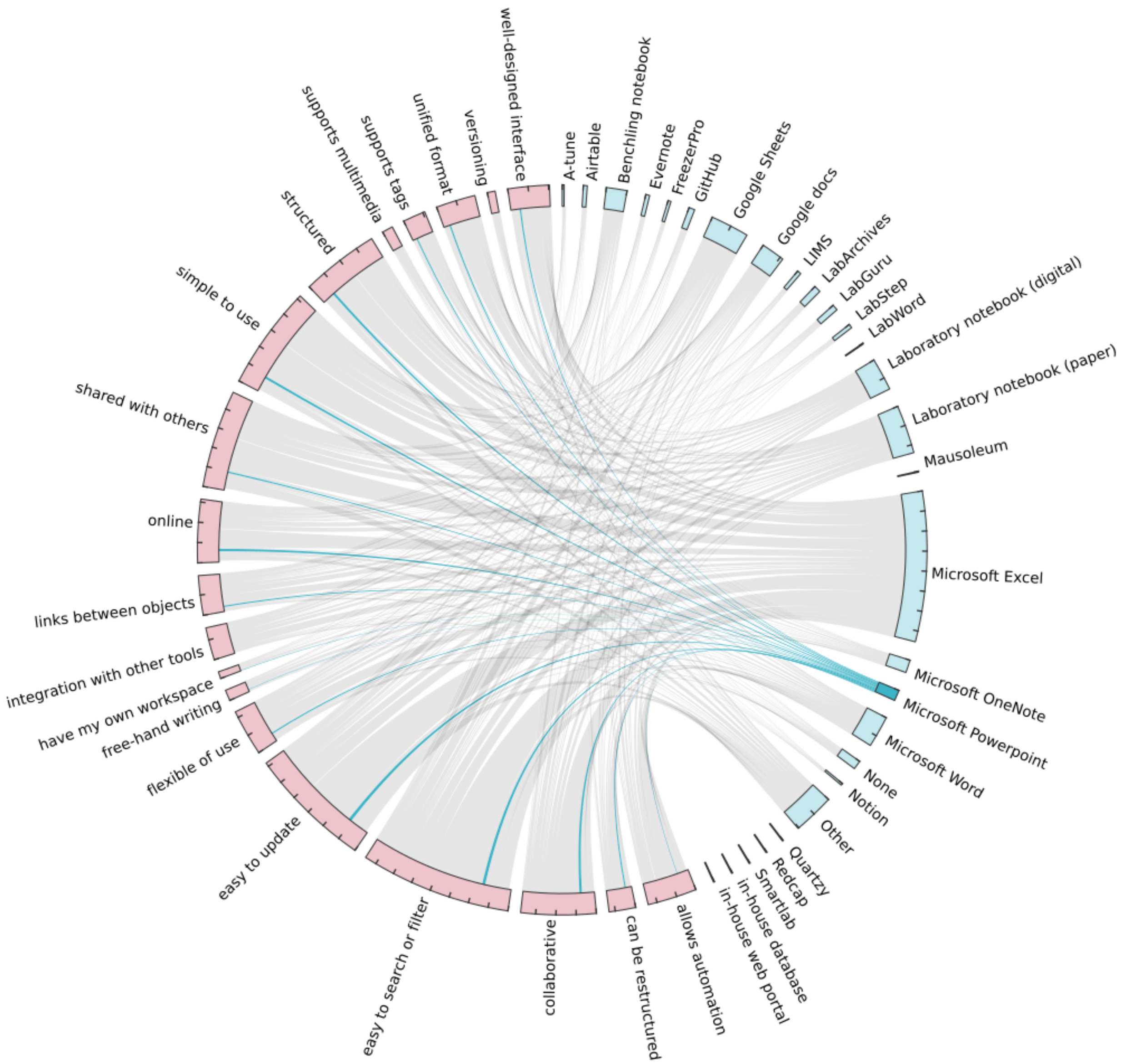


Sample-management solutions (blue arcs) vs desired features (red arcs). Arc length depicts the number of respondents, with ticks denoting 20 respondents. Link thickness shows the fraction of respondents using a solution and valuing a criterion. The chords connected to the arc 'Microsoft Powerpoint' are highlighted in color.'

Microsoft Word

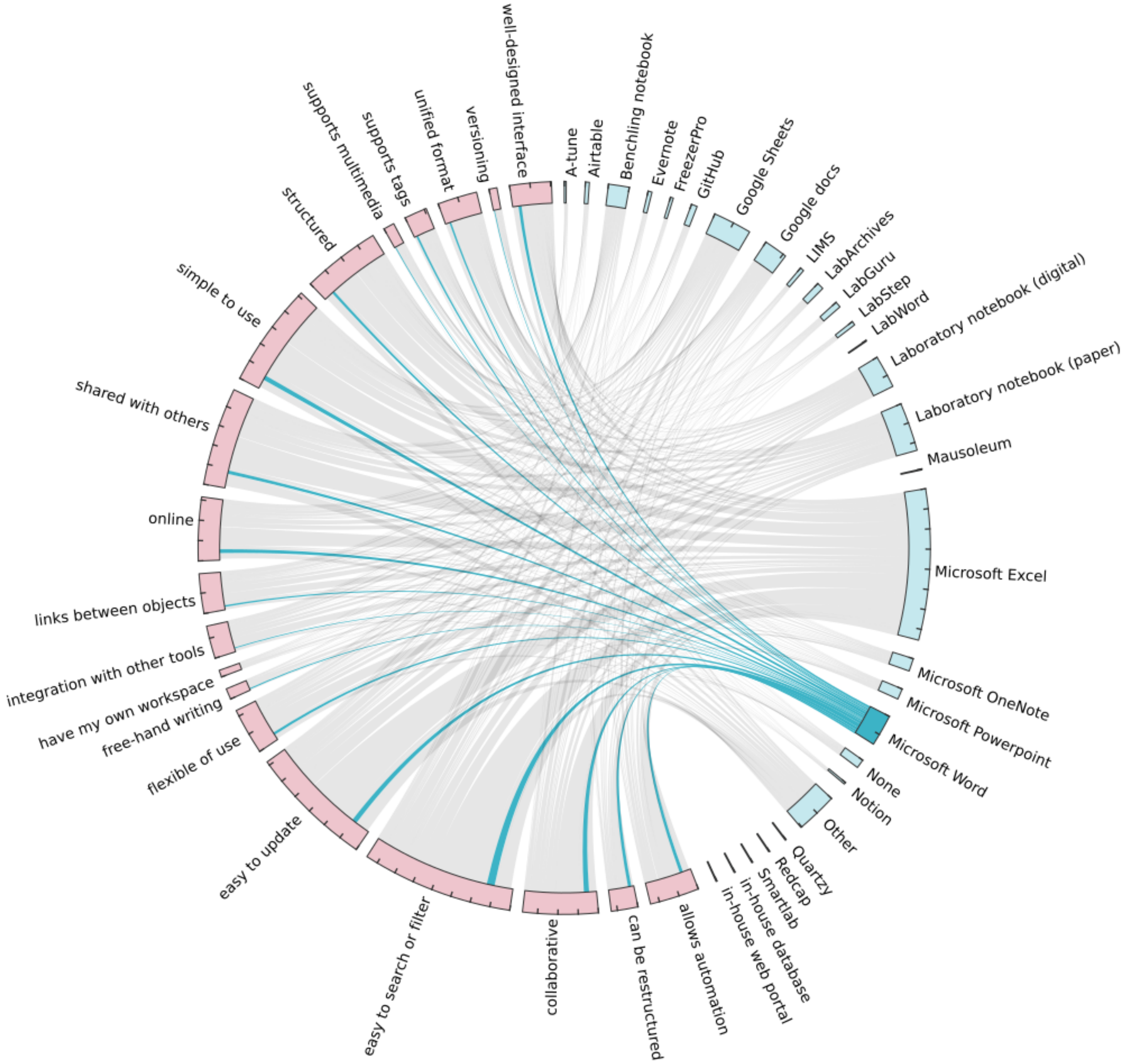


Sample-management solutions (blue arcs) vs desired features (red arcs). Arc length depicts the number of respondents, with ticks denoting 20 respondents. Link thickness shows the fraction of respondents using a solution and valuing a criterion. The chords connected to the arc 'Microsoft Word' are highlighted in color.'

None

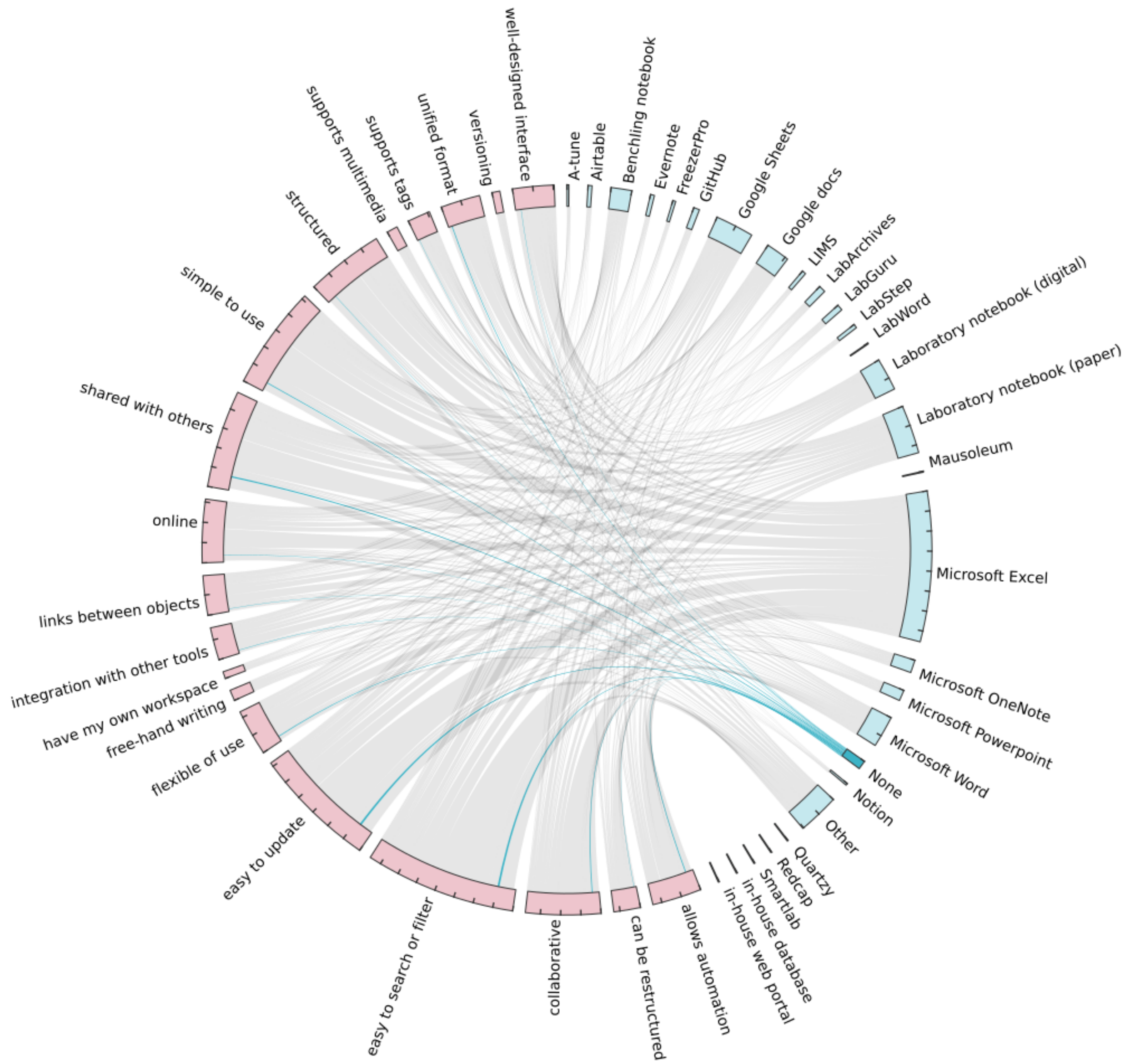


Sample-management solutions (blue arcs) vs desired features (red arcs). Arc length depicts the number of respondents, with ticks denoting 20 respondents. Link thickness shows the fraction of respondents using a solution and valuing a criterion. The chords connected to the arc 'None' are highlighted in color.'

Notion

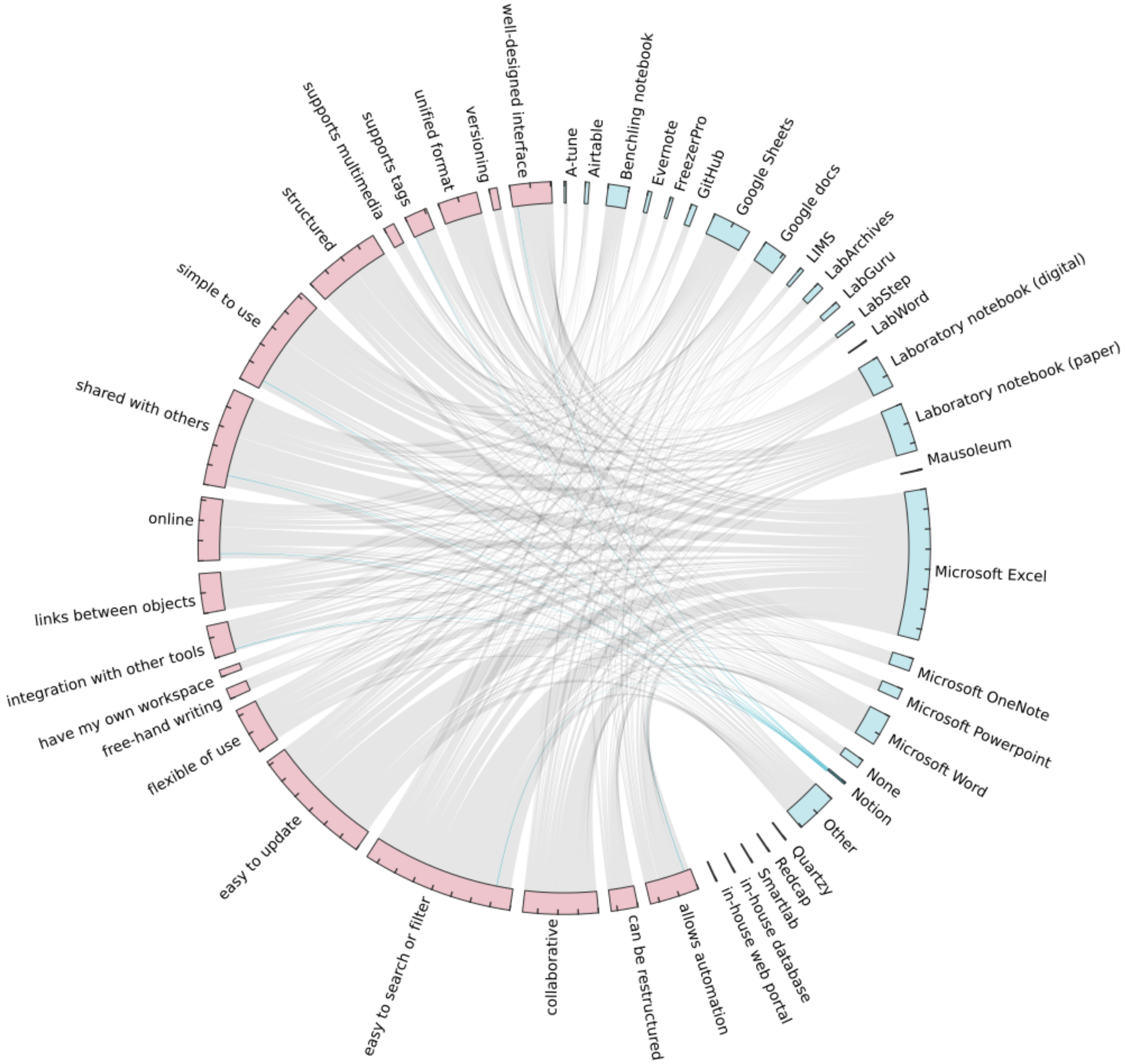


Sample-management solutions (blue arcs) vs desired features (red arcs). Arc length depicts the number of respondents, with ticks denoting 20 respondents. Link thickness shows the fraction of respondents using a solution and valuing a criterion. The chords connected to the arc 'Notion' are highlighted in color.'

Other

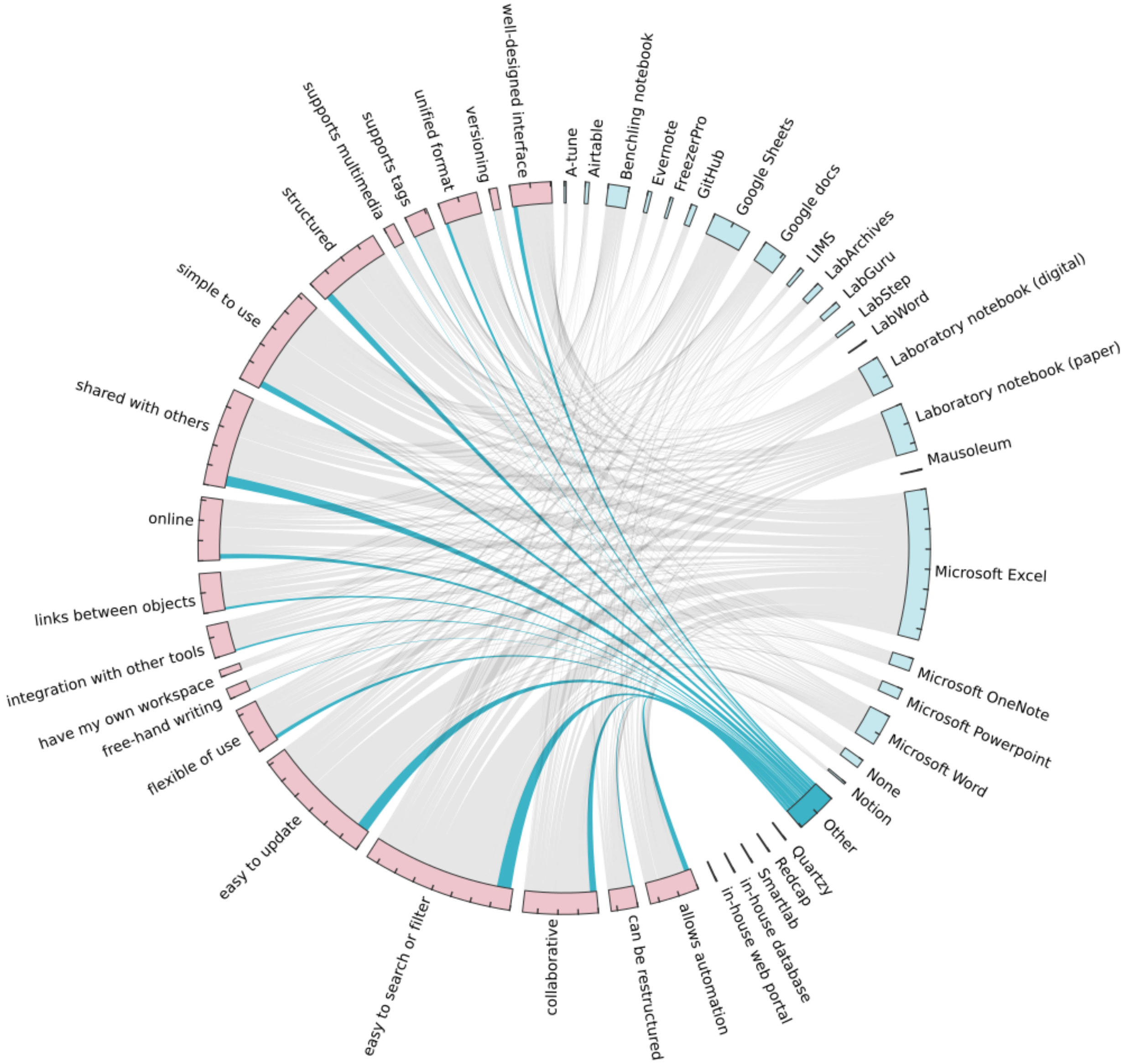


Sample-management solutions (blue arcs) vs desired features (red arcs). Arc length depicts the number of respondents, with ticks denoting 20 respondents. Link thickness shows the fraction of respondents using a solution and valuing a criterion. The chords connected to the arc 'Other' are highlighted in color.'

Quartzy

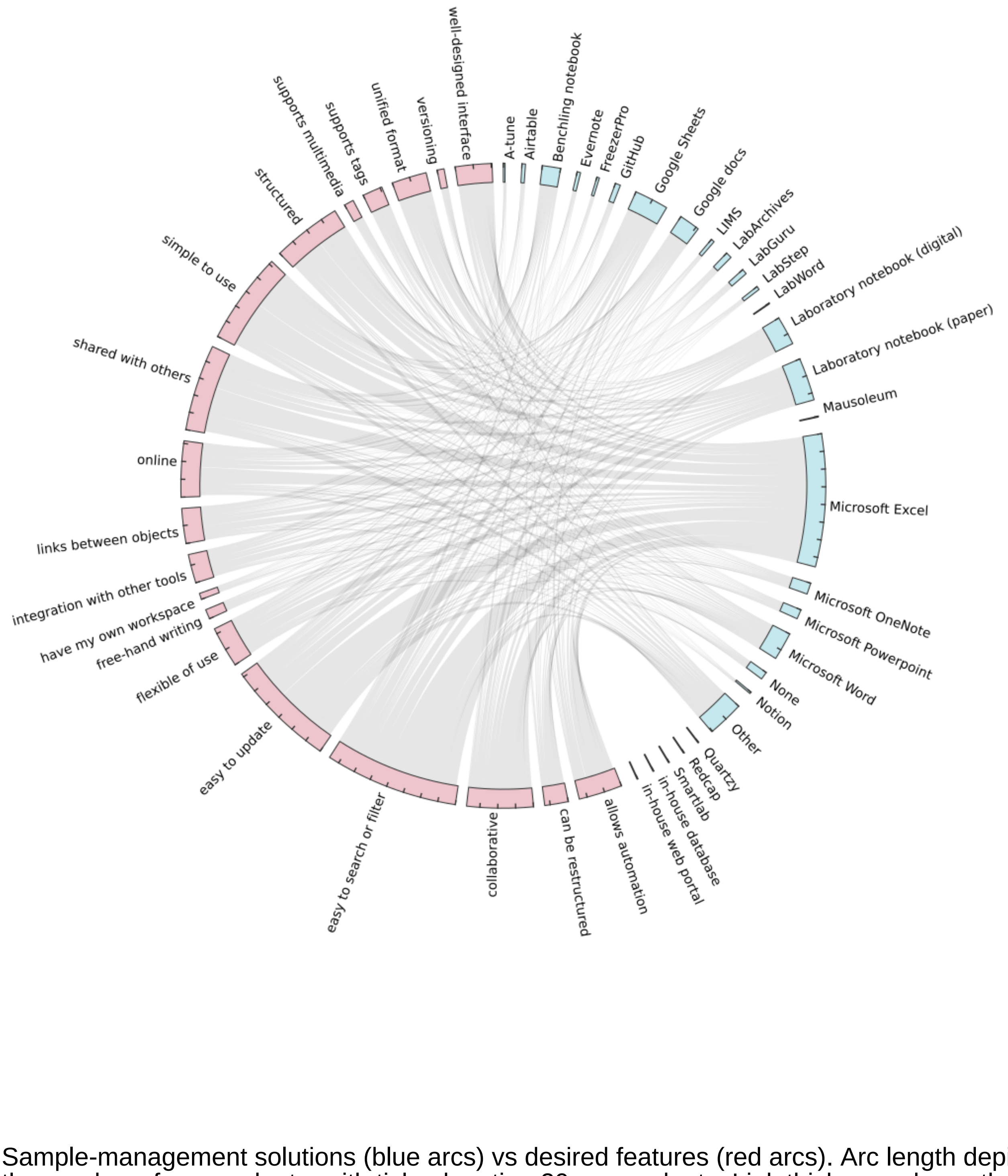


Sample-management solutions (blue arcs) vs desired features (red arcs). Arc length depicts the number of respondents, with ticks denoting 20 respondents. Link thickness shows the fraction of respondents using a solution and valuing a criterion. The chords connected to the arc 'Quartzy' are highlighted in color.'

Redcap

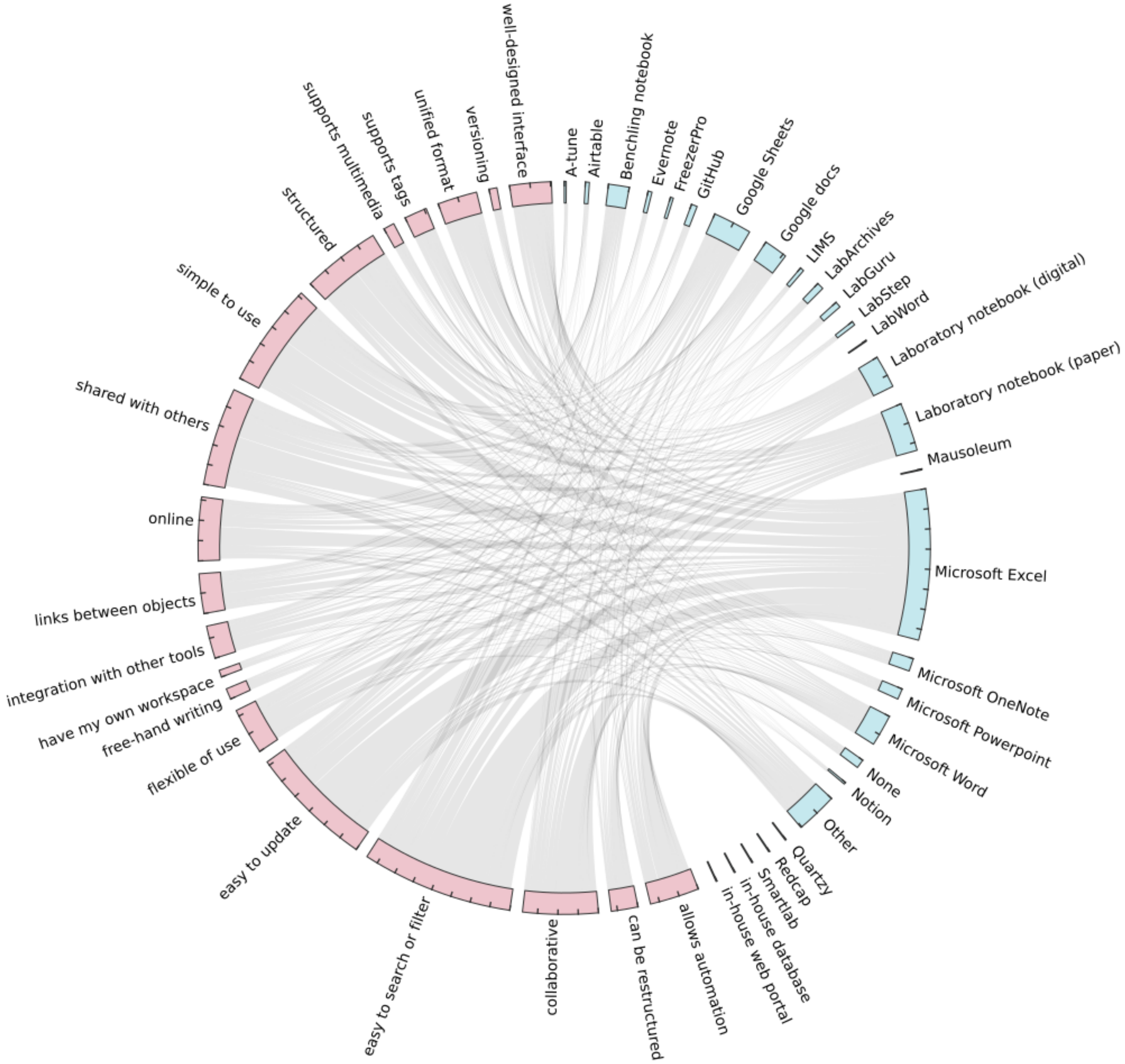


Sample-management solutions (blue arcs) vs desired features (red arcs). Arc length depicts the number of respondents, with ticks denoting 20 respondents. Link thickness shows the fraction of respondents using a solution and valuing a criterion. The chords connected to the arc 'Redcap' are highlighted in color.'

Smartlab

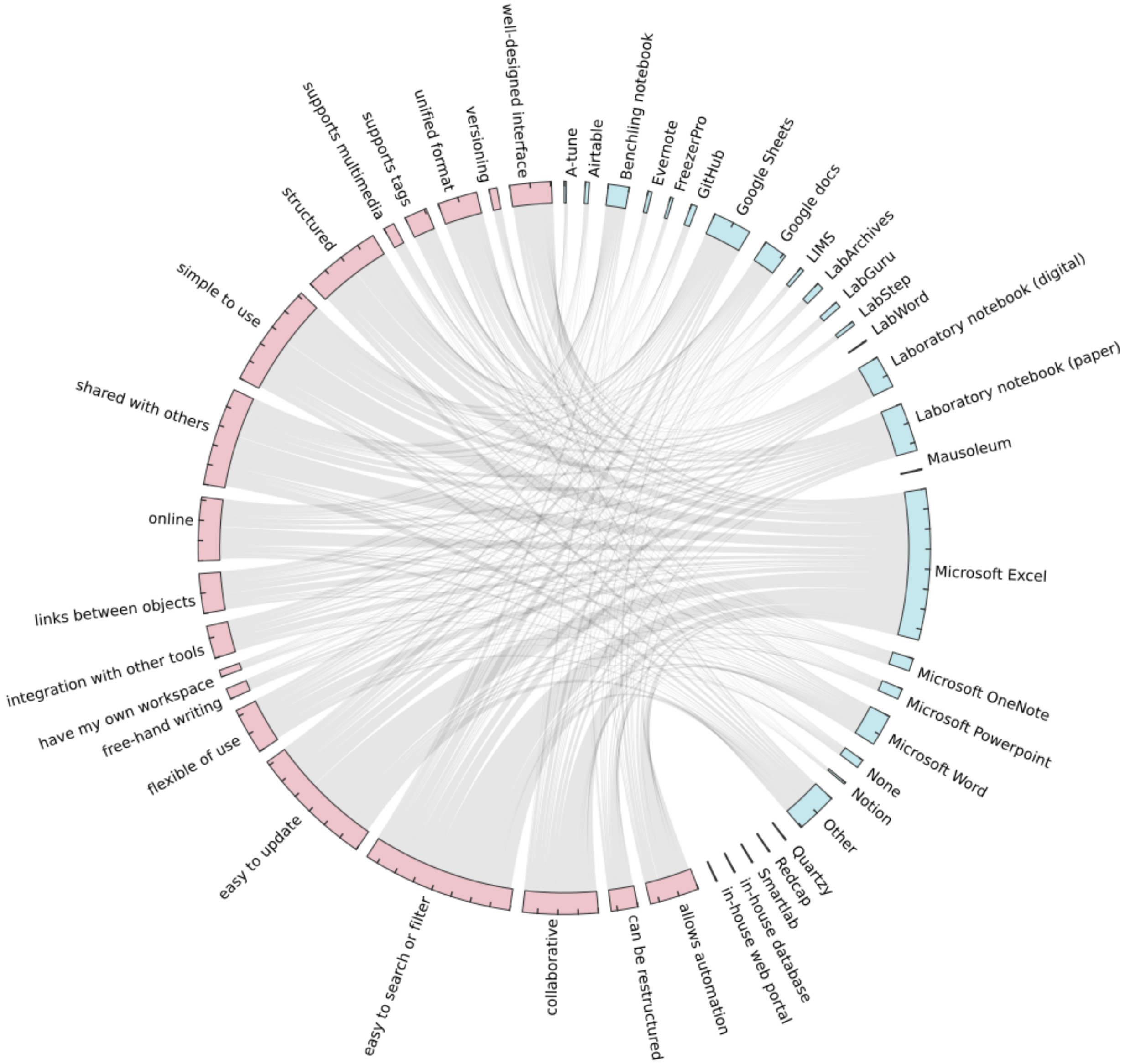


Sample-management solutions (blue arcs) vs desired features (red arcs). Arc length depicts the number of respondents, with ticks denoting 20 respondents. Link thickness shows the fraction of respondents using a solution and valuing a criterion. The chords connected to the arc 'Smartlab' are highlighted in color.'

in-house database

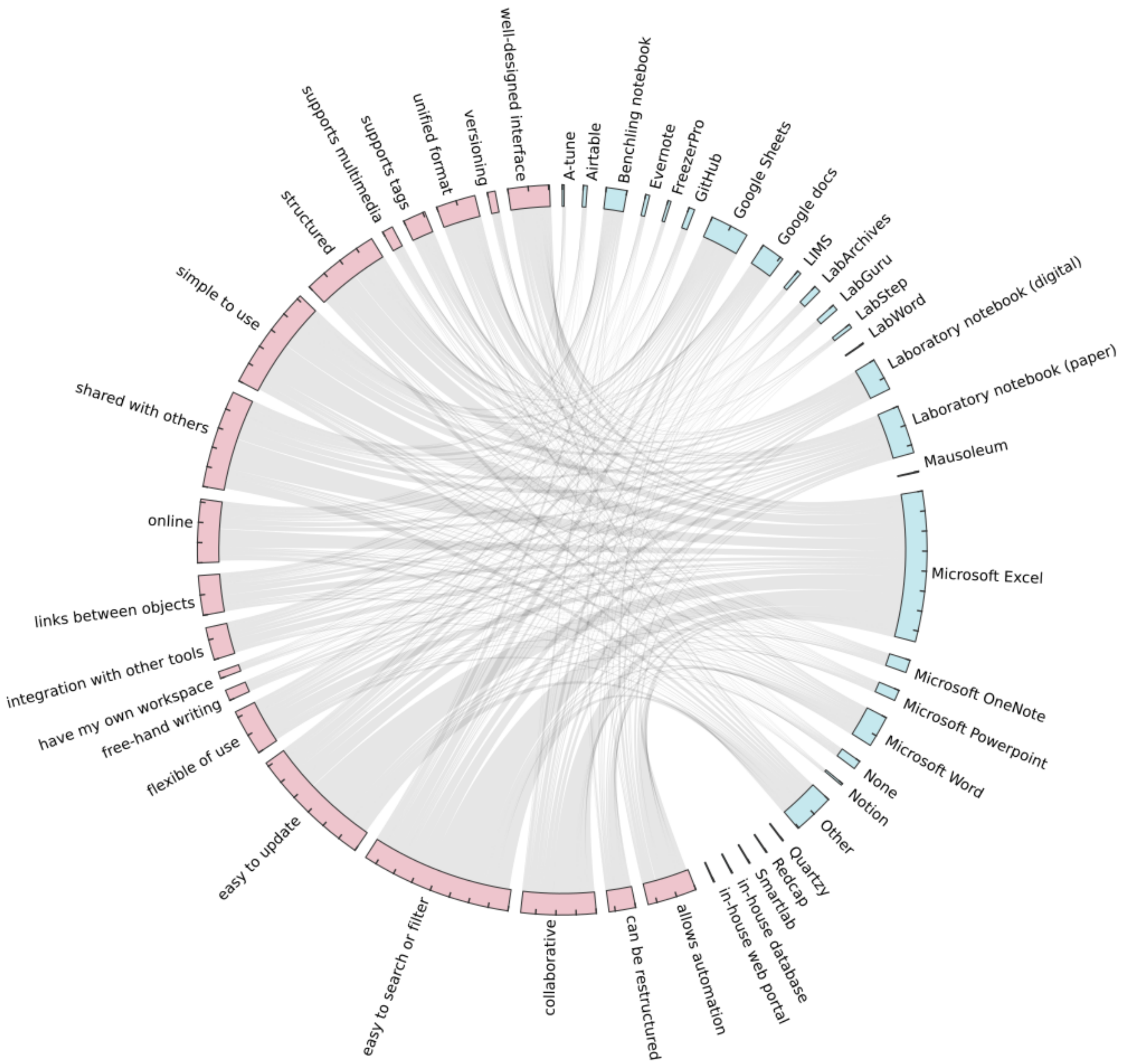


Sample-management solutions (blue arcs) vs desired features (red arcs). Arc length depicts the number of respondents, with ticks denoting 20 respondents. Link thickness shows the fraction of respondents using a solution and valuing a criterion. The chords connected to the arc 'in-house database' are highlighted in color.'

in-house web portal

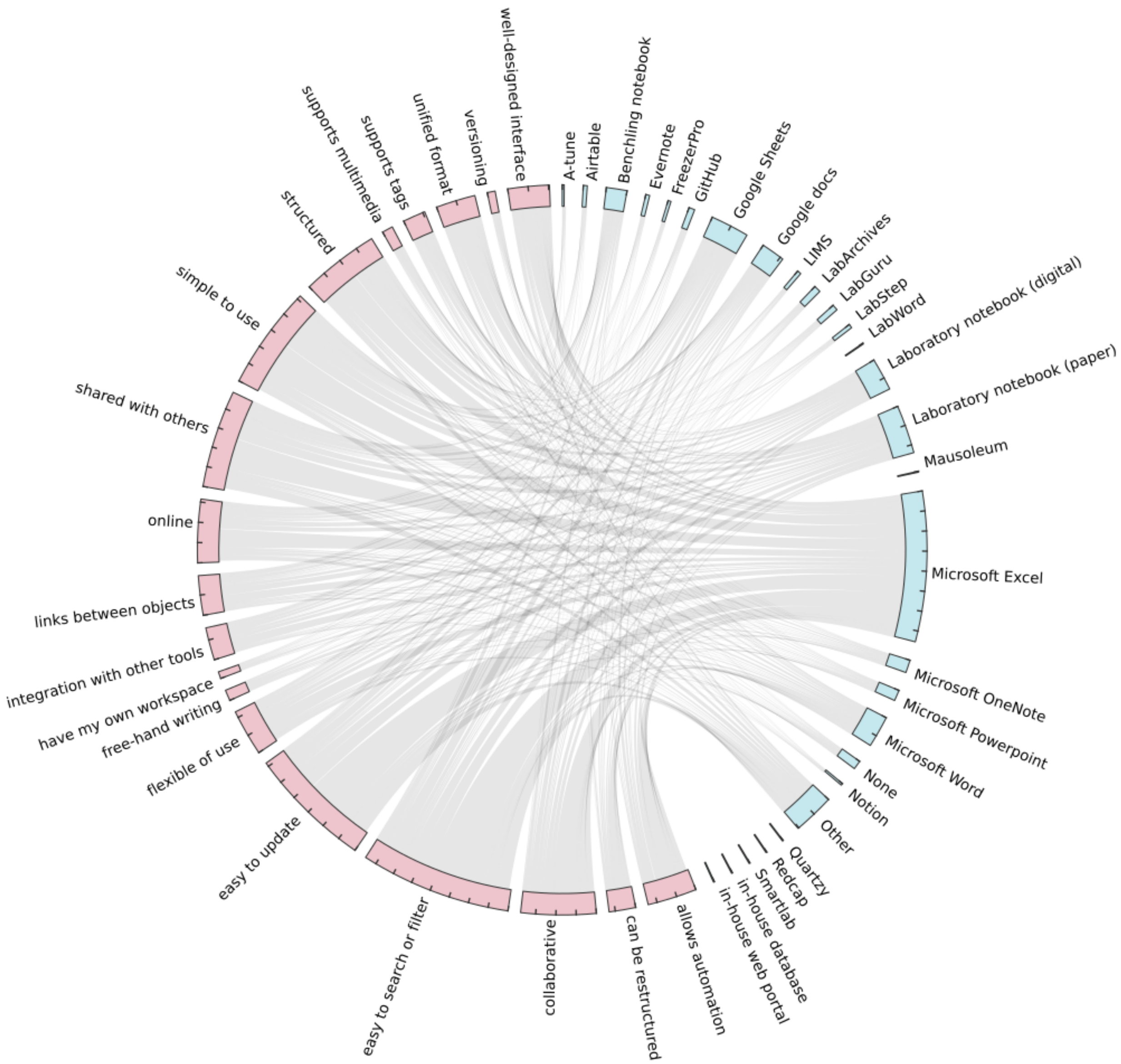


Sample-management solutions (blue arcs) vs desired features (red arcs). Arc length depicts the number of respondents, with ticks denoting 20 respondents. Link thickness shows the fraction of respondents using a solution and valuing a criterion. The chords connected to the arc 'in-house web portal' are highlighted in color.'

allows automation

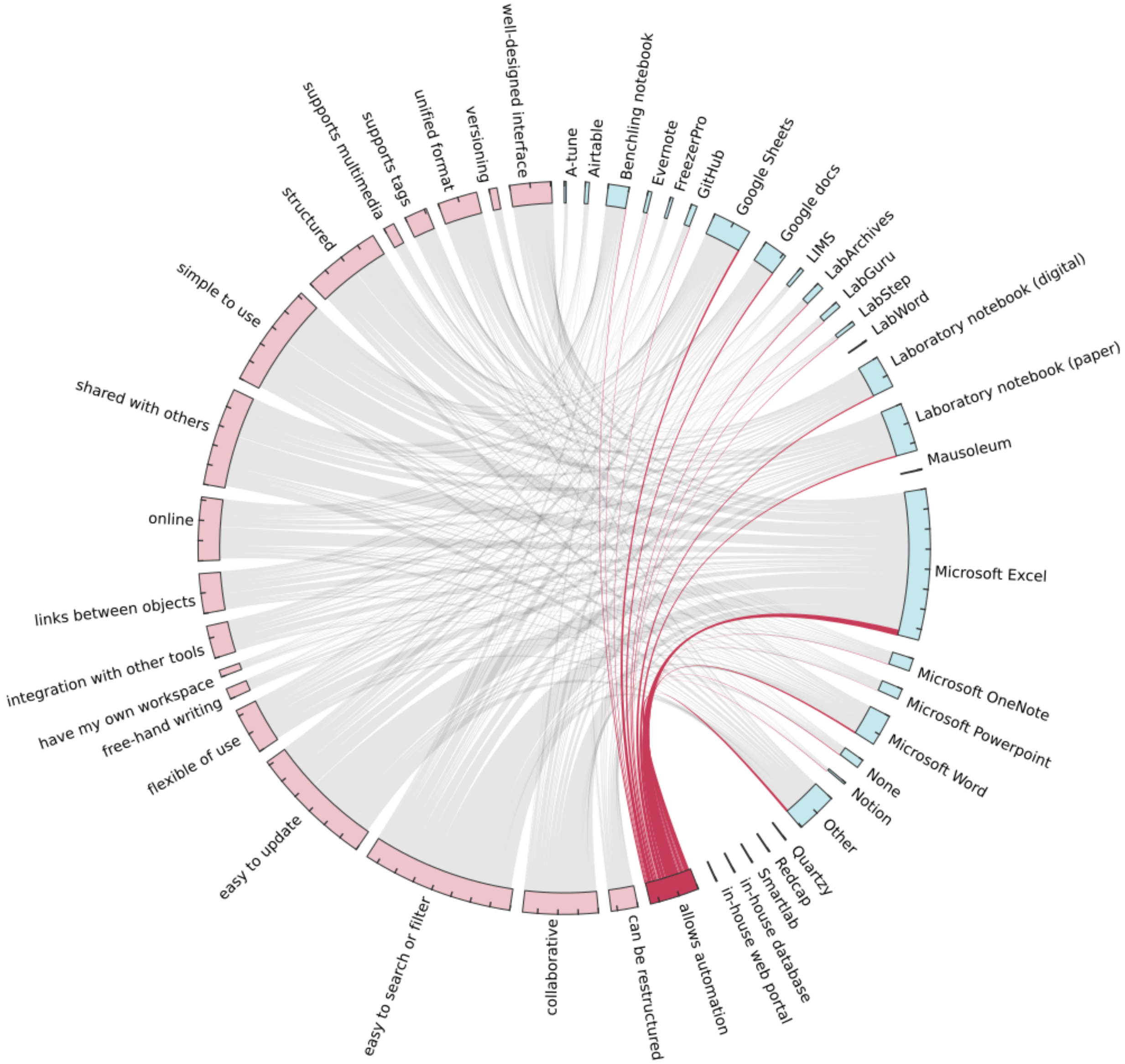


Sample-management solutions (blue arcs) vs desired features (red arcs). Arc length depicts the number of respondents, with ticks denoting 20 respondents. Link thickness shows the fraction of respondents using a solution and valuing a criterion. The chords connected to the arc 'allows automation' are highlighted in color.'

can be restructured

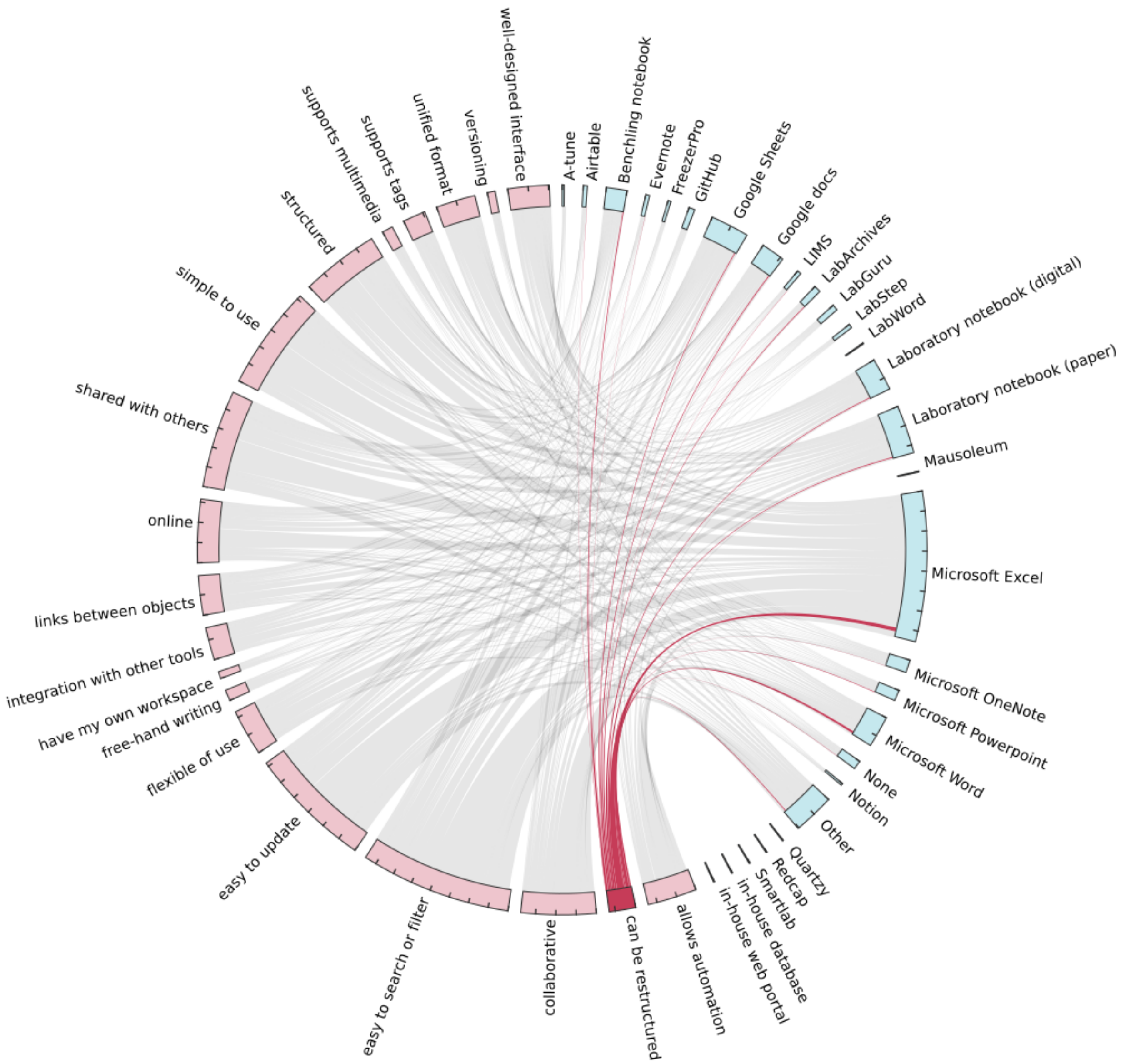


Sample-management solutions (blue arcs) vs desired features (red arcs). Arc length depicts the number of respondents, with ticks denoting 20 respondents. Link thickness shows the fraction of respondents using a solution and valuing a criterion. The chords connected to the arc 'can be restructured' are highlighted in color.'

collaborative

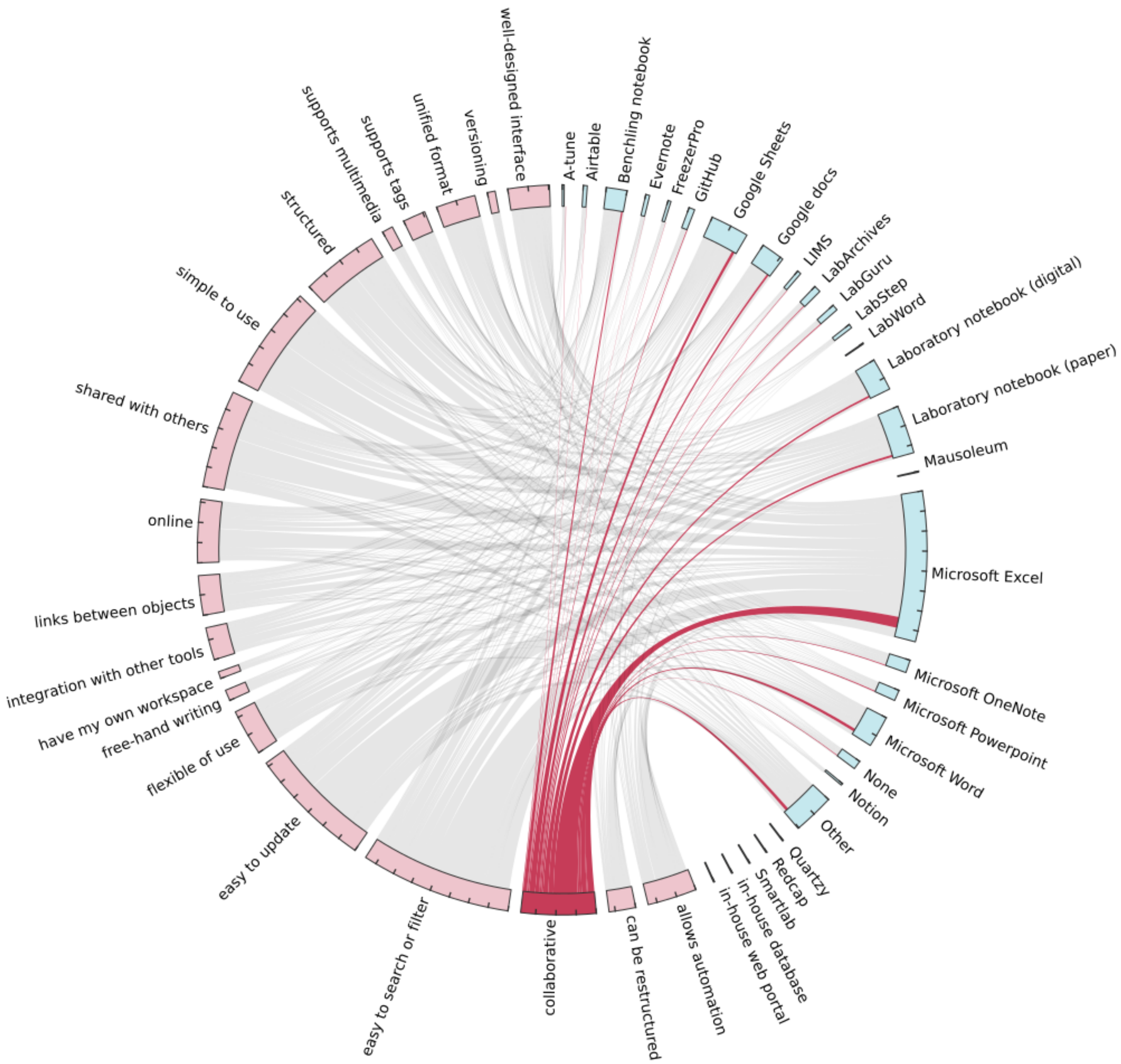


Sample-management solutions (blue arcs) vs desired features (red arcs). Arc length depicts the number of respondents, with ticks denoting 20 respondents. Link thickness shows the fraction of respondents using a solution and valuing a criterion. The chords connected to the arc 'collaborative' are highlighted in color.'

easy to search or filter

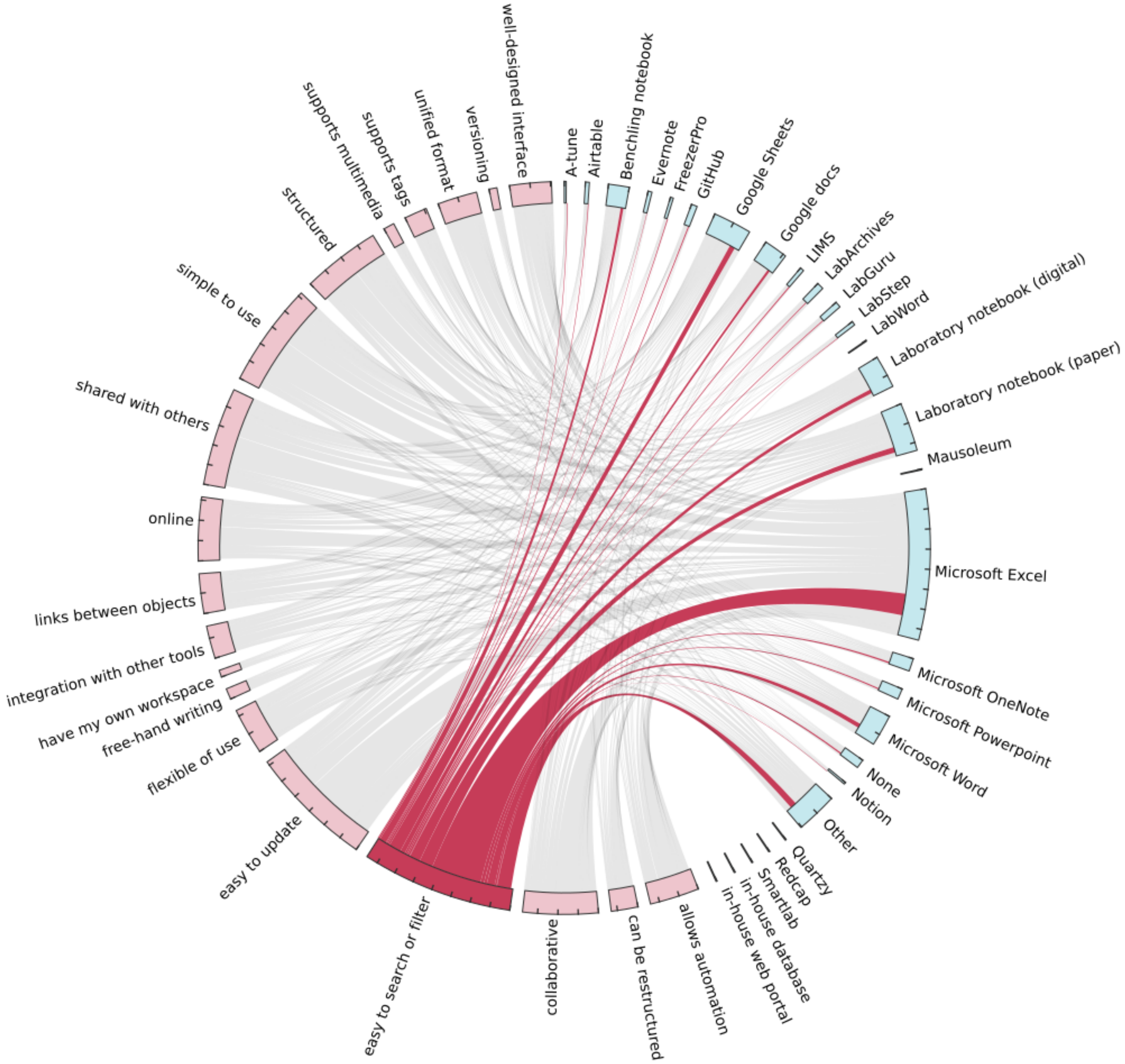


Sample-management solutions (blue arcs) vs desired features (red arcs). Arc length depicts the number of respondents, with ticks denoting 20 respondents. Link thickness shows the fraction of respondents using a solution and valuing a criterion. The chords connected to the arc 'easy to search or filter' are highlighted in color.'

easy to update

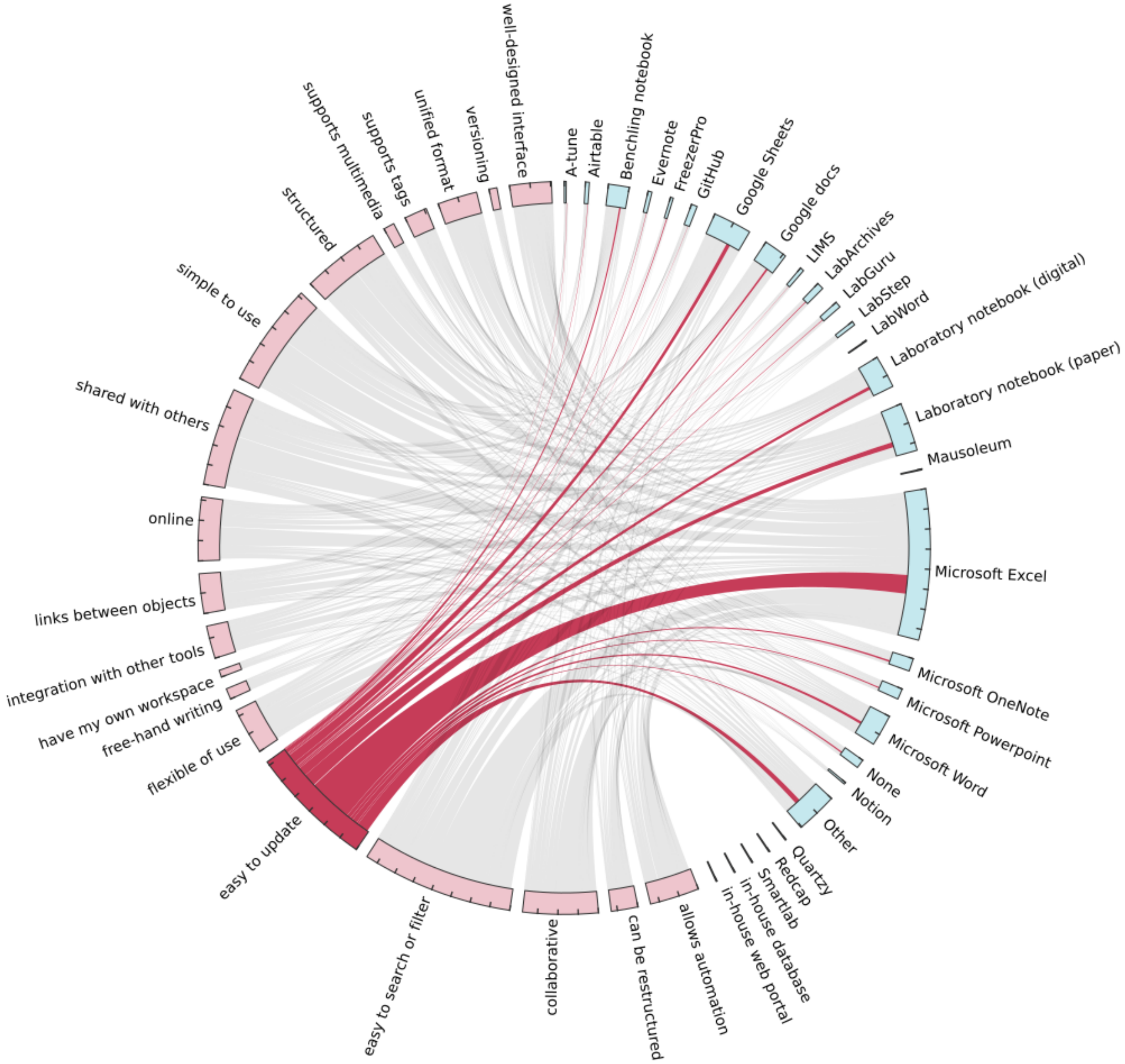


Sample-management solutions (blue arcs) vs desired features (red arcs). Arc length depicts the number of respondents, with ticks denoting 20 respondents. Link thickness shows the fraction of respondents using a solution and valuing a criterion. The chords connected to the arc 'easy to update' are highlighted in color.'

flexible of use

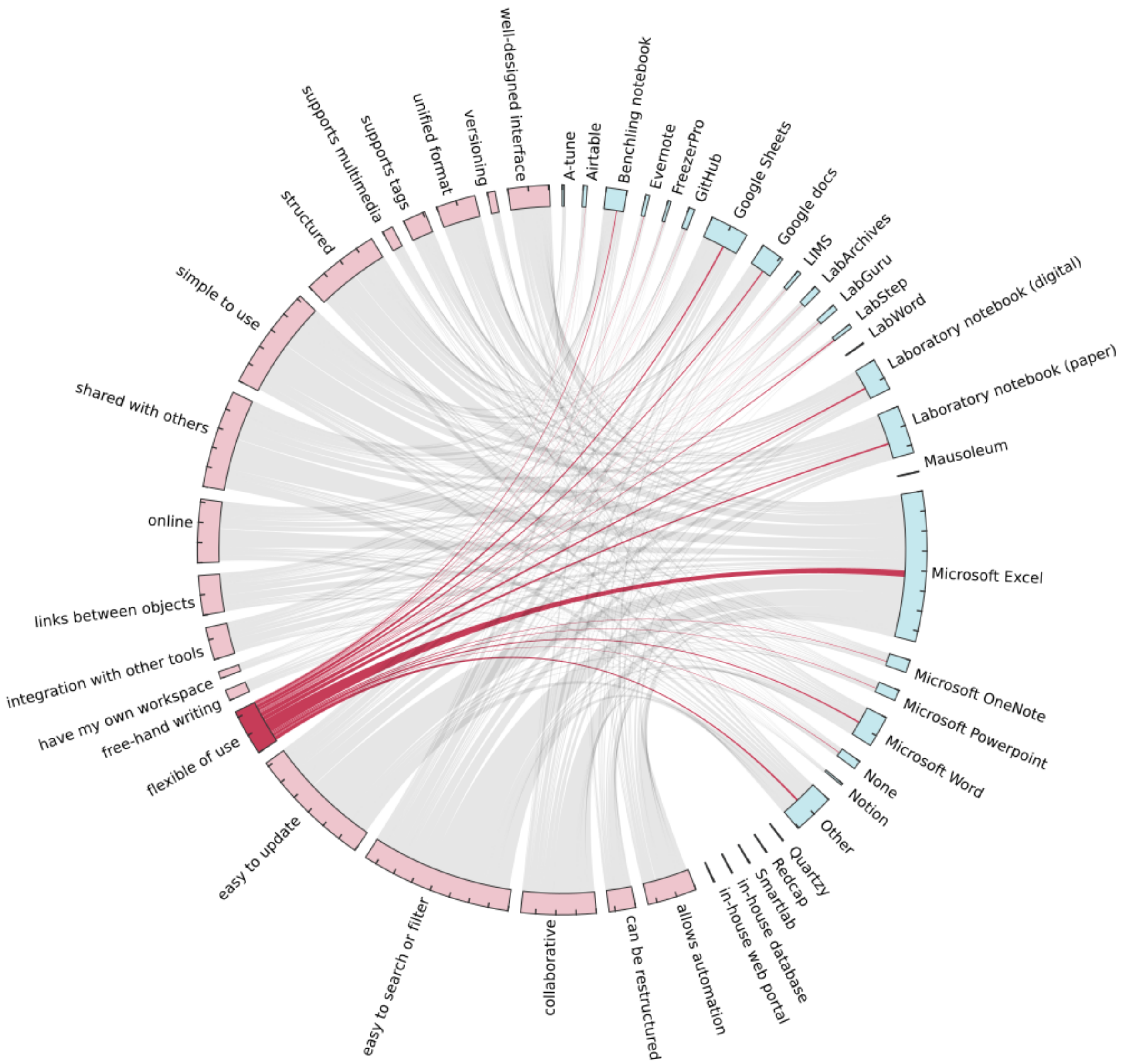


Sample-management solutions (blue arcs) vs desired features (red arcs). Arc length depicts the number of respondents, with ticks denoting 20 respondents. Link thickness shows the fraction of respondents using a solution and valuing a criterion. The chords connected to the arc 'flexible of use' are highlighted in color.'

free-hand writing

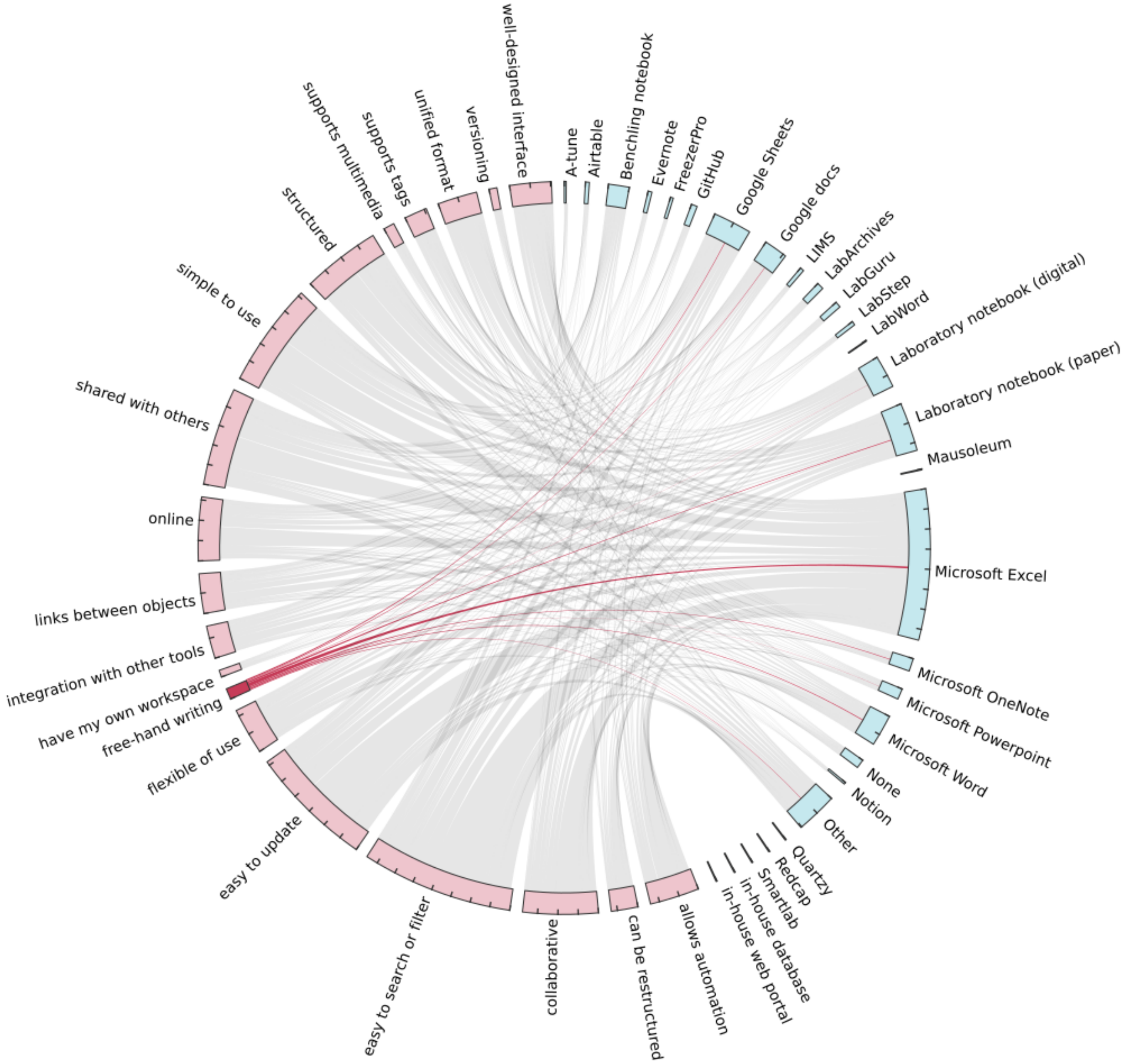


Sample-management solutions (blue arcs) vs desired features (red arcs). Arc length depicts the number of respondents, with ticks denoting 20 respondents. Link thickness shows the fraction of respondents using a solution and valuing a criterion. The chords connected to the arc 'free-hand writing' are highlighted in color.'

have my own workspace

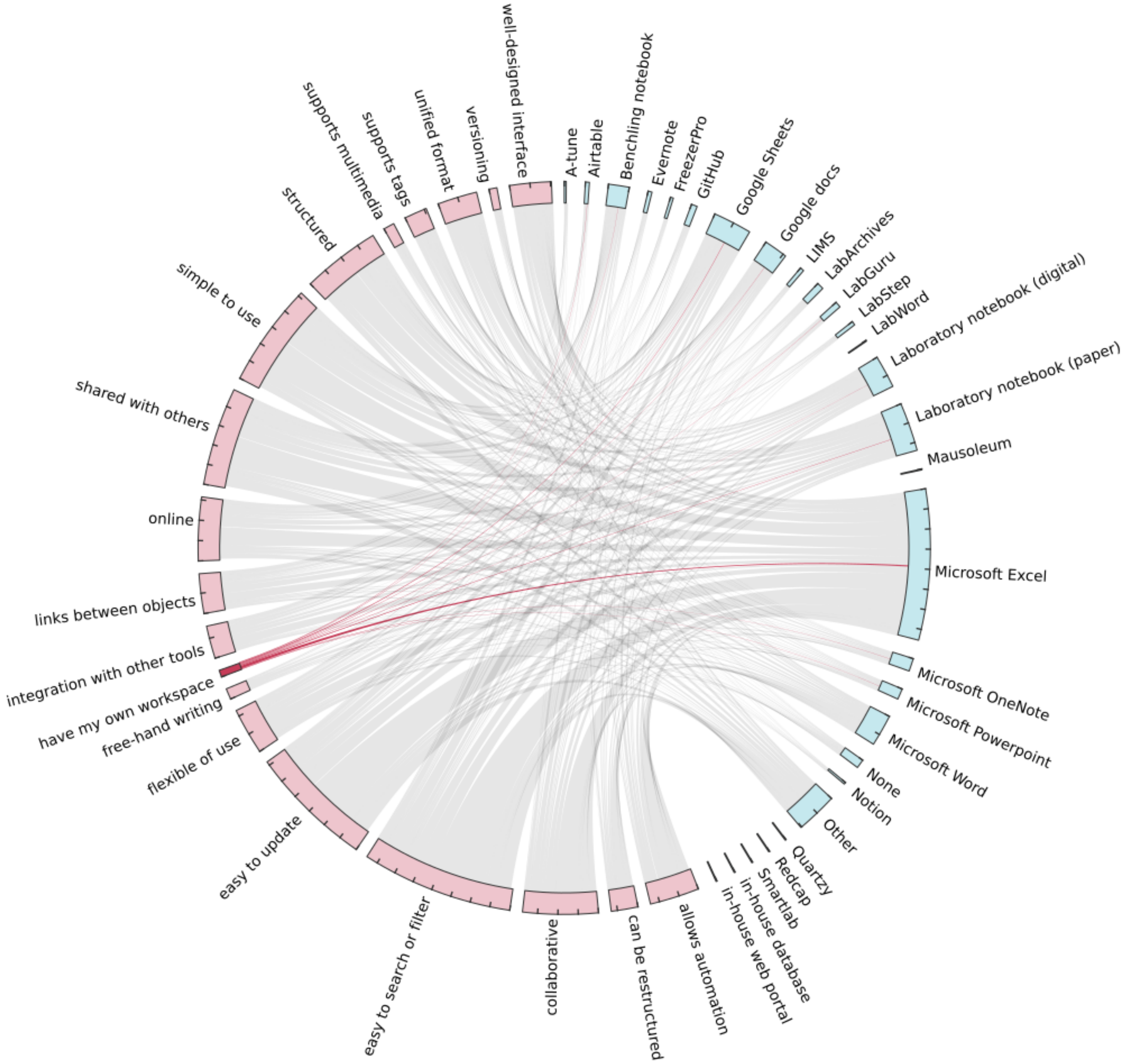


Sample-management solutions (blue arcs) vs desired features (red arcs). Arc length depicts the number of respondents, with ticks denoting 20 respondents. Link thickness shows the fraction of respondents using a solution and valuing a criterion. The chords connected to the arc 'have my own workspace' are highlighted in color.'

integration with other tools

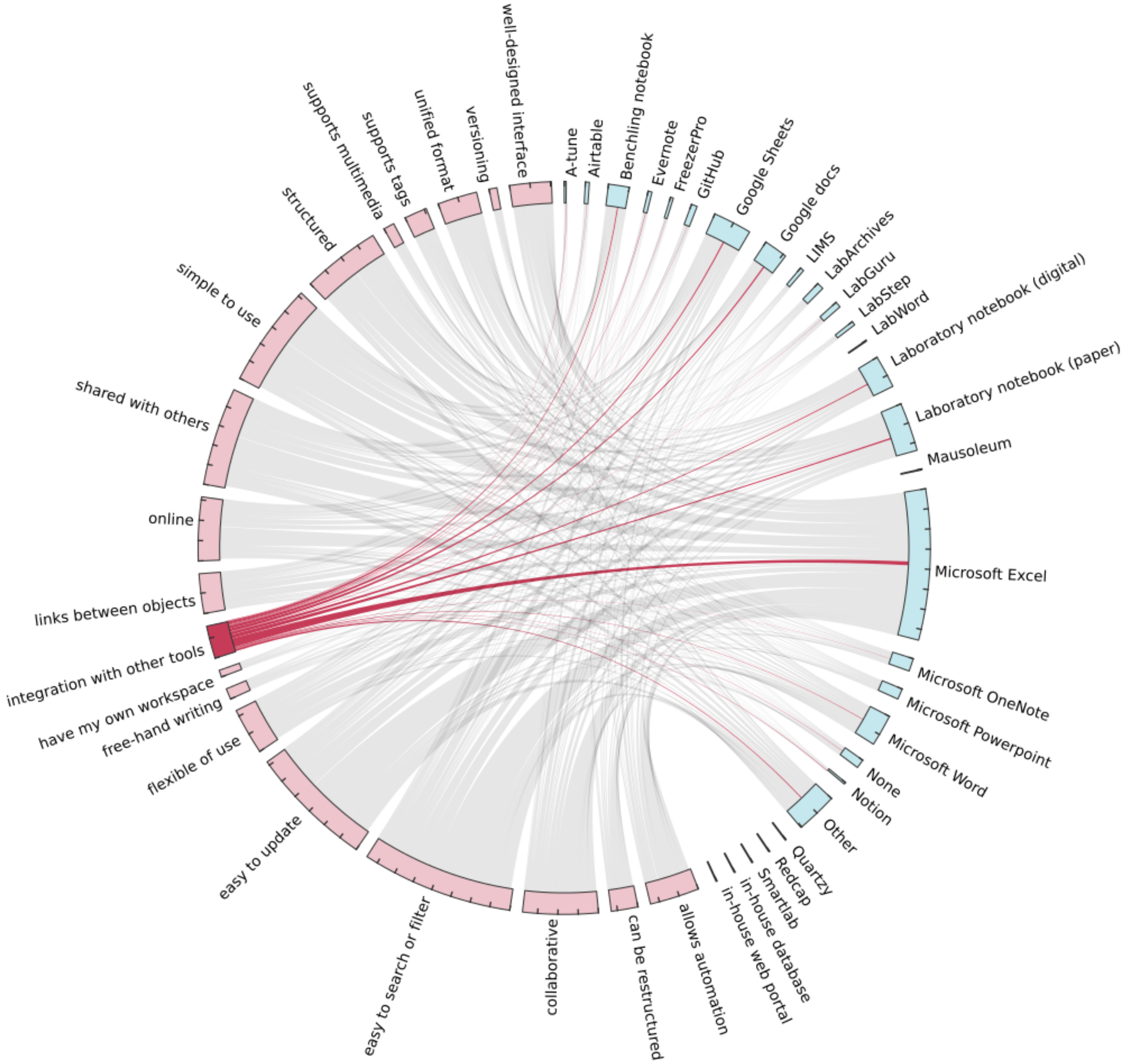


Sample-management solutions (blue arcs) vs desired features (red arcs). Arc length depicts the number of respondents, with ticks denoting 20 respondents. Link thickness shows the fraction of respondents using a solution and valuing a criterion. The chords connected to the arc 'integration with other tools' are highlighted in color.'

links between objects

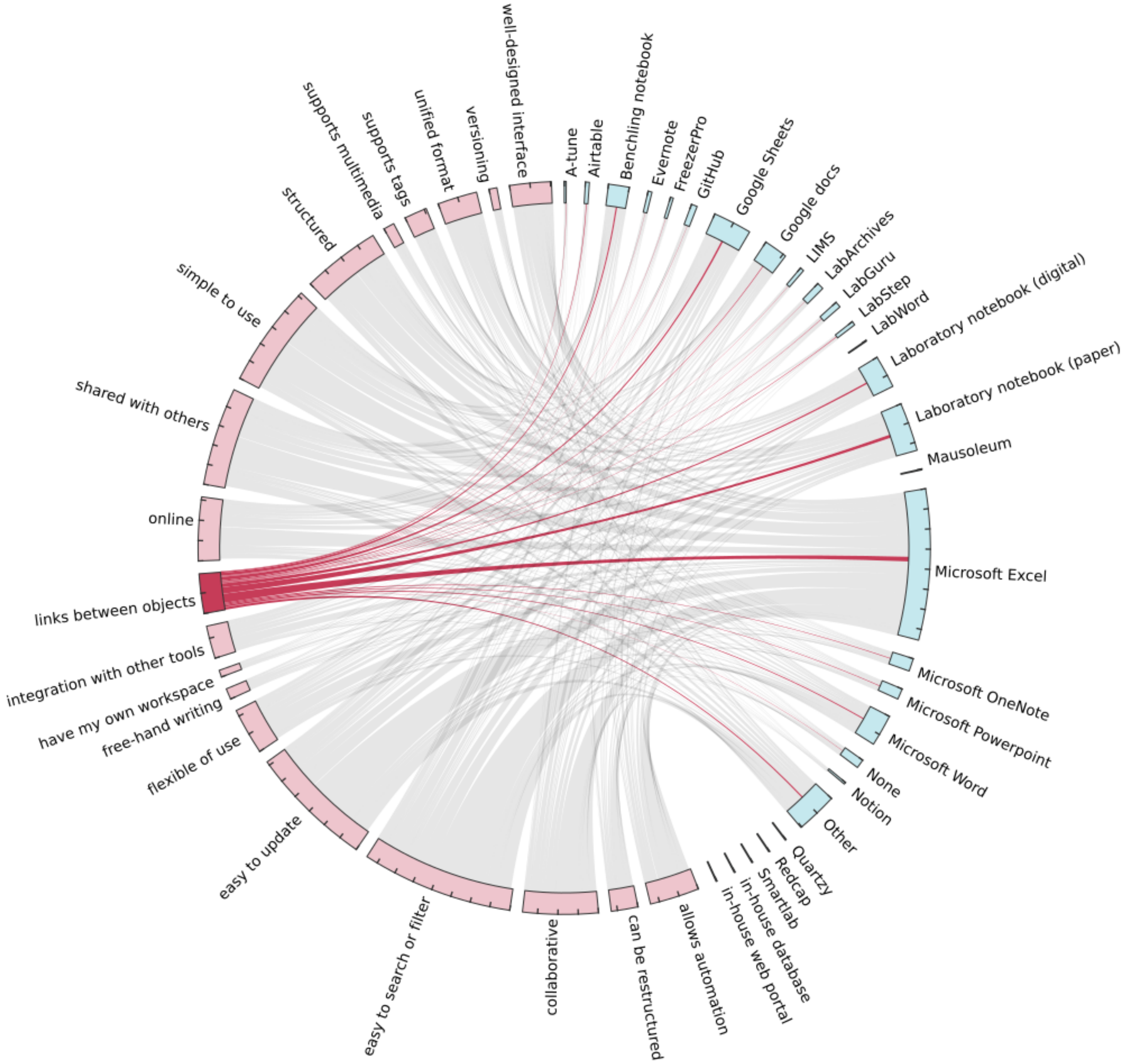


Sample-management solutions (blue arcs) vs desired features (red arcs). Arc length depicts the number of respondents, with ticks denoting 20 respondents. Link thickness shows the fraction of respondents using a solution and valuing a criterion. The chords connected to the arc 'links between objects' are highlighted in color.'

online

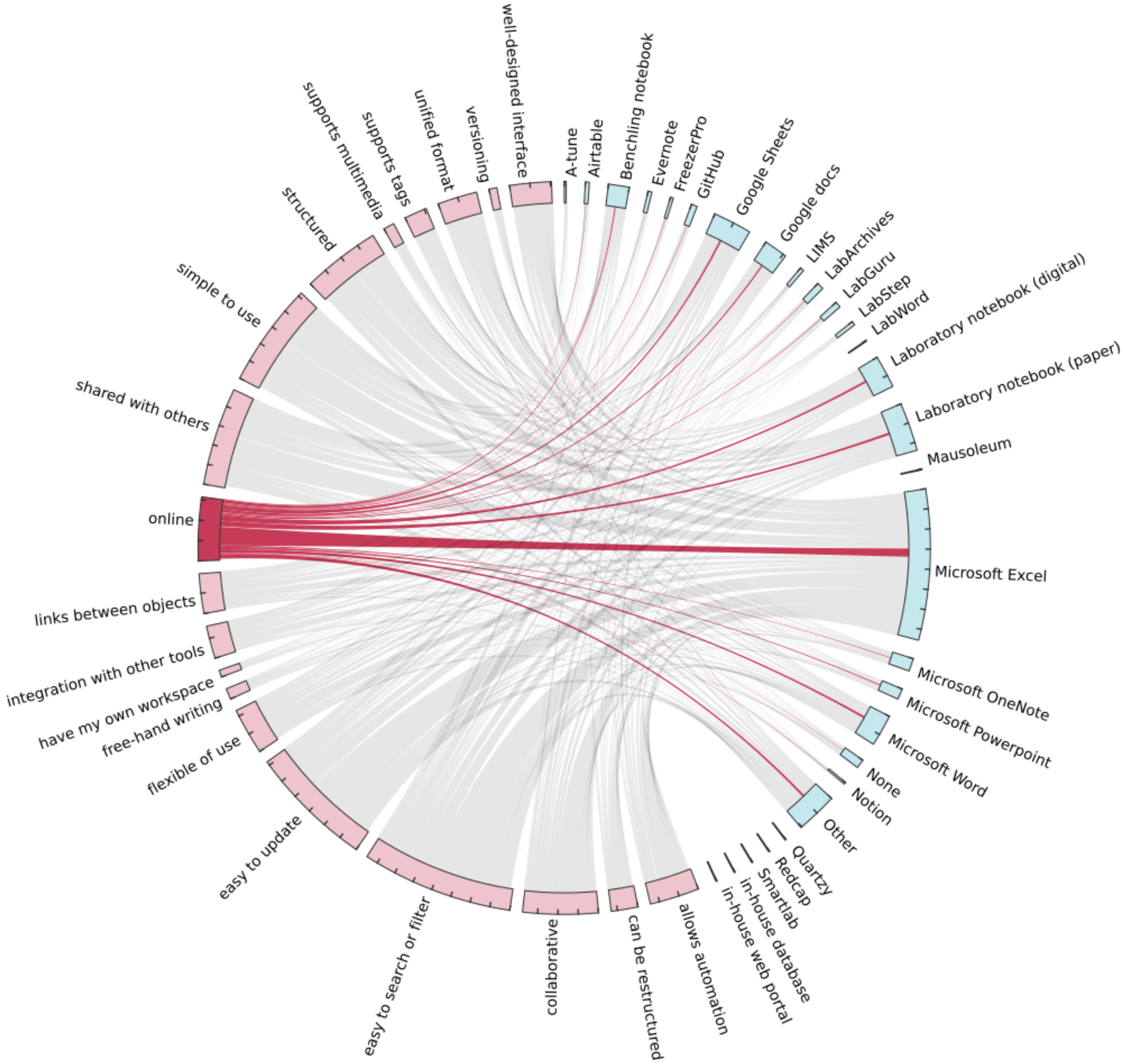


Sample-management solutions (blue arcs) vs desired features (red arcs). Arc length depicts the number of respondents, with ticks denoting 20 respondents. Link thickness shows the fraction of respondents using a solution and valuing a criterion. The chords connected to the arc 'online' are highlighted in color.'

shared with others

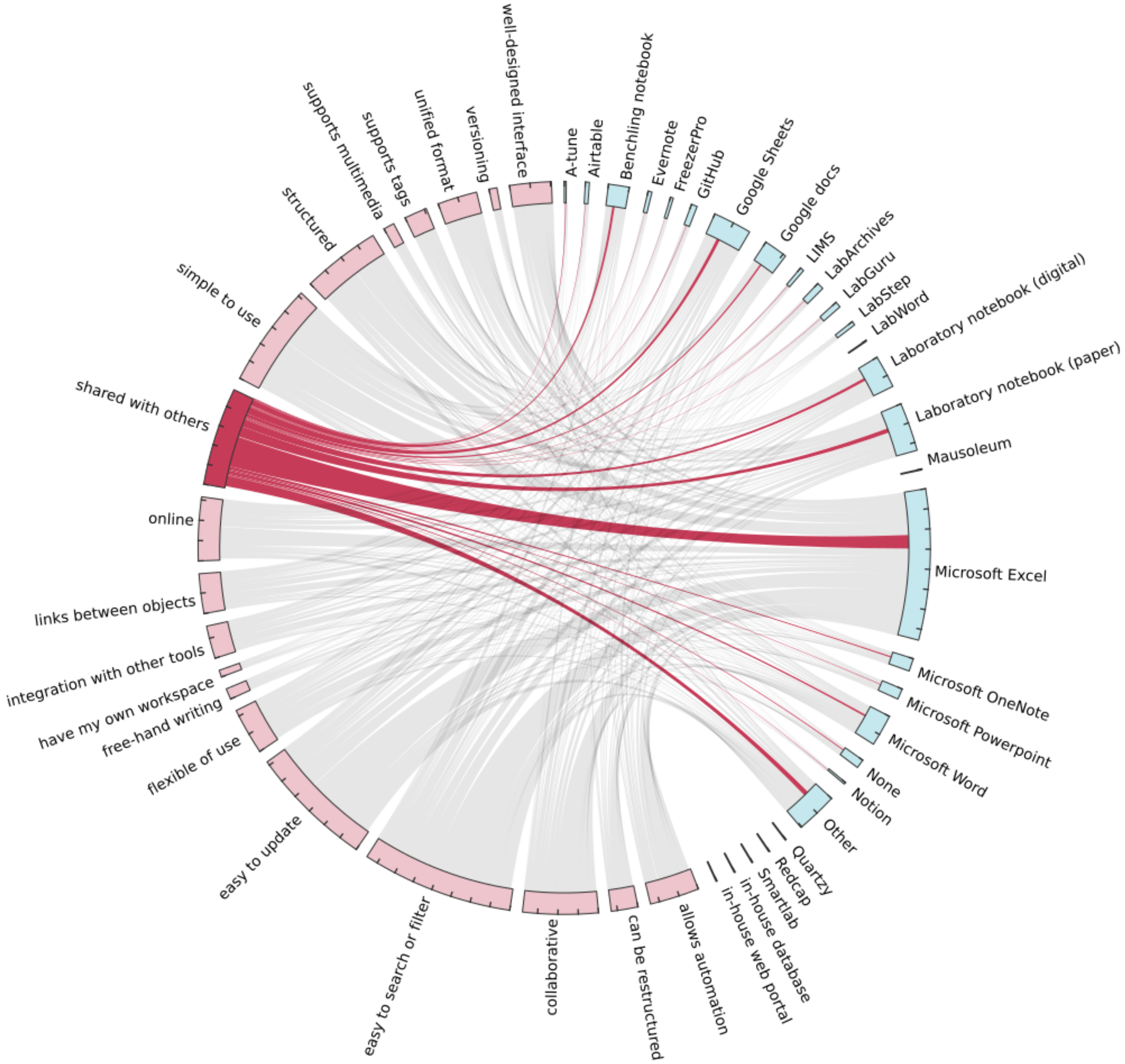


Sample-management solutions (blue arcs) vs desired features (red arcs). Arc length depicts the number of respondents, with ticks denoting 20 respondents. Link thickness shows the fraction of respondents using a solution and valuing a criterion. The chords connected to the arc 'shared with others' are highlighted in color.'

simple to use

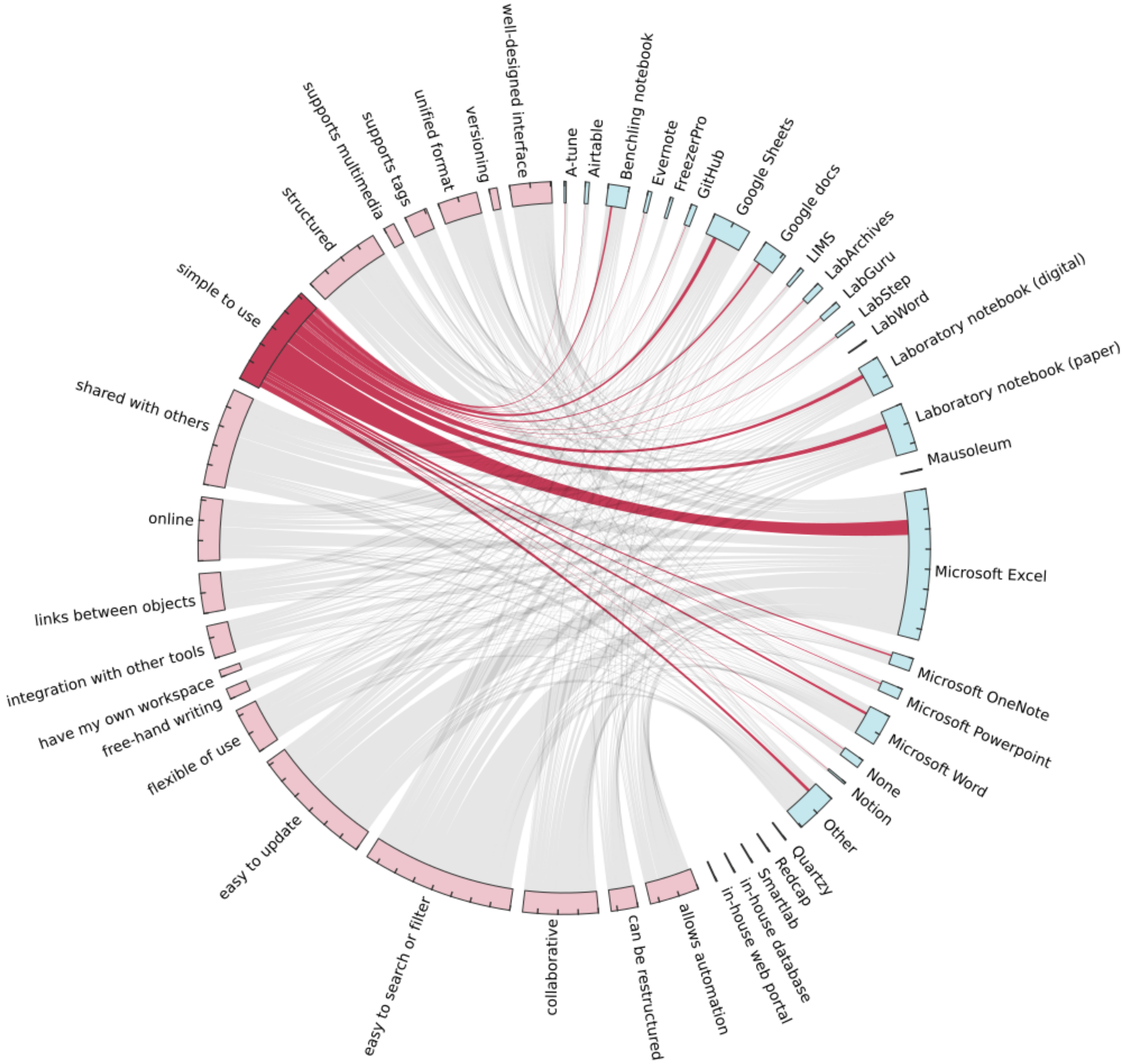


Sample-management solutions (blue arcs) vs desired features (red arcs). Arc length depicts the number of respondents, with ticks denoting 20 respondents. Link thickness shows the fraction of respondents using a solution and valuing a criterion. The chords connected to the arc 'simple to use' are highlighted in color.'

structured

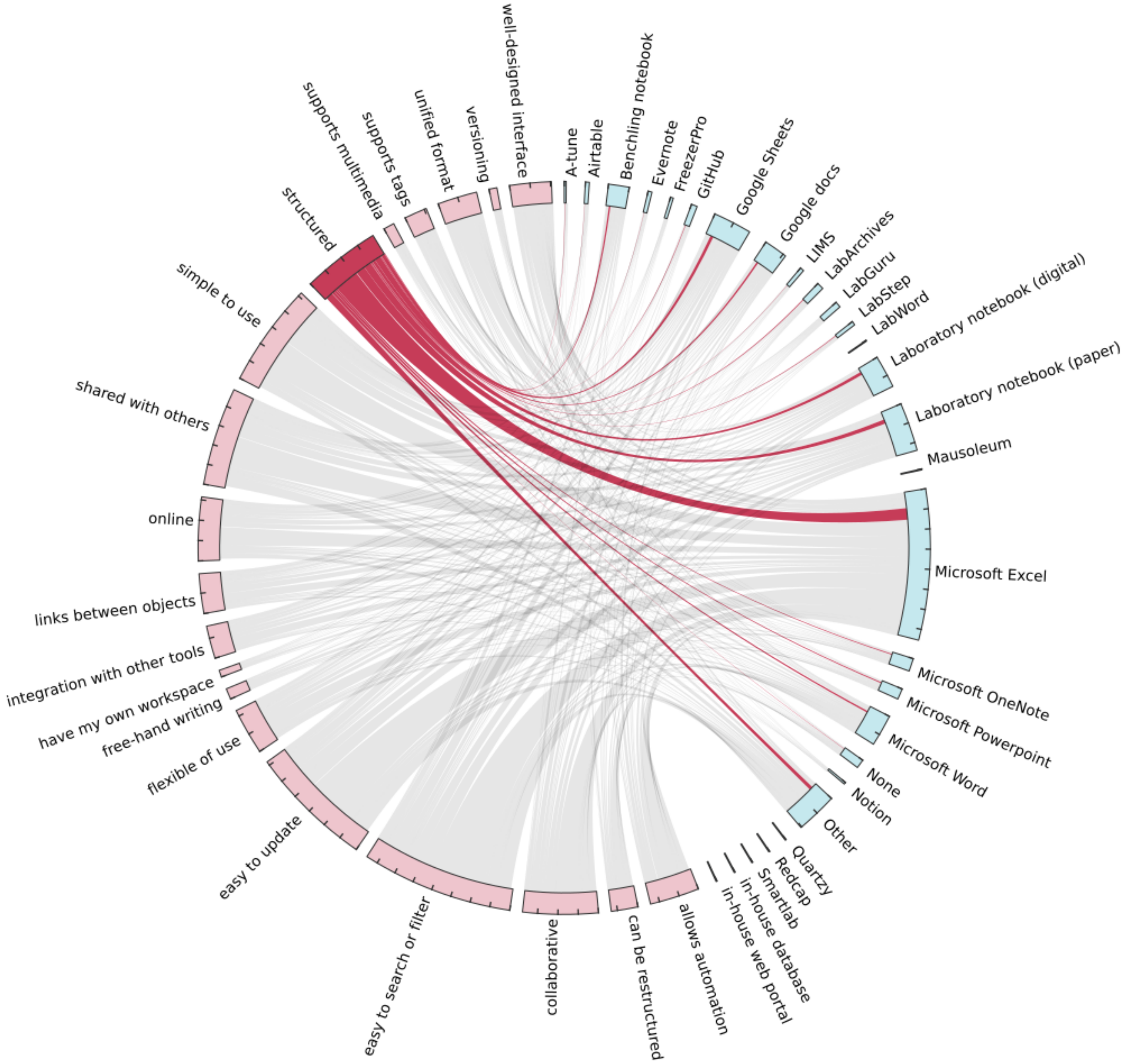


Sample-management solutions (blue arcs) vs desired features (red arcs). Arc length depicts the number of respondents, with ticks denoting 20 respondents. Link thickness shows the fraction of respondents using a solution and valuing a criterion. The chords connected to the arc 'structured' are highlighted in color.'

supports multimedia

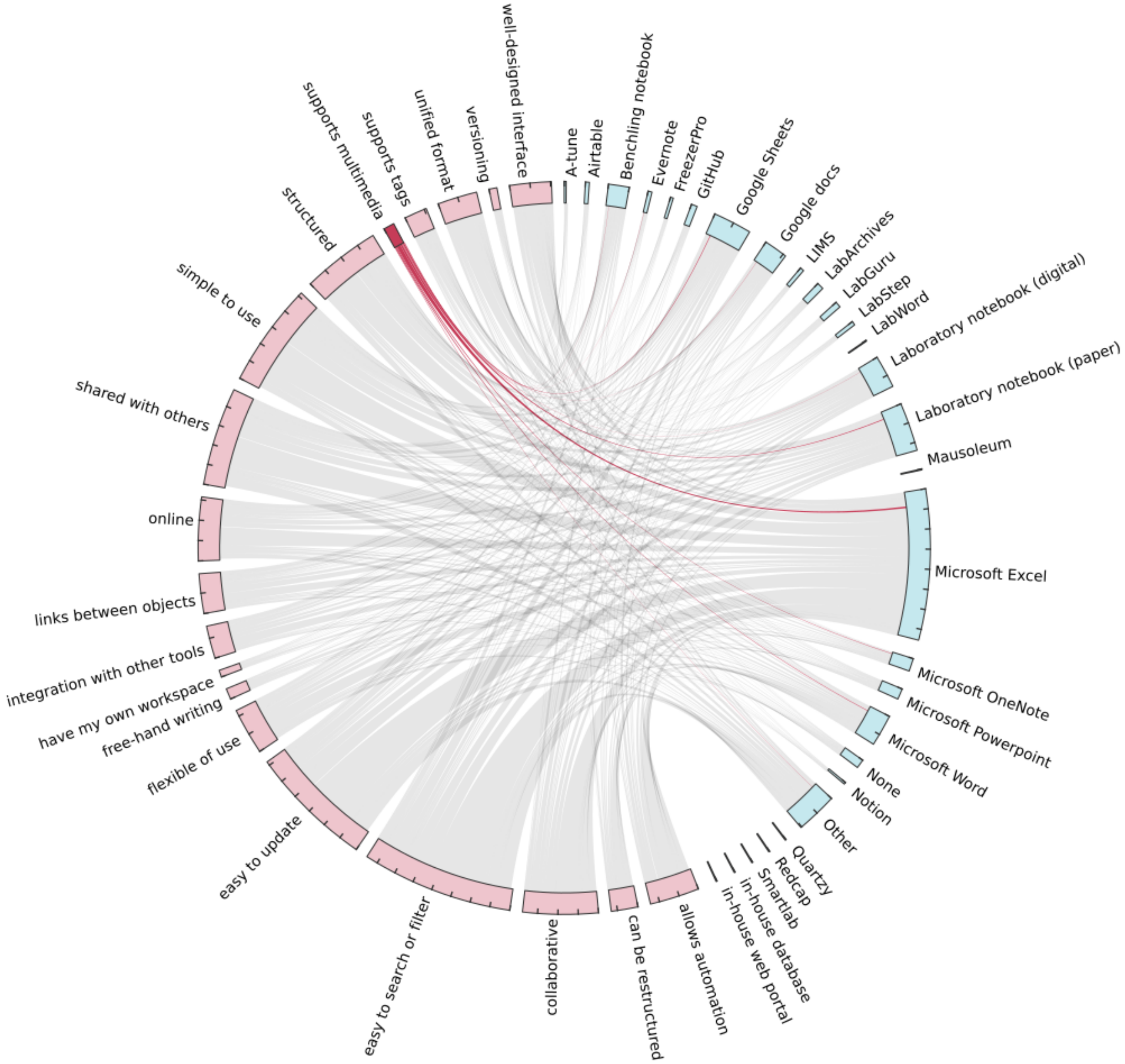


Sample-management solutions (blue arcs) vs desired features (red arcs). Arc length depicts the number of respondents, with ticks denoting 20 respondents. Link thickness shows the fraction of respondents using a solution and valuing a criterion. The chords connected to the arc 'supports multimedia' are highlighted in color.'

supports tags

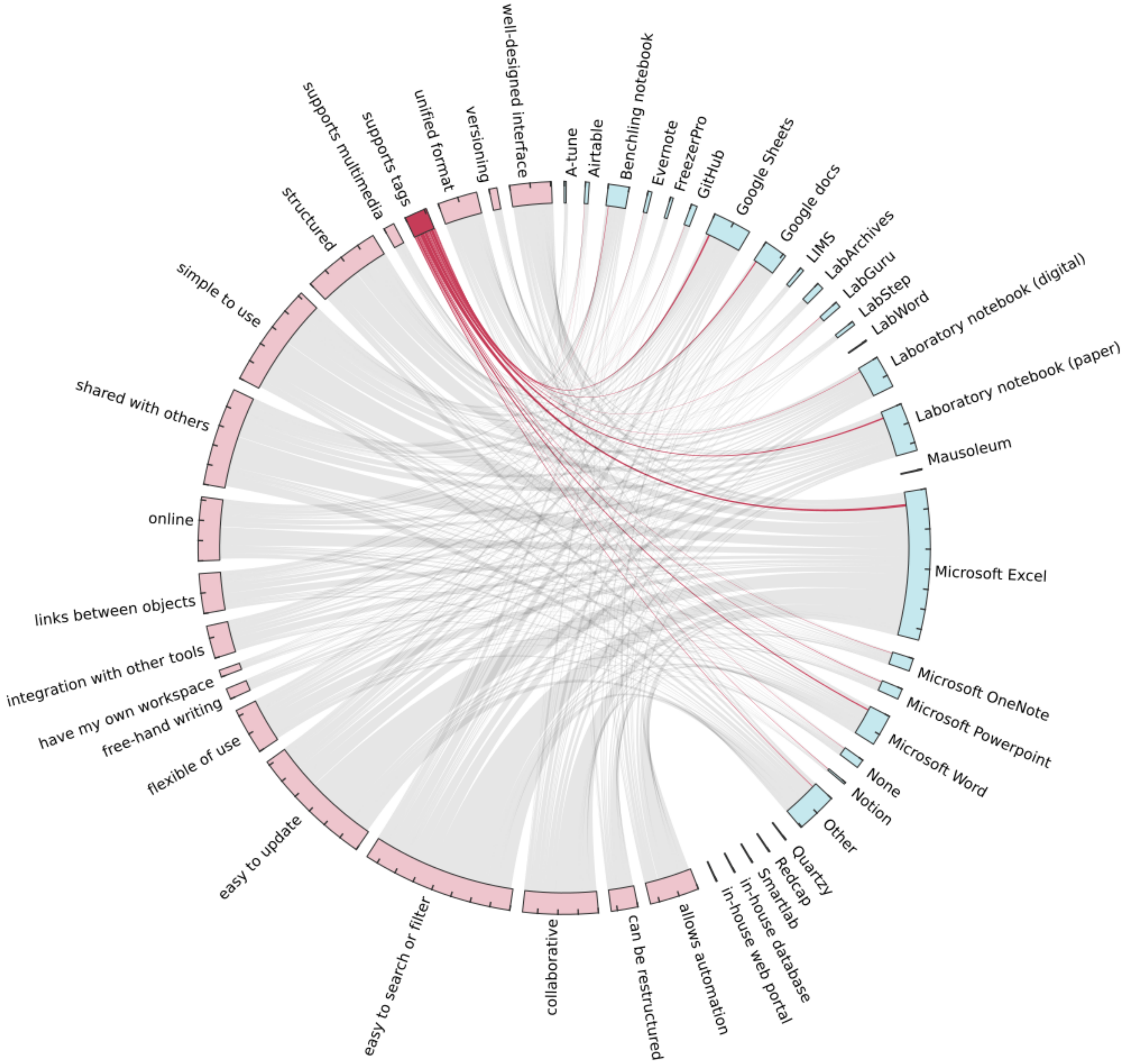


Sample-management solutions (blue arcs) vs desired features (red arcs). Arc length depicts the number of respondents, with ticks denoting 20 respondents. Link thickness shows the fraction of respondents using a solution and valuing a criterion. The chords connected to the arc 'supports tags' are highlighted in color.'

unified format

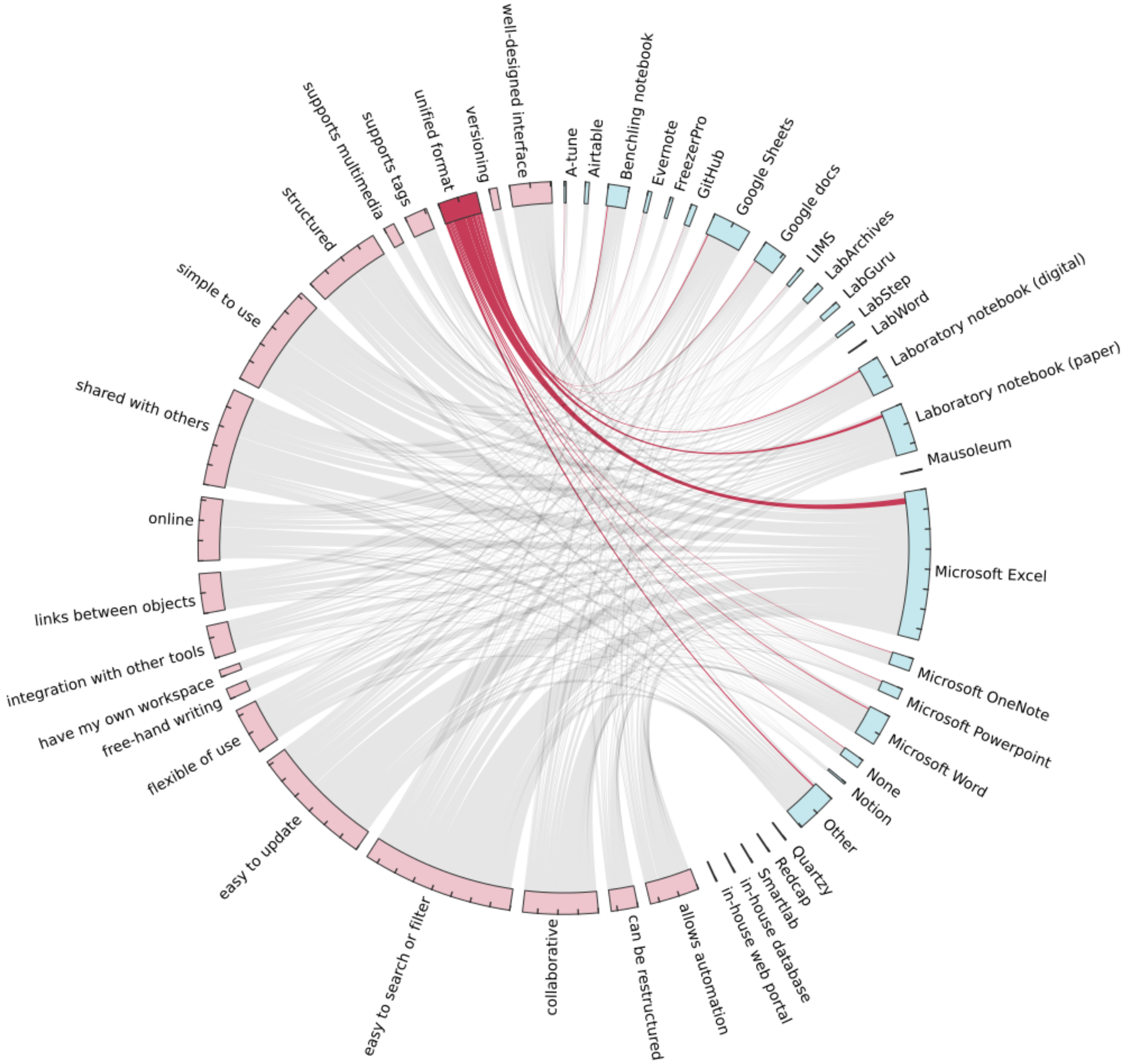


Sample-management solutions (blue arcs) vs desired features (red arcs). Arc length depicts the number of respondents, with ticks denoting 20 respondents. Link thickness shows the fraction of respondents using a solution and valuing a criterion. The chords connected to the arc 'unified format' are highlighted in color.'

versioning

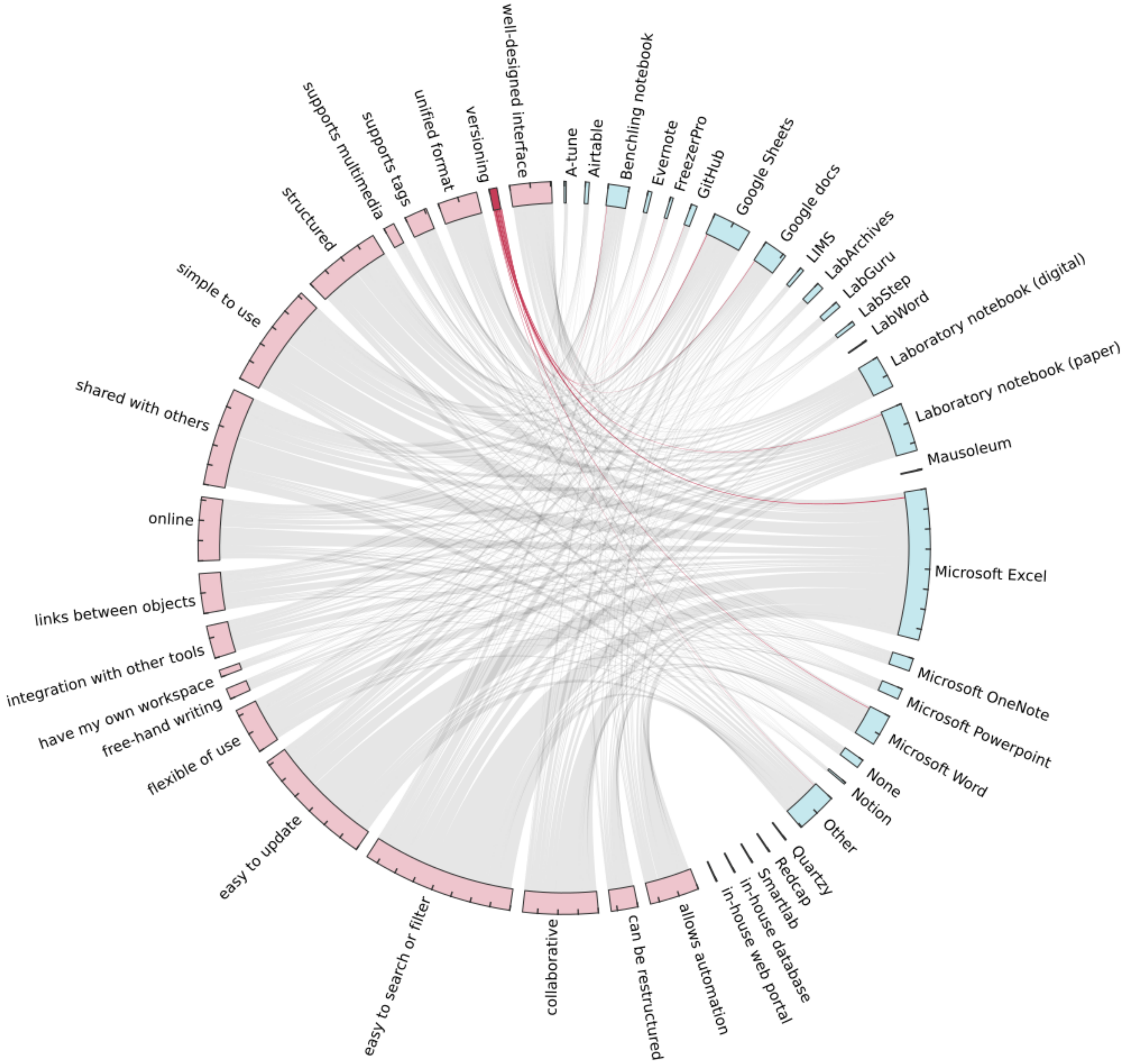


Sample-management solutions (blue arcs) vs desired features (red arcs). Arc length depicts the number of respondents, with ticks denoting 20 respondents. Link thickness shows the fraction of respondents using a solution and valuing a criterion. The chords connected to the arc 'versioning' are highlighted in color.'

well-designed interface

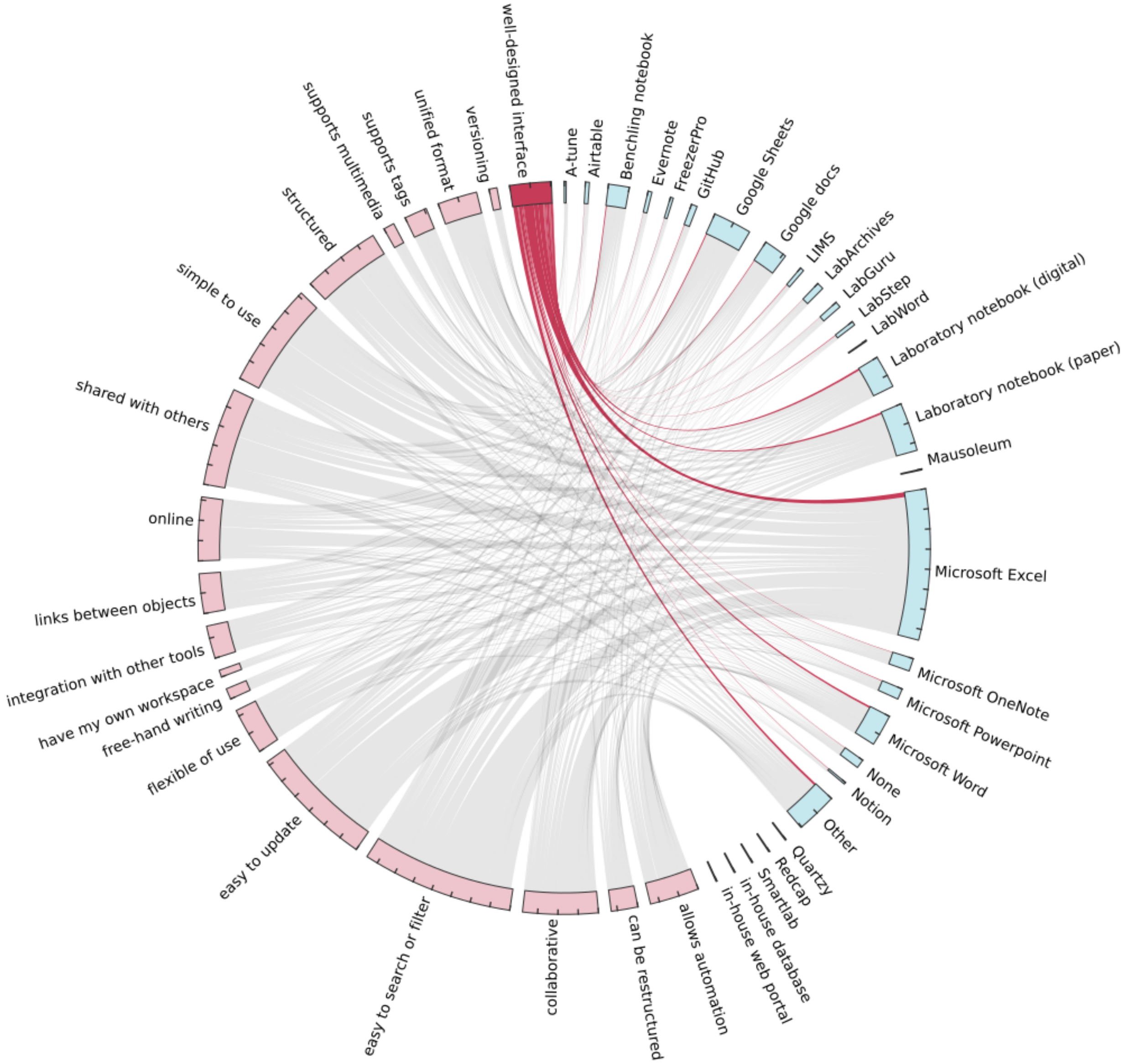


Sample-management solutions (blue arcs) vs desired features (red arcs). Arc length depicts the number of respondents, with ticks denoting 20 respondents. Link thickness shows the fraction of respondents using a solution and valuing a criterion. The chords connected to the arc 'well-designed interface' are highlighted in color.'
