## Supplementary Data 1 for "A platform for lab management, note-keeping and automation": Supplementary Data 1 - Survey Form.pdf

### How do you register samples and notes?

~60 sec to complete

Your position

Research subject

A few keywords that describe your field of work

Please provide the name of the PI of your lab or your company name *

This information is needed to account for multiple responses from the same lab

What note-taking solution do you use? *

- [ ] Airtable
- [ ] Benchling notebook
- [ ] Evernote
- [ ] Github
- [ ] Google docs
- [ ] Google Sheets
- [ ] LabArchives
- [ ] LabGuru

- [ ] Laboratory notebook (digital)
- [ ] Laboratory notebook (paper)
- [ ] LabStep
- [ ] Microsoft Excel
- [ ] Microsoft OneNote
- [ ] Microsoft Powerpoint
- [ ] Microsoft Word
- [ ] Notion
- [ ] None
- [ ] Other
- [ ] ElabJournal
- [ ] LabWord
- [ ] Signals
- [ ] Microsoft Sharepoint
- [ ] TextEdit
- [ ] Confluence
- [ ] Labfolder

If other, please specify:

Does everyone use the same solution in your lab?

- ( ) Yes
- ( ) No

Are notes shared with other lab members?

- ( ) Yes
- ( ) No

Anything to add on notes?

How do you keep track of samples? *

Cells, mice, DNA samples, etc

- [ ] Airtable
- [ ] Benchling notebook
- [ ] Evernote
- [ ] GitHub
- [ ] Google docs
- [ ] Google Sheets
- [ ] LabArchives
- [ ] LabGuru
- [ ] Laboratory notebook (digital)
- [ ] Laboratory notebook (paper)
- [ ] LabStep
- [ ] Microsoft Excel
- [ ] Microsoft OneNote
- [ ] Microsoft Powerpoint
- [ ] Microsoft Word
- [ ] Notion
- [ ] None
- [ ] Other
- [ ] ElabJournal
- [ ] in-house sample tracking system
- [ ] in-house web portal
- [ ] in-house database
- [ ] LabWord
- [ ] File Maker
- [ ] Smartlab
- [ ] A-tune
- [ ] FreezerPro
- [ ] Quartzy
- [ ] ChemInv
- [ ] LIMS
- [ ] Mausoleum
- [ ] Redcap

If other, please specify:

Does everyone use the same system in your lab?

- ◯ Yes
- ◯ No

Are sample registries shared between lab members?

- ◯ Yes, we all use the same system
- ◯ No, we organise our samples separately

Are sample registries standardised?

- ◯ Yes, I use template-based description
- ◯ No, I use free-form description

How do you label samples in the lab?

Samples include tubes, plates, flasks etc

- ◯ I use the full name, genotype etc
- ◯ I simplify the name, genotype etc
- ◯ I use numbers and keep a list of what is what
- ◯ I use the same number as in the registry
- ◯ Other

Anything to add about sample management?

Do you have time to answer two more questions?

- ◯ Sure
- ◯ Mm 😒

Clear Selection

#### Note-taking: what's most important?

- [ ] allows automation
- [ ] can be restructured
- [ ] collaborative
- [ ] easy to search or filter
- [ ] easy to update
- [ ] flexible of use
- [ ] free-hand writing
- [ ] have my own workspace
- [ ] integration with other tools
- [ ] links between objects
- [ ] online
- [ ] shared with others
- [ ] simple to use
- [ ] structured
- [ ] supports multimedia
- [ ] supports tags
- [ ] unified format
- [ ] versioning
- [ ] well-designed interface

#### Sample management: what's most important?

- [ ] allows automation
- [ ] can be restructured
- [ ] collaborative
- [ ] easy to search or filter
- [ ] easy to update
- [ ] flexible of use
- [ ] free-hand writing
- [ ] have my own workspace
- [ ] integration with other tools
- [ ] links between objects
- [ ] online
- [ ] shared with others
- [ ] simple to use
- [ ] structured
- [ ] supports multimedia
- [ ] supports tags

- [ ] unified format
- [ ] versioning
- [ ] well-designed interface

#### Any more comments?

#### email

Submit

Do not submit passwords through this form. Report malicious form
